# C-terminal domain mobility underpins substrate engagement by the *Chaetomium thermophilum* EDEM:PDI ERAD-L checkpoint complex

**DOI:** 10.1101/2025.01.29.635535

**Authors:** Charlie J. Hitchman, Andrea Lia, Mohammed Bhogadia, Geander M. Pessoa, Gabriela N. Chiriţoiu, Cristian V. A. Munteanu, Juan R. Ortigosa, Jorge O. Lannot, Simona Ghenea, Ikuo Wada, Maria De Benedictis, Gabor Tax, Yusupha Bayo, Irene Crescioli, Laura Sanna, Daniele Di Bella, James Birch, Michela Bollati, Dominic L. Alonzi, Andrew Quigley, Carlos P. Modenutti, Ştefana M. Petrescu, Nobuko Hosokawa, Angelo Santino, Bibek Gooptu, Pietro Roversi

## Abstract

The ERAD-L checkpoint complex de-mannosylates misfolded glycoproteins with lumenal defects, targeting them to retrotranslocation, ubiquitination, and proteasomal degradation. Commitment to ERAD-L requires an Endoplasmic Reticulum-Degradation Enhancing ***α***-Mannosidase-like protein (EDEM) and its associated Protein Disulfide Isomerase (PDI). We determined Cryo-EM structures of the *Chaetomium thermophilum* EDEM:PDI heterodimer, both by itself and in complex with a classic ERAD substrate, the ***α***1-antitrypsin Null Hong Kong mutant (A1AT-NHK). The EDEM catalytic domain nestles within the PDI arc. One intermolecular disulfide, between the a′ domain of PDI and the first conserved cysteine of the EDEM linker (Cys A), stably links the two proteins. A second intermolecular disulfide, between the second conserved cysteine of the EDEM linker (Cys B) and the PDI a domain, stabilises the compact apo conformation. In the substrate-bound complex, the Cys B intermolecular disulfide is reduced and the catalytic CXXC motif of the PDI a domain oxidised. Release of the Cys B linkage increases the conformational freedom of the EDEM linker, allowing the EDEM C-terminal domains to adopt both proximal and distal conformations relative to the catalytic domain. The structures provide a framework for discovery of EDEM:PDI modulators, with potential applications in virology, rare genetic disease and cancer, and as reagents for glycoprotein quality-control engineering.

## Introduction

Proteins entering the eukaryotic secretory pathway must fold correctly before they can perform their biological functions. To ensure proteome integrity, the endoplasmic reticulum (ER) operates a sophisticated protein quality-control system that promotes productive folding while identifying proteins that fail to attain a native conformation. Terminally misfolded glycoproteins are ultimately eliminated through Endoplasmic Reticulum-Associated Degradation (ERAD) [1–4], targeting them for retrotranslocation to the cytosol, ubiquitination, and proteasomal degradation. ERAD contributes to proteostasis of healthy cells [5]; in the background of a pathogenic mutation in a gene encoding a secretory protein, ERAD relieves ER stress and alleviates acute proteostatic challenges, although it can underpin pathogenic loss-of-function phenotypes [6–9]. When secretory protein misfolding within the ER overwhelms ER degradation capacity, the ER stress response is triggered [10]. Persistently misfolded secretory proteins not processed by ERAD are prone to toxic aggregation [11, 12]. Prolonged ER stress is a major mediator of chronic disease [13, 14].

Recognition of ERAD glycoprotein clients bearing defects in their ER-luminal domains (ERAD-L) [15] is effected by an Endoplasmic Reticulum-Degradation Enhancing 1,2-*α*-Mannosidase-like protein (EDEM) [16]. Fungal EDEMs and EDEM1s/EDEM3s flag a terminally misfolded glycoprotein for degradation by selective demannosylation of one of its *N*-linked glycans [17–20]. In particular, once the outer mannose from the C branch of an *N*-glycan on the client is removed (see Supplementary Figure S1), the demannosylated glycan exposes a *α*-D-Man-(1→6)-*α*-D-Man moiety, which in turn is recruited to either the OS-9 or XTP3-B lectin [20, 21] associated with the ERAD retrotranslocon [20, 22]. Through the latter, the misfolded glycoprotein is exported to the cytoplasm, where it is ubiquitinated and eventually digested by a proteasome [23–26].

In fungi, only one EDEM gene is present [27, 28] whereas plants and animals encode more than one paralogue. All EDEMs contain a Glycosyl Hydrolase 47 (GH47) catalytic domain (Pfam family PF01532, Glyco hydro 47 (GH47) [29]). In fungal EDEMs and in EDEM3s the GH47 domain is followed by a linker and one or more C-terminal domains [30, 31]. In ERAD checkpoint complexes [4, 32] an EDEM is disulfide bonded to a Protein Disulfide Isomerase (PDI) family member [33]. The first of these EDEM:PDI heterodimers to be discovered was the *Saccharomyces cerevisiae* (*Sc*) *Sc*Mnl1/Htm1:*Sc*Pdi1 heterodimer [17, 34–36] whose Cryo-EM structure was recently determined [31]. EDEM1 forms a heterodimer with P4HB (*aka* PDIA1) [33]^1^; EDEM3 forms a heterodimer with ERp46 [33, 42]; EDEM2^2^ associates with TXNDC11 [19, 33, 40, 45].

The exact function of the PDI in the ERAD misfolding checkpoint complex has been the subject of some debate in the field. PDI redox chemistry plays at least one certain role in the complex: it covalently tethers EDEM and PDI. All known EDEM:PDI complexes are linked by (at least) one disulfide bridge [17, 19, 31, 33–36, 43, 44]. In the recent *Sc*Mnl1:*Sc*Pdi1 Cryo-EM structure (PDB ID 8ZPW), *Sc*Mnl1 Cys579 is disulfide bonded to Cys406 of the ^406^CXXC^409^ redox motif of the fourth thioredoxin domain (TRX4) of *Sc*Pdi1 [31, 34]. All known EDEM sequences share this Cys residue (“Cys A”, see Supplementary Figure S2). In addition, fungal EDEMs and mammalian EDEM3s have a second unpaired cysteine residue (“Cys B”), located downstream from Cys A - in a stretch of sequence following GH47 domain - and preceding their C-terminal domains. In the *Sc*Mnl1:*Sc*Pdi1 Cryo-EM structure, this Cys B (*Sc*Mnl1 Cys644) is disulfide bonded to Cys61 of the ^61^CXXC^64^ motif of the TRX1 domain of *Sc*Pdi1 [31, 34]. The exact role of the redox chemistry at the EDEM Cys B site remains unclear.

We provide here the evolutionary, structural and functional bridge connecting the fungal EDEM checkpoint to mammalian EDEMs. We report Cryo-EM structures of the EDEM:PDI heterodimer from the thermophilic filamentous fungus *Chaetomium thermophilum* (*Ct*), alone and in complex with a classic ERAD substrate, the Null Hong Kong variant of *α*1-antitrypsin (A1AT-NHK, [16, 46]). We show that EDEM:PDI intermolecular disulfide redox chemistry, as well as mediating complex formation, is essential for EDEM folding, substrate recruitment and enzymatic efficiency.

## Results

### Phylogenetic analysis places *Ct*EDEM within the fungal lineage co-orthologous to mammalian EDEM1 and EDEM3

The evolutionary relationships among fungal, plant and animal EDEMs have been examined by treating EDEMs as members of the GH47 family and consequently emphasizing the evolution of their catalytic domains [28, 32, 47, 48]. We reconstructed what to our knowledge is the first comprehensive phylogeny of the EDEM family, specifically addressing the ancestry of the EDEM1/3 paralogues, and their evolutionary relationship with EDEM2s, using the newly revealed structural/domain relationships and the sequences of their conserved GH47 catalytic domain, including the stretch of sequence immediately following it (see Extended Data section 18). The resulting tree (Supplementary Fig. SS3) contains a basal clade of fungal EDEMs and is consistent with diversification of the plant and animal EDEM paralogues through successive duplication events. EDEM1 and EDEM3 cluster most closely with the fungal EDEM lineage, whereas EDEM2 occupies a more divergent branch. Notably, the analysis confirms the *Arabidopsis thaliana* MNS5 and MNS4 as the plant orthologues of EDEM3 and EDEM2 [48], respectively, providing phylogenetically supported assignment of plant EDEMs. These findings identify fungal EDEMs as representatives of an ancestral ERAD checkpoint lineage and provide an evolutionary context for the structural and functional characterization of *Ct*EDEM:*Ct*PDI presented below.

### *Ct*EDEM functions as a conserved ERAD checkpoint mannosidase across kingdoms

Structural biology on eukaryotic proteins can profit from the enhanced protein stability of *Chaetomium thermophilum (Ct)* [49]. A search in the *Ct* genome using the protein sequence of the *Saccharomyces cerevisiae* EDEM, *Sc*Mnl1, yielded UniProt entry https://www.uniprot.org/uniprot/G0SCX7 (hereinafter *Ct*EDEM), annotated as *α*-1,2-mannosidase. Although the orthologous budding yeast protein is commonly termed Mnl1 (or Htm1), we use the designation *Ct*EDEM because the fungal protein belongs to the single-copy fungal EDEM lineage and our phylogenetic analysis places this lineage close to mammalian EDEMs. A search in the same genome using the protein sequence of *Sc*PDI1 yielded UniProt entry https://www.uniprot.org/uniprot/G0SGS2 (hereinafter *Ct*PDI), annotated as a protein disulphide isomerase.

We first set out to test *Ct*EDEM ERAD mannosidase activity *in vivo* using an available plant model [50, 51]. The *Arabidopsis thaliana bri1-5* mutant encodes a misfolded brassinosteroid receptor that is retained in the ER and degraded in an MNS4/MNS5-dependent manner, resulting in a characteristic dwarf phenotype. The expression of *Ct*EDEM gene in the *At mns4-1 mns5-1 bri1-5* triple mutant background restores the *bri1-5* dwarf phenotype, re-endowing the plant with ERAD mannosidase activity (Supplementary Figure S4). *Ct*EDEM is therefore active in recognising the misfolded receptor *bri1-5*, removing mannose residues from the glycans on *bri1-5* and dispatching the receptor to ERAD degradation. *Ct*EDEM can therefore functionally engage the plant ERAD machinery. In plants, the luminal factor(s) that assist MNS4/MNS5 in folding-status recognition remain unknown [52]. In our experiments, *Ct*EDEM may therefore either function without a dedicated PDI partner in the plant ER lumen or be assisted by a PDI orthologous to *Sc*PDI1^3^.

To test whether *Ct*EDEM ERAD activity extends to human cells, two distinct canonical ERAD substrates were assayed in HEK293T EDEM3 KO cells: A1AT-NHK and soluble Tyrosinase (ST-TYR); the cells were transfected with *Ct*EDEM and/or *Ct*PDI expressing plasmids. A1AT-NHK has one free cysteine residue (Cys 256) which forms a disulfide bond with human ERAD checkpoint complexes [54]. The same A1AT-NHK Cys residue mediates A1AT-NHK disulfide bond dimerisation and accumulation of covalent A1AT-NHK dimers is selectively prevented by the over-expression of EDEM1 [55]. Only over-expression of *Ct*PDI alters the A1AT-NHK dimer/monomer, with more monomers being detected, see Figure 1A.

**Figure 1.**
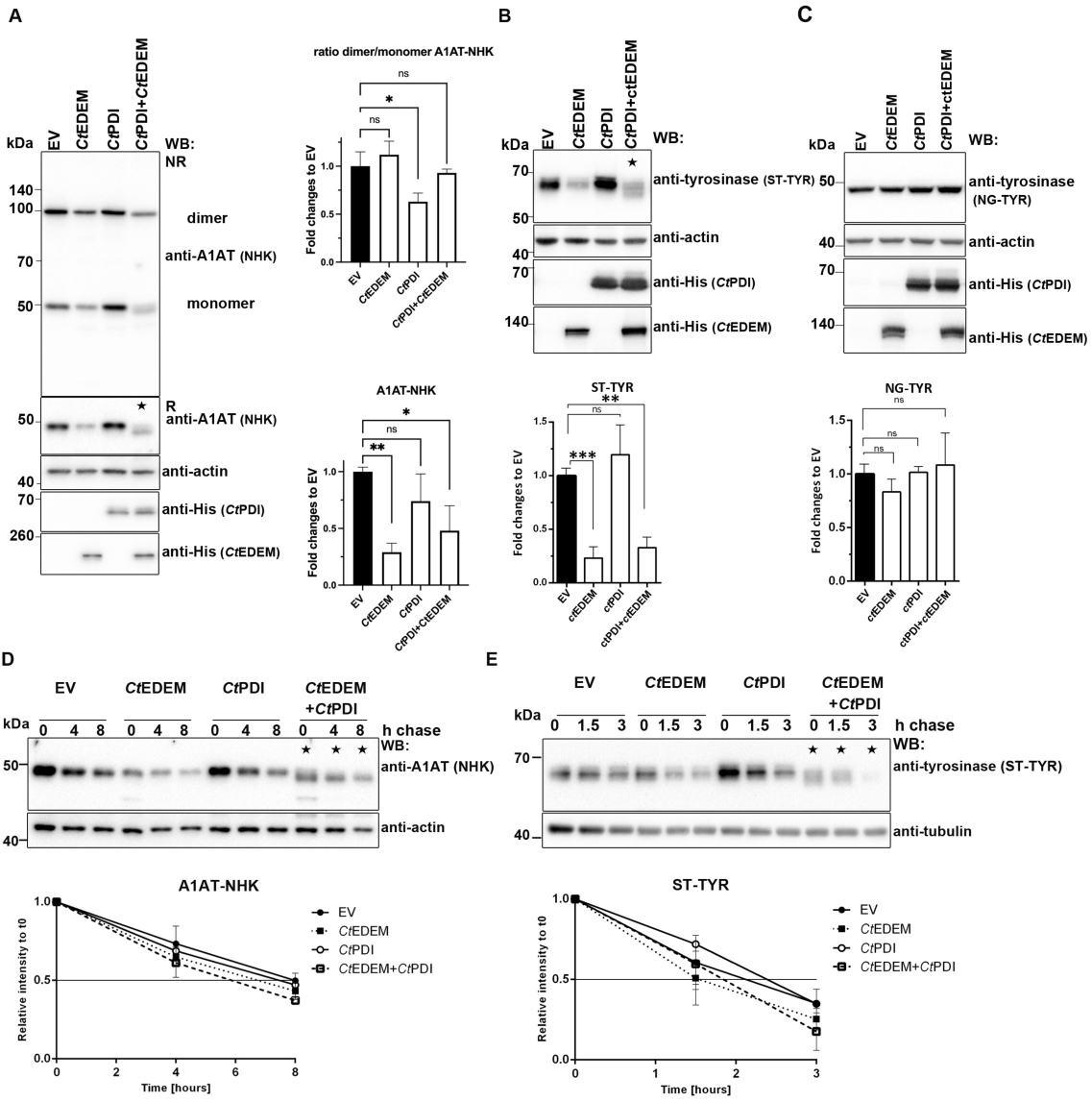
*Ct*EDEM has ERAD mannosidase activity in human cells. **A:** Left panel: Western Blot showing the effect of *Ct*EDEM and *Ct*PDI on A1AT-NHK in HEK293T EDEM3-KO cells co-transfected with A1AT-NHK and either empty vector (EV), *Ct*EDEM, *Ct*PDI, or both. Equal amounts of protein were resolved under non-reducing (NR) or reducing (R) conditions transferred to nitrocellulose and probed with the following antibodies: rabbit anti-A1AT, mouse anti-His (*Ct*PDI, *Ct*EDEM) and mouse anti-actin (loading control). Right upper panel: Quantification of the A1AT-NHK dimer/monomer ratios from Image J densitometry, expressed as fold change relative to EV (mean of n=3, ±SD). Right lower panel: Quantification of A1AT-NHK band intensity under reducing conditions, expressed as fold change relative to EV (mean of n=3, ±SD). One-way ANOVA with Dunnett’s test was used for statistical analysis; ns=not significant; *: p *<*0.05 and **: p *<*0.01. **B:** Upper panel: Western Blot showing *Ct*EDEM and *Ct*PDI effects on soluble tyrosinase (ST-TYR). Samples were prepared as in (**A**) under reducing conditions and probed with the indicated antibodies. Lower panel: Quantification of ST-TYR band intensity, expressed as fold change relative to EV (mean of n=3, ±SD). One-way ANOVA with Dunnett’s test, ***: p *<*0.001. **C:** Upper panel: Western Blot showing *Ct*EDEM and *Ct*PDI effects on non-glycosylated tyrosinase (NG-TYR). Samples were prepared as in (**A**) under reducing conditions and probed with the indicated antibodies. Lower panel: Quantification of NG-TYR band intensity, expressed as fold change relative to EV (mean of n=3, ±SD). Statistics as in (**A**). **D–E:** Cycloheximide chase assays assessing the half-life of A1AT-NHK (D) and ST-TYR (**E**). HEK293T EDEM3-KO cells were co-transfected with the ERAD substrate (A1AT-NHK or ST-TYR) and either EV, *Ct*EDEM, *Ct*PDI, or both, then treated with cycloheximide for the indicated times. Lysates were resolved under reducing conditions and probed with anti-A1AT (**D**) or anti-tyrosinase (**E**), with anti-actin (**D**) or anti-tubulin (**E**) as loading controls. Lower panels: Quantification of substrate levels relative to t0, based on densitometry of blots in the corresponding upper panels. A star indicates de-mannosylated species.

In cells transfected with *Ct*EDEM (both alone and with *Ct*PDI), degradation of A1AT-NHK and ST-TYR is enhanced, see Figure 1, confirming that *Ct*EDEM has ERAD mannosidase activity also in human cells against these misfolded substrates.

### Cryo-EM structures of *Ct*EDEM:PDI

Having confirmed ERAD activity of *Ct*EDEM, we cloned mature *Ct*EDEM and *Ct*PDI constructs, lacking predicted membrane-anchoring or ER-retrieval sequences and carrying C-terminal His-tags for purification, in the pHLsec vector [58] for secreted expression in mammalian cells (Supplementary Figure S5). The *Ct*EDEM:PDI heterodimer was expressed and purified, alone and in complex with A1AT-NHK (Supplementary Figures S6, S7 and S8), and the apo *Ct*EDEM:PDI and *Ct*EDEM:PDI:A1AT-NHK Cryo-EM structures were determined to an overall resolution of 2.6 Å and 3.5 Å respectively, see Figure 2 and Supplementary Figures S9, S10, S11 and S12. The refined models and Cryo-EM maps of the *Ct*EDEM:PDI apo and A1AT-NHK bound structures were deposited in the Protein Databank (PDB) and in the Electron Microscopy Data Bank (EMDB), with deposition IDs 9TPA and EMD-56105 (apo structure)^4^ and PDB ID 29TE, Extended PDB ID pdb 000029TE, EMD ID EMD-57361 (A1AT complex), respectively.

**Figure 2.**
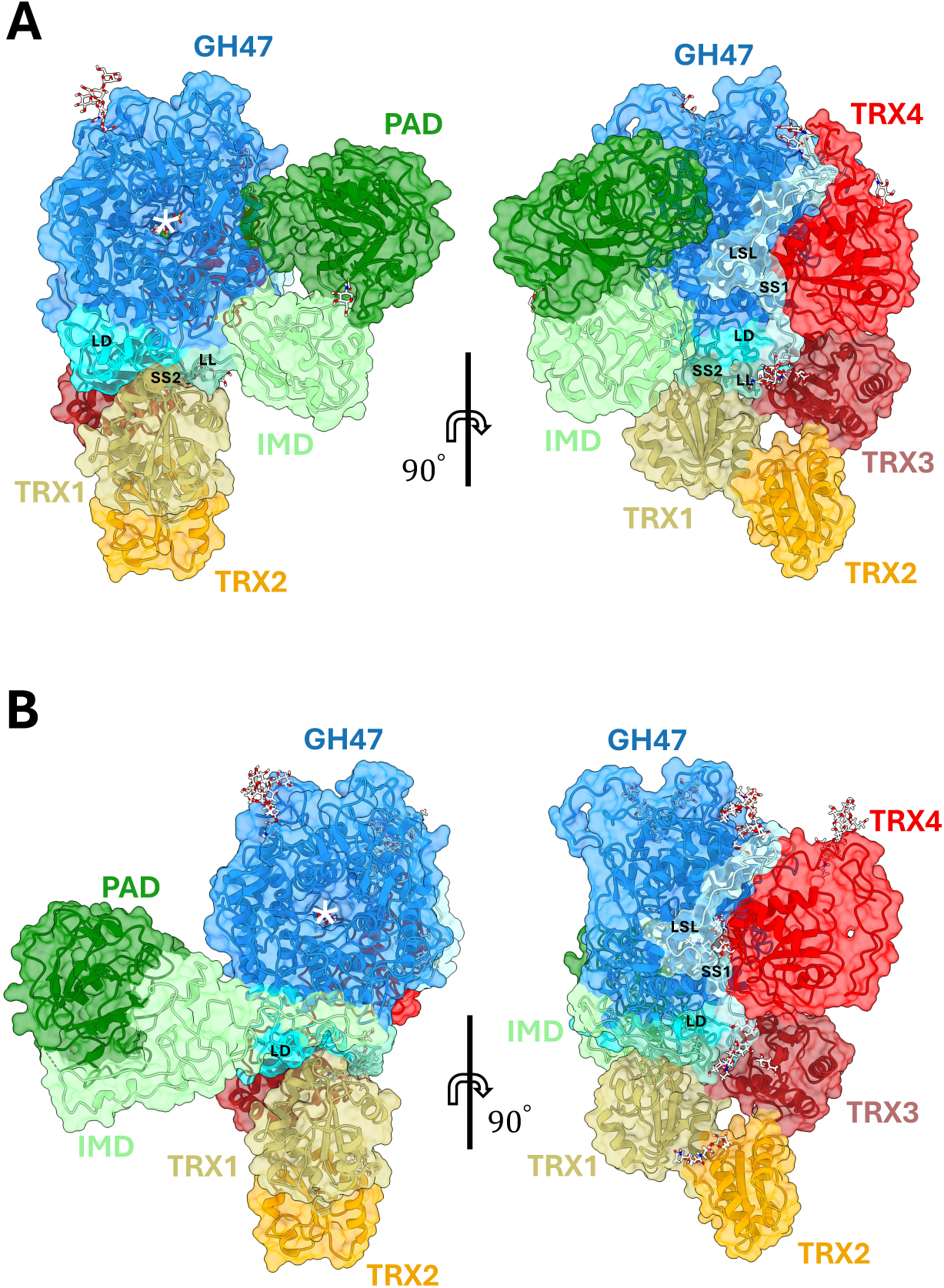
*Ct*EDEM:PDI Cryo-EM structures. Left- and right-hand side views are related by a rotation of 90*^◦^* around the vertical axis. *Ct*EDEM: GH47 catalytic domain (64-615, blue); L-shaped linker, linker domain and linear linker (LSL, 616-655, pale cyan; LD, 656-713, cyan; and LL, 714-725, green cyan); Intermediate Domain (IMD 726-821 and 1069-1092, pale green); Protease Associated Domain (PAD 822-1067, green). *Ct*PDI: TRX1 (18-128, dark yellow); TRX2 (129-228, orange); TRX3 (229-342, brick red); TRX4 (343-484, red). The inter-chain *Ct*EDEM C647:*Ct*PDI C385 disulfide bond is labelled SS1. White asterisk: GH47 active site. *N*-linked glycans are in stick representation, coloured by atomic element (C: white; N: blue; O: red). The *Ct*EDEM N-terminal sequence 1-63 [56] was removed by cloning, and so was the ER retrieval C-terminal HDEL motif of *Ct*PDI. **A**: Apo *Ct*EDEM:PDI. The second inter-chain disulfide bond (*Ct*EDEM C719:*Ct*PDI C50) is labelled SS2; a 56-residue loop of the PAD (845-900) and the C-termini (*Ct*EDEM 1084-1093 and *Ct*PDI 484-515) could not be traced. **B**: *Ct*EDEM:PDI in complex with A1AT-NHK. A 73-residue stretch of *Ct*EDEM PAD sequence (834-906) could not be traced. Figure was made with ChimeraX [57].

In both the apo heterodimer and the A1AT-NHK-bound complex, the GH47 domain nestles within the curved arc formed by PDI (Fig. 2; Supplementary Fig. S13A), as observed in the recently determined *Sc*Mnl1:*Sc*PDI1 heterodimer structure (PDB ID 8ZPW) [31]. The first three TRX domains of *Ct*PDI adopt the three-foil/cloverleaf structure typical of the N-terminal portion of most PDIs (yellow, orange and brick-red domains in Figure 2 and Supplementary Figure S13A), with TRX2 jutting out and not contacting *Ct*EDEM at all. The observed *Ct*EDEM:PDI interface rationalises the loss of function of mutants of *Ct*EDEM orthologues, see Supplementary Figure S14.

Due to the flexibility of the IMD-PAD region in both the apo- and more extensively in the substrate-bound states, the local resolutions vary between these and the core domains. The core domains for the apo-state show a local resolution range of 2.5-3.3 Å, and for the substrate bound complex of 3.0-5.0 Å (see Supplementary Figures S9 and S10) and superpose closely with an RMSD_C_ of 1.1 Å (over 661 C*_α_*s). The IMD-PAD local resolution ranges are 4.0-5.4 Å for the apo-state, and 6.0-10.0 Å for the substrate-bound state, see Supplementary Figures S9 and S10. The resolution of the A1AT-NHK portion in the complex is too low to attempt building an atomic model for the client and the deposited model only comprises the *Ct*EDEM:PDI heterodimer in the A1AT-NHK bound conformation, see Figure 2B.

***Ct*EDEM GH47 domain** The catalytic domain of the mannosidase (*Ct*EDEM residues 64-615) has the classic GH47 (*α/α*)7-barrel fold, with the active site at the base of the barrel [60]. The brim of the barrel is decorated by 7 *β*-hairpins; its bottom is plugged by a *β*-hairpin at the C-terminus (*Ct*EDEM residues 602-615), see Supplementary Figures S13B and S15A [29, 61, 62]. The *Ct*EDEM GH47 domain has a very similar structure to the one of *Sc*Mnl1in the *Sc*Mnl1:*Sc*PDI1 structure (PDB ID 8ZPW [31]): RMSD*_C_* =1.0 Å over 429 residues, see Supplementary Figure S15A). No major differences are visible in the active site between the apo and the A1AT complex structures, except that in the apo Cryo-EM map there is density for a ligand bound in the active site – which was built as *D*-Mannose (see Supplementary Figure S13C). The *Ct*EDEM conserved acidic residues in the active site are E191, D370, E522, while the conserved Thr residue coordinating the catalytic Ca^2+^ ion with its main chain O and its side chain O*_γ_*_1_ is T608 (structurally and functionally equivalent to *Hs*EDEM1 E225, D370, E493 and T579) [63]. The conserved *Ct*EDEM Cys99-Cys558 intramolecular disulfide bond (see Supplementary Figure S2) clips the first *β* hairpin of the GH47 7-barrel to the last one, and in so doing it zips shut the barrel. Table S1 summarises the cysteine residues discussed throughout the manuscript, using *Ct*EDEM:*Ct*PDI as the structural reference system and mapping conserved cysteine classes to the corresponding residues in *Sc*Mnl1:Pdi1 and selected mammalian EDEM:PDI complexes.

***Ct*EDEM linker regions** The GH47 domain is followed by *Ct*EDEM residues 616-725 which mediate most of the contacts with the *Ct*PDI. This linker region counts three sub-regions:

1. **L-shaped linker:** the first stretch of this portion (*Ct*EDEM residues 616-651, coloured pale cyan in Figure 2) is an L-shaped linker which starts after the GH47 C-terminal hairpin, sits over *Ct*EDEM helices *α*9*/α*10 and *α*13*/α*14 and wraps over the surface of the *Ct*PDI-TRX4 domain, eventually running parallel to the *Ct*PDI-TRX3/4 linker. The L-shaped linker contains the conserved *Ct*EDEM Cys647 (commonly referred to as “Cys A”, see Supplementary Figure S2 [19, 34, 54]). *Ct*EDEM Cys647 forms an inter-molecular disulfide bond with *Ct*PDI Cys385 in the ^385^CXXC^388^ redox motif of the *Ct*PDI-TRX4 domain^5^. Such SS bond is labelled “SS1” in Figure 2 and it is shown in detail in Supplementary Figure S16B. This disulfide bond is necessary for *Ct*EDEM, as well as for other EDEMs, to function correctly [19, 42, 45]. The L-shaped linker has similar size and shape in *Ct*EDEM (both apo and A1AT-NHK complex) and in the *Sc*Mnl1:*Sc*PDI1 structure (PDB ID 8ZPW [31]), where the ordered portion of the L-shaped linker (*Sc*Mnl1 503-588) contains the intermolecular disulfide bridge between *Sc*Mnl1 Cys579 and *Sc*PDI1 Cys406 (Supplementary Figure S15B). It is worth noting that the EDEM-PDI intermolecular disulfide bond can form by isomerism (starting with the free Cys on the EDEM and the oxidised disulfide bridge on the PDI) and its establishment does not require external redox equivalents.
2. **Helical linker sub-domain:** following the L-shaped linker, a helical linker sub-domain (*Ct*EDEM residues 656-713, cyan in Figures 2) nestles between helices *α*14*/α*15 of the GH47 domain and the TRX1 domain of the PDI. The *Ct*EDEM helical linker sub-domain is similar in size and shape to the one in the *Sc*Mnl1 structure (residues 594-634 in PDB ID 8ZPW [31], Supplementary Figure S15B). Dali and Foldseek searches [65, 66] do not detect structural homology between this helical linker sub-domain and any structure in the PDB. The helical linker sub-domain is an EDEM point of contact with the PDI: the exposed *Ct*EDEM Leu656 forms an intermolecular hydrophobic contact with the *Ct*PDI surface pocket formed by Pro235, Tyr238, Tyr250, Tyr287 and His290, equivalent to the hydrophobic contact surface established between *Sc*Mnl1 Phe591, Trp592 and Tyr593 and the *Sc*PDI1 TRX3 residues Gly246, Phe249, Tyr261, Phe298 and His301 [31], see Supplementary Figure S15C.
3. **Linear linker:** following the helical linker sub-domain, a short linear linker (*Ct*EDEM residues 714-725, green cyan in Figure 2 and red sticks in Supplementary Figure S15B) contains *Ct*EDEM Cys719, conserved across fungal EDEMs and usually referred to as Cys B [19, 34, 54]. The linear linkers (and the C-terminal domains that follow it) differ in the *Ct*EDEM:PDI apo heterodimer and its A1AT-NHK bound complex, see Supplementary Figure S16A:

a. in the apo heterodimer, *Ct*EDEM Cys719 forms an inter-molecular SS bond with *Ct*PDI Cys50 in the ^50^CGHC^53^ redox motif of *Ct*PDI-TRX1 domain, while *Ct*PDI Cys53 is a free thiol. This disulfide bond is labelled “SS2” in Figure 2A (a close-up view is in Supplementary Figure S16B);
b. in the A1AT-NHK bound complex, *Ct*EDEM Cys719 is free and the *Ct*PDI-TRX1 Cys50-Cys53 disulfide bridge is oxidised; a close-up view is in Supplementary Figure S16C. The extra conformational freedom accompanying the reduction of this intermolecular SS bond in the A1AT-NHK bound heterodimer allows the linear linker to detach itself from the GH47 domains and the C-terminal domains move away from the EDEM catalytic domain, see Figures 2B and 3.

**Figure 3.**
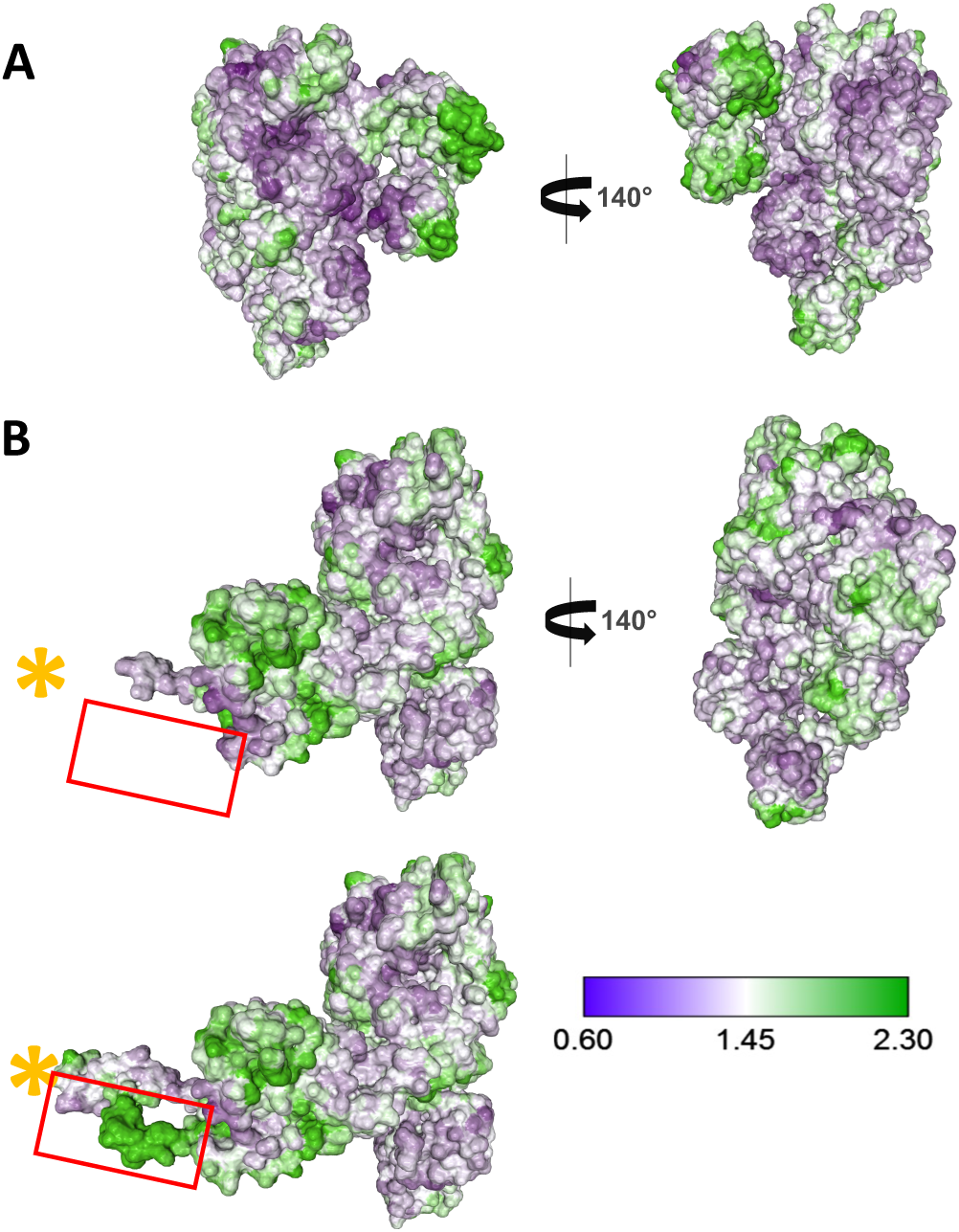
*Ct*EDEM:PDI surface hydrophobic patches localise to the flexible IMD-PAD domains. Two views (related by a rotation of 140° around the vertical axis) of the *Ct*EDEM:PDI heterodimer model, in surface representation and coloured by surface hydrophobicity, calculated as the ratio of non-polar to polar residues [67]. Hydrophilic regions are painted pale violet and hydrophobic regions are painted green, as indicated by the hydrophobicity scale bar. Hydrophobic patches localise to the flexible IMD-PAD domains. (**A**): apo form and (**B**): A1AT-NHK bound. The upper **B** panel omits the untraceable PAD region (residues 834–906; red rectangle); the bottom **B** panel shows in the same rectangle one possible conformation for the same region, in order to highlight the hydrophobicity of the stretch. The yellow asterisk (∗) marks a point inside the volume occupied by the A1AT-NHK substrate in the map.

***Ct*EDEM C-terminal domains** In fungal EDEMs and in mammalian EDEM3s, the linkers are followed by one [31] or two [30] C-terminal domains - which in EDEM3 were termed Intermediate Domain (IMD) and Protease-Associated Domain (PAD) [30]. The EDEM3 C-terminal domains have been implicated as mediators of recruitment of misfolded glycoprotein substrates in that they modulate the ERAD turnover of the mutant clients A1AT-NHK and ST-TYR [30]. A mutant of *Sc*Mnl1 lacking the C-terminal domain has only low mannosidase activity against its substrate and it is inactive in ERAD; the same *Sc*Mnl1 recombinant C-terminal domain preferentially bound completely unfolded polypeptides [31].

In the *Ct*EDEM:PDI structures, the PAD domain is inserted in the fold of the IMD, and the two *Ct*EDEM C-terminal domains are topologically intertwined: the PAD (residues 822-1067) is encoded by a stretch of sequence inserted in the two sequences encoding the IMD (726-821 and 1068-1092), with the last 24 residues of the protein forming a two-stranded *β* hairpin that completes the *β* sandwich fold of the IMD. AlphaFold correctly predicted this topological feature but did not predict correctly the relative orientation of the IMD-PAD tandem with respect to the GH47 domain [59].

As described above, the distinct redox states of the Cys719-containing linear linker in the apo and A1AT-NHK bound complexes are accompanied by large-scale changes in the relative disposition of the C-terminal *Ct*EDEM domains:

- in the *Ct*EDEM apo structure, the IMD-PAD domains hang on the brim of the EDEM GH47 catalytic pocket (see Figure 2A). The main IMD:GH47 contact is formed by the *Ct*EDEM hairpin residues 778-782, nestling in a shallow surface pocket of the GH47 domain (*Ct*EDEM residues 437-460). Overall, the *Ct*EDEM IMD domain in the apo structure is similar in shape and structure to the *Sc*Mnl1 C-terminal domain: RMSD*_C_* =2.2 Å over 94 residues. In particular, the stretch of *Ct*EDEM IMD residues immediately following the linear linker (725-745) forms the outer strand of the central three-stranded *β* sheet of the IMD (see Supplementary Figure S16B), equivalent to residues 650-665 in the IMD of *Sc*Mnl1. The rest of the *Ct*EDEM IMD (746-821) in the apo structure resembles the same portion of the complex with A1AT-NHK: RMSD*_C_* =1.4 Å over 76 residues. In the apo structure, the PAD loop 845-900 (contiguous with the density where the A1AT-NHK client is in the ternary complex) is disordered;
- in the complex of *Ct*EDEM with A1AT-NHK, the inter-molecular disulfide bridge between *Ct*EDEM Cys719 and *Ct*PDI Cys50 is reduced, and the portion immediately following the linear linker (*Ct*EDEM 726-745) lies across the surface of the GH47 domain – pointing to the opposite face of the complex, so that they no longer belong to the central *β* sheet of the IMD. Rather, they form an extension of the linear linker, resting on a different surface of the GH47 domain, and bringing the C-terminal domains away from the *Ct*EDEM catalytic site (Figure 2B and Supplementary Figure S16C).

As in the apo *Ct*EDEM:*Ct*PDI heterodimer, part of the PAD remains untraceable in the A1AT-NHK-bound complex; however, the disordered region expands from residues 845–900 in the apo structure to residues 834–906 in the client-bound structure. However, owing to the rearrangement of the IMD and PAD domains in the latter structure, the two hinge regions flanking the untraceable PAD region become contiguous with the density assigned to the A1AT-NHK client (Figure 3 and Supplementary Figures S10A and D), suggesting that residues within this flexible region may directly contact the substrate. The face of the *Ct*EDEM IMD-PAD arm surrounding this loop (*Ct*EDEM IMD residues 728-732, 745-749, 792-794; PAD residues 832-833, 907-912, 939-947) is identified as a non-polar surface (green in Figure 3) using the protein-sol server (https://protein-sol.manchester.ac.uk/) [67]. The C-terminal region of the fungal ERAD mannosidase *Sc*Mnl1 also contains a hydrophobic substrate-recognition groove required for degradation of the model ERAD substrate CPY and for binding partially folded proteins [31]. The proximity of the A1AT-NHK density to the hydrophobic *Ct*EDEM IMD-PAD surface in our structure provides structural support for a conserved role of EDEM C-terminal domains in client recognition.

The large rearrangement of the EDEM linker and C-terminal domains between the apo and the A1AT-NHK bound states is accompanied by a minor re-orientation of the TRX1-3 domains of the PDI - which 3D variability analysis detects as a flexible region in both the apo and ternary complex set of particles (see Supplementary Figures S19, S20 and Supplementary Videos 1-4).

### A conserved architecture at the *Ct*EDEM active-site entrance underlies C-branch specificity

Structurally homologous GH47 domains are present in the three ERAD mannosidases (EDEM1 [16, 18, 68–70], EDEM2 [71], and EDEM3 [46]), as well as in ER *α*1,2-mannosidase I (ERManI/MAN1B1, [72–75]), in the Golgi *α*1,2-mannosidases (GMIB/MAN1A2 and GMIC/MAN1C1 [76]) and in GMIA/MAN1A1, which has been reported to localise to quality-control vesicles and to participate in glycoprotein targeting to ERAD [77]. Structures of ERManI and GMIA have been determined in the absence and presence of inhibitors, substrate analogues and substrate glycans [61, 62, 73, 74, 76]. In contrast to the highly conserved −1 and +1 subsites, the remainder of the substrate-binding groove and the surface loops decorating the *Ct*EDEM GH47 barrel differ substantially from those observed in these enzymes. These structural elements are thought to determine selectivity for the specific branch of the high-mannose glycan engaged by the active site [73, 76]. Consistent with this specialization, ERManI and EDEM2 remove the terminal *α*1,2-linked mannose from branch B, whereas EDEM1, EDEM3 and the single EDEM orthologues found in fungi trim branch C of M8B to produce M7BC (GlcNAc_2_Man_7_*_BC_*).

In particular, a conserved region (*Ct*EDEM, “I” in Supplementary Figure S21) spans residues 100–118 and it is present in ∼800 EDEM1, EDEM3 and fungal EDEM orthologues. In these proteins, region “I” includes an *α*-helix (*Ct*EDEM helix *α*2, residues 111–118 (see Figures 4A,C and Supplementary Figure S2), which immediately follows the conserved intramolecular disulfide bond that stabilises the GH47 catalytic domain by linking the first and last *β*-hairpins of the (*α/α*)_7_ barrel (Supplementary Figure S2) [19, 34, 42]. *In silico* docking of the M8B glycan substrate to the *Ct*EDEM active site reveals that this *α*2-helix contributes directly to substrate binding. Together with Trp516, helix *α*2 defines a composite binding surface for the M8B substrate. At the entrance of the catalytic pocket, helix *α*2 constrains the trajectory of the glycan such that the second GlcNAc is positioned against Trp516 and effectively sandwiched between its indole side chain and Glu113, see Figures 4A,C.

**Figure 4.**
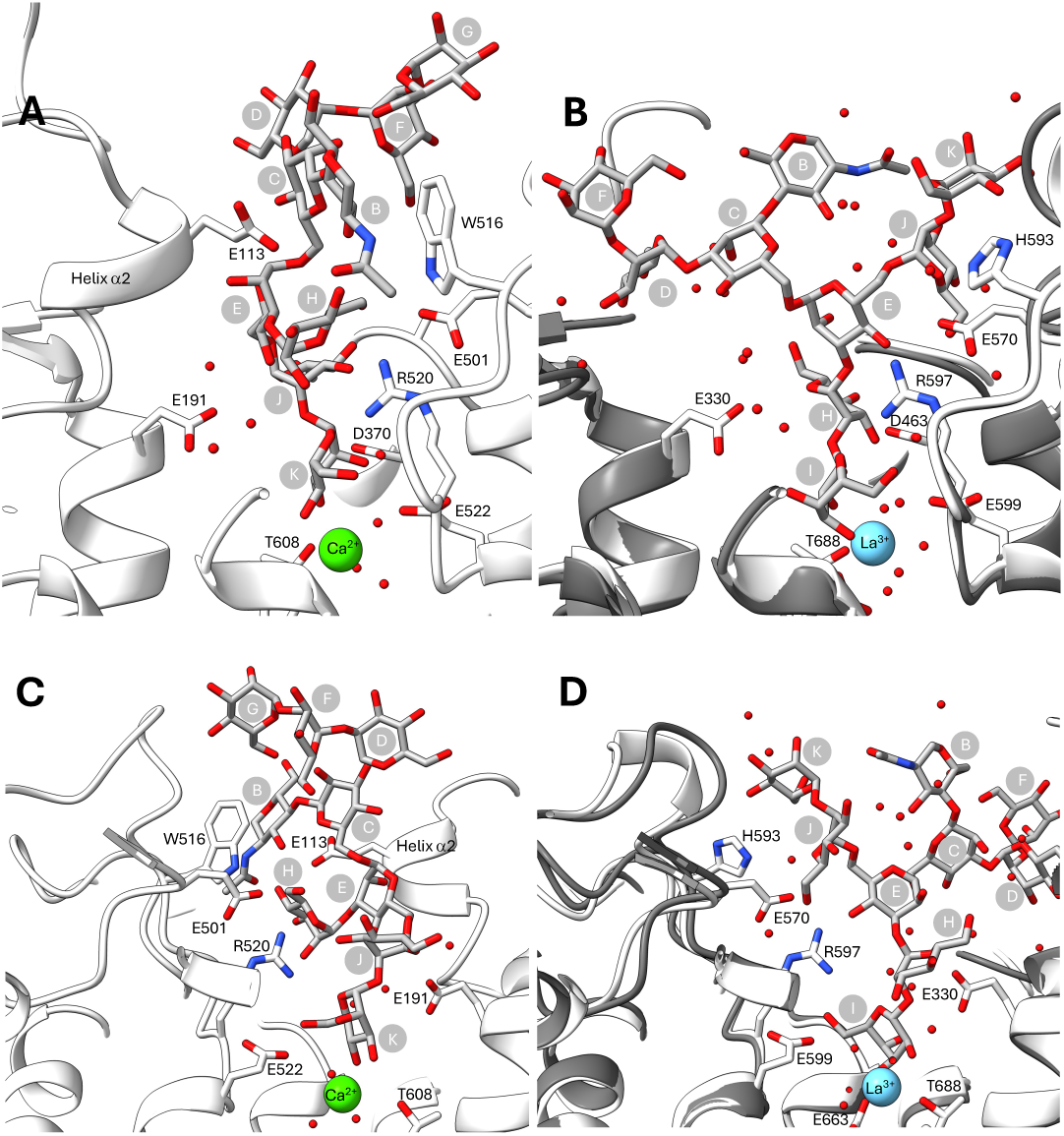
Comparison of *N*-glycan binding by the *Ct*EDEM and human ERManI active sites. **A,C:** Two views of the model of GlcNAc_1_Man_8_*_B_ N*-glycan (sticks with grey C atoms) docked to the *Ct*EDEM GH47 active site. The GlcNAc residue (“B”) at the entrance of the catalytic pocket is sandwiched between W516 and helix *α*2 (E113, see Supplementary Figure S2). The same residues W516 and E113 also form the walls of a shallow pocket in which the Mannose “H” of the cleaved branch B of the M8B glycan sits; the floor of this pocket is formed by the conserved salt bridge R520:E501. Branch C of the glycan occupies the catalytic site (Man “J” and “K”). The *Ct*EDEM catalytic acidic residues E191, D370, E522, and the T608 residue coordinating the catalytic Ca^2+^ ion (green sphere) are in sticks representation. **B,D:** Equivalent views to the ones in A,C of the crystal structure of the GlcNAc_1_Man_8_*_A_ N*-glycan (sticks with grey C atoms) bound to human ERManI (PDB IDs 5KIJ, [76]). The ERManI structure (white C atoms) is overlaid on an AlphaFold3 model of human EDEM2 (dark grey, cartoon only). The human ERManI catalytic acidic residues E390, D463, E599 [74] and the T688 residue coordinating the La^3+^ ion (cyan sphere) substituting the Ca^2+^ are in sticks representation. Helix *α*2 is absent from these structures and at the position of *Ct*EDEM Trp516 ERManI His593 is found. In the human EDEM2 model, Trp516 is replaced by Gly366. The region of space occupied by helix *α*2 in *Ct*EDEM is filled by the GlcNAc_1_Man_8_*_A_ N*-glycan, specifically by the first branching mannose (“C”) and the proximal mannose of the A branch (“D” and “F”). The salt bridge R597:E570 orients Man “E” and “J” - leading branch B of the glycan (Man “H” and “I”) into the catalytic site.

Further into the binding groove, the same *α*2-helix and Trp516 contribute to formation of the pocket accommodating the B-branch mannose (Man “H”; for glycan-residue labelling, see Figure S1), which bears the outer mannose cleaved by either ERManI or EDEM2 before the *N*-glycan can enter the GH47 active site, see Figures 4A,C. In the *Ct*EDEM model, Glu113 forms a contact with the O4 hydroxyl of Man “H”, while Trp516 engages the C6–O6 region of the same sugar. Thus, helix *α*2 and Trp516 jointly define both the orientation of the glycan at the entrance of the active site and the positioning of the cleaved branch B within the binding groove. The *α*2-helix is conserved in EDEM1/EDEM3/fungal EDEM orthologues, while the position corresponding to Trp516 contains either Trp or Gln/Glu (see Supplementary Figure S2). Conversely, GH47 *α*1,2-mannosidases that do not recognise branch C - including human EDEM2, human ERManI and yeast ERManI - lack the characteristic architecture of the *Ct*EDEM glycan-entry tunnel: in these enzymes, helix *α*2 is absent, allowing the glycan to occupy the space filled by this helix in *Ct*EDEM, and Trp516 is replaced by smaller residues, see Figures 4B,D.

Taken together, these observations indicate that C-branch specificity in fungal EDEMs, EDEM1s and EDEM3s, depends on a coupled structural module centred on helix *α*2 and a bulky side chain corresponding to *Ct*EDEM Trp516, which together define the trajectory of the glycan at the entrance of the active site: they engage the middle GlcNAc (“B”) and the cleaved B-branch mannose (Man “H”), and position the glycan in a conformation conducive to C-branch processing.

### Redox chemistry in the *Ct*EDEM:PDI complex

The foldase activity of PDIs depends upon their CXXC redox motifs (*e.g.* CGHC in *Ct*PDI). The first cysteine in these motifs can either form an intra-molecular disulfide bond with the second Cys of the motif (“resolving Cys”) or it can form an inter-molecular disulfide bond to a Cys residue in a client protein: the competition between these alternatives allows binding and release of the clients to catalyse the reshuffling of client disulfide bonding patterns [78, 79]. For example, *Hs*ERp46 (the partner of *Hs*EDEM3) forms mixed disulfide bonds with client glycoproteins while acting as a refolding enzyme, catalysing the reduction of non-native disulfides [80].

Whether similar mixed disulfides are formed with ERAD clients, or contribute to their recruitment, has not been investigated, although strong formation of mixed disulphides between *Hs*ERp46 and client glycoproteins containing Cys residues was observed when the cells were treated with castanospermine, a broad-spectrum mannosidase inhibitor that blocks entrance and exit of folding glycoproteins into/from the calnexin cycle, suggesting a EDEM client-recruiting role for ERp46 in the context of the EDEM3:ERp46 ERAD misfolding checkpoint complex [80]. Association of EDEM1 with the misfolded A1AT-NHK was shown to be thiol-dependent and to require a single Cys residue in the substrate, but not EDEM1 thiols [70], suggesting the possibility of mixed disulfides between the PDI and glycoprotein substrates. Indeed, the free Cys 256 of A1AT-NHK is functionally important during ERAD commitment: EDEM1 suppresses the accumulation of disulfide-linked NHK dimers [55]. These observations prompt the question whether EDEM-associated oxidoreductases may form mixed disulfides with ERAD glycoprotein clients.

In order to test the hypothesis that *Ct*PDI may recruit a glycoprotein client to *Ct*EDEM via mixed disulfide bonding, HEK293F cells were co-transfected with *Ct*EDEM, A1AT-NHK and wild-type *Ct*PDI. Mutation of the resolving cysteine in a PDI CXXC motif stabilises mixed disulfide intermediates with substrates [80, 81]. For example, an *Hs*ERp46 mutant lacking all resolving Cys residues in its TRX domains (“CXXA” mutant) trapped human EDEM3 in high-molecular weight complexes, resulting in the disappearance of EDEM3 monomer and dimer species [42]. As well as the wild-type *Ct*PDI, we also tested the *Ct*PDI Cys53Ser (Cys B trapping mutant).

*Ct*EDEM:PDI heterodimers and *Ct*EDEM:PDI:A1AT-NHK ternary complexes were purified from the media by Immobilised Metal Affinity Chromatography (IMAC) followed by Size Exclusion Chromatography (SEC), see Supplementary Figures S7 and S8. Under non-reducing conditions, the anti-His Western blot of the SEC fraction corresponding to the heterodimer peak detects His-tagged *Ct*PDI and/or *Ct*EDEM-containing species (Figure 5A, left-hand side lane). In the wild-type sample, the main His-positive band is compatible with monomeric *Ct*PDI, whereas the *Ct*PDI Cys53Ser trapping mutant shifts the signal towards higher-molecular-weight species, including bands compatible with disulfide-linked *Ct*EDEM:PDI complexes (Figure 5A, middle lane). The increased accumulation of disulfide-linked *Ct*EDEM:PDI species in the trapping-mutant sample is also consistent with the *Ct*EDEM Cys719-SS-*Ct*PDI Cys50 intermolecular disulfide observed in the apo Cryo-EM structure, and suggests reversible redox chemistry at the EDEM Cys B–PDI Cys50 disulfide bond.

**Figure 5.**
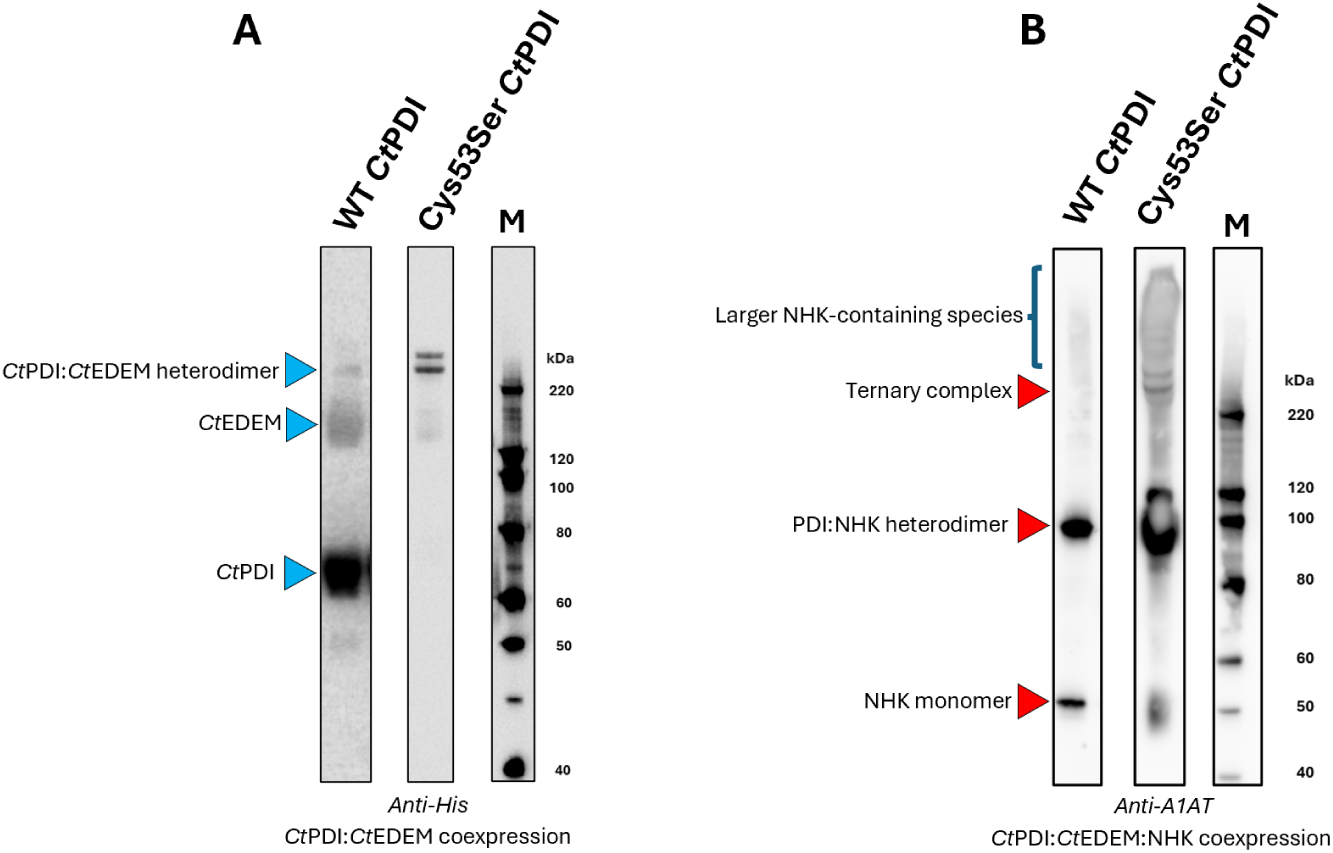
The Cys53Ser *Ct*PDI trapping mutant favours disulfide bond formation with *Ct*EDEM and A1AT-NHK. HEK293F cells were co-transfected with *Ct*EDEM A1AT-NHK and either wild-type *Ct*PDI or *Ct*PDI Cys53Ser trapping mutant. *Ct*EDEM:*Ct*PDI heterodimers or their ternary complexes with A1AT-NHK were purified from the media by IMAC followed by SEC. Selected fractions from both SEC experiments were loaded in non-reducing conditions in 4-12% polyacrylamide gels, transferred on nitrocellulose membrane and probed with indicated antibodies: rabbit anti-A1AT, mouse anti-His for detection of *Ct*PDI and *Ct*EDEM. **A**: Anti-His Western blot from *Ct*EDEM:PDI (WT and mutant) heterodimer co-purification with a SEC fraction corresponding to the heterodimer peak. **B**: Anti-A1AT Western blot from ternary complex co-purification with a SEC fraction corresponding to the ternary complex peak. M indicates the lane loaded with molecular weight markers.

The anti-A1AT Western blot of the SEC fraction corresponding to the ternary-complex peak reports only A1AT-NHK-containing species. Compared with the wild-type sample (Figure 5B, left-hand side lane), the *Ct*PDI Cys53Ser trapping mu-tant produces a ladder of high-molecular-weight A1AT-NHK-containing species, while the monomeric A1AT-NHK band weakens (Figure 5B, middle lane). A band migrating at approximately 100 kDa is consistent with a A1AT-NHK dimer. Together, these data indicate that the Cys53Ser trapping mutant stabilises disulfide-linked *Ct*PDI:A1AT-NHK and larger *Ct*EDEM:PDI:A1AT-NHK-containing complexes, supporting the idea of a role for *Ct*PDI redox chemistry during client engagement.

### Mammalian EDEM3:ERp46 function depends on intra- and inter-molecular redox chemistry

To gain insight into the role of redox chemistry in the *Mm*EDEM3:*Hs*ERp46 heterodimer, we generated *Mm*EDEM3 and *Hs*ERp46 Cys mutants. The single mutants *Mm*EDEM3 C529S and C558S abrogate the SS1 and SS2 intermolecular disulfide, respectively; the double mutant C529S/C558S cannot form SS1 nor SS2. In human *Hs*ERp46 we generated the triple trapping mutant CXXA, *aka* “CA”-*Hs*ERp46C92A/C220A/C353A - altering the resolving Cys of each of its TRX domains CXXC redox motifs: *Hs*ERp46 ^89^CGHC^92^, ^217^CGHC^220^ and ^350^CGHC^353^.

To probe the oxidation state of the EDEM, single and double Cys-to-Ser mutants that impair EDEM intramolecular disulfide bridge formation were also studied: *Mm*EDEM3 C83S, C442S and C83S/C442S. These mutants cannot form the EDEM GH47 domain intramolecular disulfide bond [42] so that only the reduced EDEM3 band is visible in SDS-PAGE (lower band of the *Mm*EDEM3 doublet around 120 kDa in lanes 5-10 in Figure 6A).

**Figure 6.**
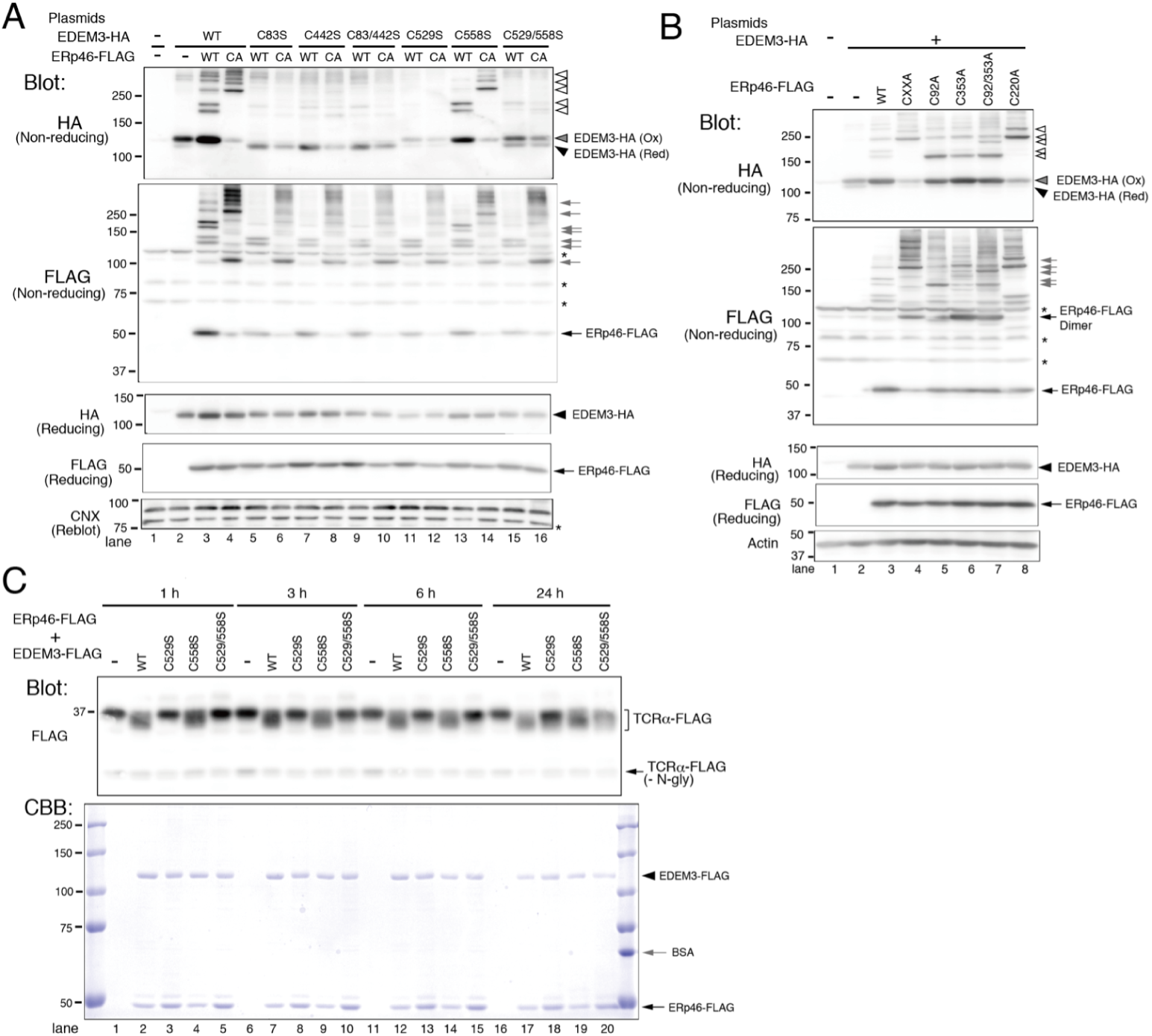
Mammalian EDEM3:ERp46 disulfide bonds and activity assay. HEK293T were transfected with *Mm*EDEM3-HA and *Hs*ERp46-FLAG and cell lysates were separated by SDS-PAGE under reducing or non-reducing conditions followed by anti-HA and anti-FLAG Western blotting. Black filled triangles indicate reduced *Mm*EDEM3-HA, gray filled triangles indicate oxidised *Mm*EDEM3-HA and open white triangles indicate disulfide-bonded high molecular mass complexes containing *Mm*EDEM3-HA. Black arrows indicate monomeric *Hs*ERp46-FLAG, and gray arrows denote disulfide-bonded high-molecular-mass complexes containing *Hs*ERp46-FLAG. Asterisks indicate non-specific antibody signals. **A:** *Mm*EDEM3 WT or a Cys-to-Ser mutants were co-expressed with WT *Hs*ERp46-FLAG or the triple trapping mutant CXXA (*Hs*ERp46 C92A/C220A/C353A, *aka* “CA”). **B:** WT *Mm*EDEM3-HA was co-expressed with the indicated *Hs*ERp46-FLAG single, double or triple trapping mutant. **C:** *In vitro* mannosidase activity of the purified *Mm*EDEM3:*Hs*ERp46 complex was assayed using TCR*α*-FLAG as a substrate. Activity was monitored by the mobility shift of the ∼37 kDa TCR*α*-FLAG band following mannose trimming. The non-glycosylated TCR*α*-FLAG species is labelled (-*N*-gly). The lower panel shows Coomassie staining of the upper gel region, with BSA used as a loading reference.

**Cys A in the EDEM3 linker region forms a stable disulfide to ERp46 and plays a role in EDEM3 oxidation** We studied *Mm*EDEM3:*Hs*ERp46 disulfide-bonded heterodimer formation by analysing the EDEM3 linker Cys mutants on non-reducing SDS-PAGE (lanes 11-16 in Figure 6A). In presence of WT ERp46, the strongest band for the oxidised *Mm*EDEM3:*Hs*ERp46 disulfide-bonded heterodimer was formed by *Mm*EDEM3 WT and its C558S mutant (no Cys B in linker), see Figure 6A, upper band of the doublet around 200 kDa in lanes 3 and 13. The EDEM3 mutants in which Cys A is absent (C529S and C529S/C558S) give weaker bands for the oxidised heterodimer (Figure 6A, upper band of the doublet around 200 kDa in lanes 11,12 and 15,16). The *Mm*EDEM3 C529S (no Cys A in linker) and *Mm*EDEM3 C529S/C558S (no free Cys in linker) mutants cause an increase in the amounts of the reduced form of EDEM3 (lower band of the doublet around 120 kDa in Figure 6A, lanes 11, 12, 15 and 16). Together, these observations suggest that intermolecular disulfide redox chemistry following the establishment of an intermolecular disulfide between a Cys residue in ERp46 and EDEM3 Cys A (*Mm*EDEM3 C529) enhances EDEM3 oxidation.

In an assay of mannose-trimming activity, using TCR*α*-FLAG as a substrate, no mannosidase activity was detected for the *Mm*EDEM3 mutants lacking Cys A (EDEM3 C529S and C529S/C558S) up to 6 h of incubation (Figure 6C, lanes 3, 5, 8, 10, 13, and 15). Only after 24 h of incubation mild mannose-trimming activity was observed (Figure 6C, lanes 18 and 20), consistent with equivalent observations for the Cys A mutant of *Sc*Mnl1 in the *Sc* heterodimer [31]. These results suggest that EDEM3 can acquire catalytic competence either through slow ERp46-independent oxidation or through weak, non-covalent association with ERp46 that partially compensates for the absence of the Cys A disulfide bond.

**The *Hs*ERp46^217^CXXC^220^ motif is required for establishment of the inter-molecular disulfide to EDEM3 Cys A** Interestingly, the *Mm*EDEM3:*Hs*ERp46 heterodimer doublet band is absent from the samples co-transfected with *Mm*EDEM3 WT (or C558S) and the ERp46 CXXA triple trapping mutant (Figure 6A, lanes 4 and 14), even if these EDEM3 mutants have an intact Cys A: enhanced laddering is also observed at molecular mass ≥ 250 kDa. To probe which of the three trapping mutants most affects the formation of the disulfide bridge with *Mm*EDEM3 Cys A, we studied the *Hs*ERp46 C92A, C220A and C353A single trapping mutants, as well as the double trapping mutant C92A/C353A.

All the *Hs*ERp46 trapping mutants that do not interfere with the ^217^CGHC^220^ motif formed the *Mm*EDEM3:HsERp46 disulfide-bonded heterodimer (Figure 6B, lanes 5-7). By contrast, the C220A mutation targets the resolving cysteine of the same redox motif that provides Cys217 for SS1 and causes the usual heterodimer band around 150 kDa to disappear (Figure 6B, lanes 4 and 8) and higher MW bands intensify. This is reminiscent of the weakening of the *Mm*EDEM3:*Hs*ERp46 heterodimer band in *Mm*EDEM3 mutants lacking Cys A (Figure 6B, lanes 11 and 12). This pattern indicates that loss of Cys220 disrupts normal redox cycling of the ^217^CGHC^220^ motif, shifting *Hs*ERp46 from a discrete *Mm*EDEM3:*Hs*ERp46 heterodimer into heterogeneous high-molecular-mass mixed-disulfide species. Since the anti-FLAG blot reports all *Hs*ERp46-containing species, these bands may include both *Mm*EDEM3-containing complexes and *Hs*ERp46 trapped with endogenous clients.

Equivalent results were obtained with EDEM3 KO HEK293T cells co-transfected with *Hs*EDEM3 and *Hs*ERp46 WT and trapping mutants (Supplementary Figure S18). Introduction of the C220A trapping mutation in *Hs*ERp46 resulted in disappearance of the species corresponding to the disulfide-bonded *Hs*EDEM3:*Hs*ERp46 heterodimer in a Western blot of the gel using an anti-EDEM3 antibody (lanes labelled CXXA and C220A in Supplementary Figure S18A). Enhanced laddering is also observed at higher MWs in CXXA and C220A and a mass shift upwards of the *Hs*EDEM3:*Hs*ERp46 heterodimer is not evident in the anti-ERp46-FLAG blot (Supplementary Figure S18B). Because anti-FLAG detects all ERp46-containing complexes, these species may include both EDEM3-associated complexes and ERp46 trapped with endogenous folding glycoprotein clients, consistent with the refolding role of ERp46 as a disulfide isomerase [80]. These observations are in keeping with the homology models of mammalian EDEM3:ERp46 complexes, in which the EDEM3 linker Cys A (*i.e. Hs*EDEM3 C528 or *Mm*EDEM3 C529) is disulfide linked to C217 in the *Hs*ERp46 ^217^CXXC^220^ redox motif (Supplementary Figure S17).

## Discussion

Glycoproteins constitute about a third of the eukaryotic proteome and ERAD is essential for safeguarding eukaryotic cells against the accumulation of misfolded glycoproteins [82]. The recent study of *Sc*Mnl1:Pdi1 complex established that the fungal ERAD checkpoint comprises a bifunctional complex in which the Mnl1 mannosidase preferentially recognises partially folded glycoproteins through its C-terminal domain, while the associated Pdi1 acts as the disulfide reductase that prepares committed ERAD substrates for retrotranslocation [83], although it was not clear the extent to which these conclusions extend to other EDEMs across kingdoms, and in particular to the mammalian EDEM paralogues.

Our phylogenetic analysis places fungal EDEMs close to the lineage ancestral to mammalian EDEM1 and EDEM3, whereas EDEM2 appears to have diverged following a later duplication event. Plant MNS5 and MNS4 as the respective orthologues of EDEM1/3 and EDEM2 [48]. This evolutionary framework provides the context in which the structural and functional conservation described here should be interpreted.

All experimentally characterised EDEMs function in association with a member of the PDI family, indicating that this partnership is a conserved feature of the ERAD checkpoint. In the *Sc*Mnl1:Pdi1 complex, Pdi1 surrounds the Mnl1 mannosidase domain and modulates client selection by preventing the Mnl1 C-terminal substrate-recognition domain from engaging completely unfolded polypeptides, thereby favouring misfolded globular clients [83]. However, how the EDEM:PDI association is established and regulated through disulfide-bond chemistry, why it is required for EDEM folding and catalytic competence, and whether these principles extend from fungal EDEMs to mammalian paralogues remain unclear.

Covalent association of an EDEM with a partner PDI is important to retain the EDEM in the ER. In fungal EDEM:PDI complexes and in EDEM1:PDIA1, the PDI carries a C-terminal ER-retrieval signal (“H/KDEL”, [84]), while only EDEM3 has its own ER-retrieval signal. Fungal EDEMs and animal EDEM1s are therefore kept in the ER via association with their partner PDI. There is however a second important consequence of EDEM:PDI intimate association. EDEM GH47 catalytic domains share a conserved, surface-exposed intra-molecular disulfide. In *Ct*EDEM, this corresponds to C99-C558; the equivalent cysteine pairs in *Sc*Mnl1 and mammalian EDEMs are listed in Table S1. This disulfide bond is very close to the GH47 domain glycan binding site - it clips the first *β* hairpin of the GH47 7-barrel to the last one, and in so doing it zips shut the barrel (see Supplementary Figure S13). Published observations, re-interpreted in the light of our data, suggest that the EDEM GH47 domain is mis-folded in isolation within the ER due this intramolecular disulfide bond being reduced [34, 85]. An unpaired cysteine residue at the C-terminal end of the catalytic domain (Cys A) is present in all EDEMs and it is necessary for function. Minimally, the covalent association between EDEM and PDI is mediated by the intermolecular disulfide bridge at the conserved EDEM Cys A, which binds one of the PDI redox motifs when the latter is oxidised, simply by disulfide isomerism. Its oxidation is necessary for EDEM competency/function: after reduction of SS bonds by excess glutathione *in vitro*, EDEM3 enzymatic activity can be restored by its partner PDI ERp46, suggesting that this intramolecular SS bond is in mutual redox dependency with one of the PDI CXXC motifs [42]. Equivalent data suggest the same for *Sc*Mnl1 and *Hs*EDEM2 [19, 34].

Notably, fungal EDEMs and mammalian EDEM3s possess an additional unpaired cysteine (Cys B). In the *Sc*Mnl1:*Sc*PDI1 and *Ct*EDEM:*Ct*PDI apo structures, Cys B can form a second intermolecular SS bond with the PDI (SS2). Unlike the SS1 bond, which is oxidised/present in the yeast structure and in both *Ct* EDEM:PDI structures, SS2 appears to be redox-labile. Our biochemical trapping mutant studies suggest that both Cys A and Cys B - necessary for the establishment of the intermolecular SS1 and SS2 disulfide bonds - are important in EDEM oxidation, the highest reduced/oxidised EDEM ratio pertaining to the double mutant *Mm*EDEM3 C529S/C558S which cannot form either SS1 nor SS2 (last two lanes in Figure 6A). Initial non-covalent association of EDEM and PDI could use either of the PDI’s CXXC motifs to provide the oxidising equivalents required for formation of the intramolecular EDEM disulfide, but the resulting reduced PDI motif would need to be re-oxidised in order to establish its intermolecular bond with the EDEM by disulfide isomerism. Notably, the first Cys of the *Hs*ERp46 ^217^CXXC^220^ redox motif, which is involved in the bond to Cys A, can form an intermolecular disulfide bond with peroxiredoxin 4 (PRXD4) [86, 87]. This raises the possibility that the ERp46 ^217^CXXC^220^ motif would initially oxidise the EDEM3 intramolecular SS bond; then, PRXD4, oxidised during the reduction of hydrogen peroxide to water, through a thiol-disulfide exchange could re-oxidise the reduced ERp46 ^217^CXXC^220^ motif, which in turn would enable the formation of the SS1 mixed disulfide EDEM3 Cys A-SS-ERp46 Cys217 by disulfide isomerism. Overall, PRXD4 would act as the source of oxidizing equivalents needed to drive EDEM3 intramolecular disulfide bond formation via redox chemistry at the PDI CXXC motif contributing to SS1.

Structural comparison of the *Ct*EDEM catalytic domain with the ones of other GH47 mannosidases, together with *in silico modelling* provide new insight into EDEM branch specificity. A short *α*-helix at the EDEM active site entry restricts access, ensuring specificity for mannose cleavage from branch C, the key determinant for OS-9/Yos9/XTP3-B binding. This specificity aligns with prior observations of glycan processing requirements in ERAD [43, 44]. However, client recruitment and catalytic mannose trimming are separable steps. A1AT-NHK still associates with the EDEM:PDI checkpoint complex under kifunensine treatment, indicating that the complex can engage misfolded glycoprotein clients before productive glycan cleavage. Misfolding-dependent features of the client, including exposed hydrophobic surfaces and aberrant cysteine chemistry, may therefore contribute to recruitment, whereas mannose trimming is the catalytic output that generates the ERAD lectin-recognition signal. The structures support a model in which misfolded glycoproteins are recruited primarily through the flexible IMD–PAD/PDI side of the complex through solvent-exposed hydrophobic regions, aligning with prior observations that EDEM’s flexible IMD-PAD domain dynamically engages unfolded substrates [30, 83]. The *Ct*EDEM PAD acts similarly to what has been suggested for the C-terminal domain of *Sc*Mnl1 [31]. We therefore propose that the Cys B intermolecular disulfide acts as a redox-controlled conformational switch. Reduction of the disulfide releases the IMD–PAD arm, allowing the substrate-recognition domains to explore a larger conformational space and capture misfolded glycoprotein clients. We propose that subsequent re-formation of the intermolecular SS2 disulfide could stabilise the compact, GH47-proximal conformation observed in the apo structure, keeping a PAD-bound client into proximity of the GH47 catalytic domain for demannosylation (increased avidity). The two structures define redox-linked conformational states that are consistent with (although do not by themselves establish) the proposed sequence of events. The 3D variability analysis provides a structural framework for this model: the IMD–PAD arm samples a broad range of positions relative to the GH47:PDI core, while the TRX1–3 region of *Ct*PDI also contributes to conformational variability.

The map volume corresponding to the A1AT-NHK substrate is larger than expected for a single copy of A1AT-NHK. A1AT-NHK has previously been shown to form aberrant covalent dimers, whose accumulation is suppressed by EDEM1 [55]. Thus, one possible explanation for the larger-than-expected client density is that *Ct*EDEM:PDI can engage dimeric and/or higher-order forms of A1AT-NHK. Trapping-mutant Western blots further suggest that *Ct*PDI and ERp46 redox chemistry can generate transient mixed-disulfide species with A1AT-NHK. PDI:substrate interactions, essential for PDI chaperone function [86, 87], may therefore be preserved in the context of the ERAD checkpoint complex. In particular, the non-reducing Western blots of protein purified from cells co-transfected with *Ct*EDEM A1AT-NHK and the *Ct*PDI Cys53Ser trapping mutant suggest that PDI Cys50 can be trapped either in a disulfide with CtEDEM Cys B, or in a mixed disulfide species with the client. These data support reversible redox exchange at the Cys B–PDI redox motif and indicate that *Ct*PDI can transiently engage ERAD clients through mixed disulfides. In the *Ct*EDEM:*Ct*PDI:A1AT-NHK structure, the Cys B–PDI disulfide is reduced, indicating that this second intermolecular disulfide is redox-active during client engagement. The ability of trapping mutants in the corresponding PDI redox motif to form disulfide-linked species with clients further supports reversible redox chemistry at the Cys B site. Disappearance of the *Mm*EDEM3:*Hs*ERp46 and *Hs*EDEM3 :*Hs*ERp46 heterodimer bands in non-reducing SDS-PAGE gels when *Hs*ERp46 Cys220 is mutated is in keeping with mammalian EDEM3:ERp46 homology models, which show a disulfide bond between EDEM3 Cys A and *Hs*ERp46 Cys217 in the ^217^CGHC^220^ CXXC motif^6^. In absence of externally supplied oxidising equivalents, establishment of an intermolecular disulfide between an EDEM free Cys and the first Cys of an ERp46 CXXC motif takes place via disulfide bond isomerism: in this scenario, formation of the EDEM3 Cys A disulfide bond with *Hs*ERp46 Cys217 requires an intact *Hs*ERp46 ^217^CXXC^220^ motif. The enhanced high-molecular mass laddering observed whenever Cys220 is mutated is compatible with an increase in disulfide-bridged ERp46-SS-client complexes. Within the ^217^CGHC^220^ motif, Cys217 can form mixed disulfides with EDEM3 Cys A or with client cysteines, whereas Cys220 provides the resolving cysteine that allows intramolecular closure of the ERp46 redox motif and release of the covalent partner. The Cys217–Cys220 intramolecular disulfide may therefore act as a redox-reset step, allowing ERp46 to exit non-productive mixed-disulfide adducts and re-enter either EDEM3 association or client handling. Loss of Cys220 shifts ERp46 away from a discrete EDEM3:ERp46 heterodimer and into high-molecular-mass ERp46–SS–client and/or ERp46–SS–EDEM3-containing species.

Beyond ERAD substrate processing in general, our findings have implications for protein misfolding disorders such as *α*1-antitrypsin deficiency (A1ATD) [6, 89]. Homozygosity for the A1AT-Z variant allele, present in 2-4% of populations of North European ancestry, is responsible for the great majority of clinically-significant disease. An important minority of A1AT-Z polypeptide chains form ordered, polymeric aggregates that are retained within the ER and may have toxic gain-of-function consequences in the liver [6]. However, more than 50% of A1AT-Z is - like the NHK variant - recognised as terminally misfolded and undergoes ERAD [90, 91]. This accounts for the majority of the circulating deficiency and hence loss-of-function underlying the concomitant risk of early-onset emphysema. Disrupting EDEM:misfolded glycoprotein interactions - either by modifying hydrophobic engagement or by interfering with disulfide bond formation - could provide therapeutic strategies to mitigate the loss-of-function effects associated with A1ATD [92–95].

Together, our results clarify key mechanistic roles of the EDEM:PDI checkpoint complex in ERAD and set the foundation for targeting these interactions in protein misfolding diseases. By linking structural insights with functional validation, we provide a framework for future therapeutic strategies aimed at modulating ERAD activity in disease contexts.

## Methods

### Phylogeny

Canonical EDEM reference sequences (*Hs*EDEM1: Q92611, *Hs*EDEM2: Q9BV94, *Hs*EDEM3: Q9BZQ6, *Ct*EDEM: G0SCX7, *Sc*MNL1: P38888, *At*MNS4: Q9FG93, *At*MNS5: Q9SXC9) were used to retrieve from UniProtKB orthologous EDEM sequences with at least 50% of global sequence identity. Full-length sequences were trimmed to the boundaries annotated by the InterPro record IPR044674. The Hidden Markov Model (HMM) defining this domain derives from the PANTHER database profile PTHR45679. To avoid topology artefacts from uneven taxon sampling, EDEM1, EDEM2, and EDEM3 sequences were constrained to the same 40 vertebrate species, selected via stratified sampling across classes from the intersection of groups possessing a *bona fide* GH47 domain. Similarly, *Arabidopsis thaliana* MNS4 and MNS5 were complemented with orthologues from other plant species (35 in total), while fungal datasets comprised up to sequences from a total of 55 selected filamentous *Ascomycota* and *Saccharomycotina* species. To ensure high taxonomic diversity and prevent over-representation of individual lineages, plant and fungal species selection was capped at two species per genus. Finally, all newly retrieved plant and fungal sequences were pre-filtered to within 25% of their reference sequence length to guarantee strict structural homology.

**Multiple sequence alignment** The extracted sequences were aligned using MAFFT v7.526 [96]. Algorithm: G-INS-i --globalpair was selected because it incorporates global pairwise alignment information and it is particularly suitable for sequences of similar length exhibiting conserved global homology. Alignments used the BLOSUM62 scoring matrix with default gap penalties (gap opening: −1.53, gap extension: 0.0), and the iterative refinement was set to a maximum of 16 iterations (--maxiterate 16) to ensure convergence. Alignment columns containing more than 50% gaps were removed prior to phylogenetic inference using trimAl v1.5 (-gt 0.5) [97], reducing the alignment to 482 columns. The trimmed alignment contained 422 parsimony-informative sites (87.6%) and 34 constant sites (7.1%).

**Phylogenetic inference** IQ-TREE 2 [98] was used to select the best-fit substitution model, with ModelFinder Plus (-m MFP), 1,000 ultrafast bootstrap replicates (-bb 1000) and 1,000 SH-aLRT replicates (-alrt 1000). The optimal model, chosen by Bayesian Information Criterion (BIC) was LG+R7 (log-likelihood = ×44,816.39), incorporating a discrete free-rate distribution with seven rate categories. The analysis was run on 255 sequences aligned across 482 columns. The resulting phylogenetic tree was visualized using the iTOL Tree Server [99].

### Molecular cloning

The mature *Ct*EDEM sequence (from which the N-terminal portion residues 26-63 were omitted) was cloned into the pHLsec vector for secreted expression in mammalian cells: the construct has an N-terminal GFP fusion, followed by a 3C-protease linker (LEVLFQGP), *Ct*EDEM residues 64-1092 and a C-terminal hexa-histidine tag. *Ct*EDEM residues 26-63 were omitted from this construct in that they are predicted to be membrane-associated. The mature *Ct*PDI sequence (from which the C-terminal KDEL ER-retrieval motif was deleted, in order to enable protein purification from the mammalian cells supernatant) was also cloned in the same vector. The *Ct*PDI:pHLsec plasmid expresses the mature *Ct*PDI residues 21-515 (omitting the C-terminal ^516^HDEL^519^ ER retrieval motif) followed by a 6xHis affinity tag (see Figure S5). The molecular mass of *Ct*PDI is 55 kDa, the one of the *Ct*EDEM fusion construct is 142 kDa so the combined MW of a complex is 197 kDa; adding the putative 7+3=10 *N*-glycosylation sites and using an estimated average of 2.0 kDa per *N*-linked glycan [100], the *Ct*EDEM:PDI heterodimer we cloned is expected to have an approximate mass of 220 kDa.

The DNA sequence encoding *Ct*EDEM (UniProt G0SCX7) was obtained from GeneArt. *Ct*PDI DNA (UniProt G0SGS2) was amplified from *Ct* cDNA (a gift of Ed Hurt, Biochemistry Centre, Heidelberg University, Heidelberg, Germany). Genes were amplified via PCR using Q5 High-Fidelity Polymerase. Vectors used were Litmus28i for cloning and pHLsec for mammalian expression. Litmus28i was linearized using EcoRI and XhoI, while pHLsec was linearized with AgeI and KpnI.

Primers with 5’overlaps tailored for Gibson assembly were designed. For *Ct*EDEM, a full-length codon-optimized gene was cloned into Litmus28i and pHLsec vectors. The N-terminal GFP fusion construct was generated by amplifying GFP and *Ct*EDEM fragments, followed by Gibson assembly into the pHLsec vector. The DNA sequence encoding for GFP protein was amplified using the plasmid pHLsec:GFP-SLC35B1 (UniProt ID P78383), kindly donated by Dr Snezana Vasiljevic (Biochemistry Department, University of Oxford). For *Ct*PDI, intron removal was achieved by amplifying the two exons separately, excluding the signal peptide and HDEL retrieval signal, and cloning them into pHLsec using Gibson assembly. Gibson assembly reactions were carried out with a 3:1 insert:vector molar ratio and incubated at 50 ° C for 1 hour. Site-directed mutagenesis of *Ct*PDI was carried out following a protocol outlined in [101]. DNA bands were verified via gel electrophoresis and purified when necessary using the GeneJet Gel Extraction Kit.

Competent DH5*α E. coli* cells were transformed with Gibson assembly products via heat shock and plated on LB agar containing carbenicillin (100 µg/mL). Colony-PCR was performed to confirm successful transformations. Plasmids were purified using GeneJet Miniprep Kits for small-scale preparations and an adapted alkaline lysis protocol for large-scale maxipreps [102]. Purified plasmids were quantified via NanoDrop.

### HEK293T transient transfection and Western blot analysis

EDEM3 KO of Human embryonic kidney (HEK293T) cells were cultured in DMEM supplemented with 10% FBS and transfected at 65–70% confluence using Polyethylenimine (PEI, 2.5:1 [v/w, PEI:DNA]). Cells were co-transfected with plasmids encoding A1AT-NHK :pCDNA3.1, *Ct*EDEM:pHLsec, *Ct*PDI:pHLsec, *Mm*EDEM3:pcDNA3.1, *Hs*ERp46:pcDNA3.1 controls, using optimized DNA ratios (*e.g.*, 1:1:1 for *Ct*PDI:*Ct*EDEM:NHK triple expression or 3:1 for *Mm*EDEM3:*Hs*ERp46 co-expression). For *Hs*ERp46 trapping mutant experiments, cells were treated with 10 mM iodoacetamide prior to sampling.

Cells were lysed in Triton X-100 lysis buffer supplemented with protease inhibitors, phosphatase inhibitors, and 20 mM N-ethylmaleimide. Lysates were clarified by centrifugation at 14,000 × g for 20 minutes at 4 ° C, and protein concentrations were measured using the BCA assay. Equal protein amounts were separated by SDS-PAGE under reducing or non-reducing conditions (6–10% gels depending on the target) and transferred to 0.45 µm nitrocellulose membranes.

Membranes were blocked with 5% milk in TBS and incubated overnight at 4°C with primary antibodies (*e.g.*, anti-A1AT, anti-His, anti-HA, anti-Flag, anti-EDEM3) diluted in 2.5% BSA in TBS. After washing, HRP-conjugated secondary antibodies were applied, and signals were detected using ECL substrate and a ChemiDoc Imaging system (Bio-Rad). Band intensities were quantified with ImageJ (v. 2.1.0) and normalized to the empty vector (EV) control. Statistical analysis was performed using one-way ANOVA with Dunnett’s multiple comparisons test in GraphPad Prism v. 10.1.1. Error bars represent the standard deviation of three independent experiments.

### Protein purification

HEK293F cells were co-transfected using the method described in [103], with equal amounts of pHLsec:*Ct*EDEM and pHLsec:*Ct*PDI.

***Ct*EDEM:*Ct*PDI** The HEK293F cell culture was harvested 4 days post-transfection by centrifuging at 4,000 g for 5 minutes. The supernatant containing the secreted protein was adjusted to 1 x PBS pH 7.4 and filtered prior to IMAC. The filtered supernatant was passed onto a 1 mL HisTrap HP Ni IMAC column equilibrated against binding buffer (Na_2_HPO_4_ 50 mM pH 7.5, NaCl 300 mM). The column was washed with 5 column volumes (CV) of wash buffer (Na_2_HPO_4_ 50 mM pH 7.5, NaCl 300 mM, 10 mM imidazole). The proteins were eluted with a 15 CV gradient from 0% to 100% of elution buffer (Na_2_HPO_4_ 50 mM pH 7.5, NaCl 200 mM, 500 mM imidazole.

The protein-containing fractions from IMAC were pooled and concentrated using a polyethersulfone (PES) membrane, 10 kDa MW cutoff centrifugal ultrafiltration device. This sample was then filtered through a 0.2 *µ*m filter and applied to either a Superdex 200 Increase 10/300 GL/Superdex 200 16/600 pg (SEC) or Superdex 200 3.2/300 (microSEC) equilibrated against SEC buffer (HEPES 20 mM pH 7.5, NaCl 100 mM). SDS-PAGE of SEC fractions was used to assess purity (Thermo Fisher NuPAGE Bis-Tris 4-12% gel) and identify fractions of interest for downstream biochemical, structural, and proteomic studies.

***Ct*EDEM:*Ct*PDI:A1AT-NHK** For expression of the ternary complex, three plasmids were transfected in a mass ratio for PDI:EDEM:A1AT of 2:4:3 to a final DNA concentration of 1 µg/mL. For A1AT co-transfections, kifunensine (Merck - K1140, Germany) was added to a final concentration of 1 µg/mL. After 72-96 hours of expression, the cells were harvested and the supernatant was retained. The supernatant was adjusted to 1xPBS and made up to final 10 mM imidazole. The pH was adjusted to 7.4 and the sample was filtered using a 0.22 µm polyethersulfone membrane filter (Merck - GPWP02500, Germany). The filtered supernatant or lysate of the HEK293F cells was passed onto 5 mL HisTrap HP IMAC column (Cytiva - 29051021/17524801, USA) equilibrated against binding buffer (1xPBS 10 mM imidazole) using a peristaltic pump at room temperature, at a flow rate of approximately 2 mL/min. The column was washed with 5 CV of binding buffer, fitted to an Å KTA purifier FPLC (Cytiva, USA) in a 4 ° C cold cabinet. The column was washed a second time with a wash buffer (1xPBS 25 mM imidazole) for 5 CV or until the UV trace had returned to baseline. The proteins were eluted with a 15 CV gradient from 0% to 100% of elution buffer (1xPBS 300 mM imidazole).

The protein-containing fractions from IMAC were pooled and concentrated using a 0.22 µm polyethersulfone (PES) membrane, 10 kDa MM cutoff centrifugal ultrafiltration device (Thermo Fisher Scientific - 88513, USA). This sample was then filtered through a 0.22 µm filter (Merck - GPWP02500, Germany) and applied to a Superdex 200 Increase 10/300 GL (SEC, Cytiva - 28990944, USA) equilibrated against SEC buffer (HEPES 20 mM pH 7.5, NaCl 100 mM). Sample volume was 0.5 mL for SEC. Fractions containing the complex were identified from the SEC chromatogram and SDS-PAGE and taken forward for Cryo-EM. Concentration of each fraction was measured using the A280 nm reading.

### Cryo-electron microscopy grid preparation

***Ct*EDEM:*Ct*PDI grids** The *Ct*EDEM:PDI Cryo-EM grids were prepared with a *Ct*EDEM:PDI sample eluted from a Micro-SEC column (Supplementary Figures S5 and S6). Quantifoil Au300 R1.2/1.3 Holey Carbon grids were glow discharged using the EMS GloQube. For non-graphene oxide grids, settings at 0.2 mBar, 35 mA for 60 seconds were used. For grids to be coated in graphene oxide, settings at 0.2 mBar, 50 mA for 75 seconds. Graphene oxide grid coating was carried out according to the protocol in [104]. Grids were blotted at blot force 10 for 3 seconds before plunge freezing immediately into liquid ethane using a Vitrobot Mark IV (Thermofisher, UK).

***Ct*EDEM:*Ct*PDI:A1AT-NHK grids** The Quantifoil Au300 R1.2/1.3 Holey Carbon grids were glow discharged using the EMS GloQube (Quorum Tech, UK). The Holey Strong setting was used (0.2 mBar, 35 mA for 60 seconds) for non-graphene oxide grids. For grids to be coated in graphene oxide (GO), the Graphene Oxide setting was used (0.2 mBar, 50 mA for 75 seconds). For GO coating, a graphene oxide solution (Sigmaaldrich, Germany) at 0.2 mg/ml was spun at 300 g for 30 s to remove large aggregates. 3 µL of protein were added to each grid for 1 minute, at 4 ° C with 100% humidity, before blotting with Whatman No1 filter paper (Sigma-aldrich, Germany) and washing three times with 20 µL drops of ultrapure water. Specimens were frozen using an FEI Vitrobot Mark IV (Thermofisher, UK). Holey carbon grids were blotted at blot force 10 for 3 s before plunge freezing immediately into liquid ethane. Vitrification with GO grids included a 30 second wait time after sample application before plunge freezing.

### Cryo-electron microscopy data collection

Cryo-EM data collection was carried out on a FEI Titan Krios microscope with Gatan K3 detector at the Midlands Cryo-EM facility using the following settings: acceleration voltage was set to 300 kV; spherical aberration coefficient (Cs) was 2.7 mm, amplitude contrast 0.1. The FEI Titan Krios Gatan K3 detector was set to super-resolution counting mode bin 2, nominal magnification was 105,000x; calibrated pixel size was 0.835 Å; the second condenser aperture was set to 50 µm; the objective aperture was set to 100 µm; the defocus range was from −2.6 µm to −0.7 µm in 0.3 µm intervals; the energy filter slide width was 20 eV; the dose rate on the detector was measured over a typical specimen area and was 15 electrons/pixel/s; the dose rate on the specimen was measured over a hole and was 16.5 electrons/pixel/s; the exposure time was 3 sec/micrograph.

***Ct*EDEM:*Ct*PDI Cryo-EM data processing** Data processing was carried out in RELION 4.0 [105] and cryoSPARC v4.1 [106], with the final *Ct*EDEM:PDI reconstruction using cryoSPARC v4.1. In RELION, Motion correction used MotionCor2 [107], CTF estimation used CTFFind4 [108] and particle picking was carried out using Topaz [109] and autopicking software. In cryoSPARC, patch motion correction and patch CTF estimation were used, and particle used 2D templates from RELION for template picking. 2D classification was carried out in multiple rounds, each time removing particles that gave rise to poor quality classes. *Ab-initio* reconstruction was used to generate multiple initial 3D reconstructions. The various reconstructions were used alongside manufactured junk reconstructions to find and remove further junk particles. 3D classification was used to investigate structural heterogeneity within the dataset, and split particles into subsets. The subsets were refined using homogenous refinement and non-uniform refinement. Reconstructions were improved with local and global CTF refinements. Particle subtraction and local refinements were carried out by first generating masks against the regions of interest using Create Mask or Volume Tools in UCSF Chimera [110]. 3D variability analysis (3DVA) [111] was carried out using a consensus map from non-uniform refinement. 3DVA was run with 3 modes and filter resolution at 6Å. Output mode was set to simple with 20 components. The main components of the three modes were visualized and recorded as movies in Chimera v1.9 using Volume Series.

***Ct*PDI:*Ct*EDEM:A1AT-NHK Cryo-EM data processing** Movies were pre-processed as described for the *Ct*PDI:*Ct*EDEM dataset, and the same default settings used for 2D classification, producing an *ab-initio* model; 3D classification, and 3D refinement were used unless described otherwise (see section 18). After pre-processing 7,098 micrographs remained from 7,773 movies.

An untrained Topaz autopick was carried out on the remaining micrographs, with particle diameter set to 160 Å. Particles were extracted using Figure-of-Merit threshold at −2, at a box size of 336 pixels, re-scaled to 84 pixels (4x binning). This extracted 3,622,266 particles which were cut down to 109,167 particles over 8 rounds of 2D classification, 4 at 4x binning. and another 4 at 2x binning. Whilst re-extracting at 2x binning, the box size was increased to 432 pixels. Mask diameter for 2D classification was also increased to 200 Å. The good particles were used as input to train Topaz on this particular dataset with number of particles set to 300 per micrograph. The same process was then repeated with particle picking, using the trained model, particle extraction (432 pixels) and rounds of 2D classification to remove junk particles. After 11 rounds of 2D classification, there were 912,174 particles remaining in 36 classes.

Particles from the classes which displayed extra density were selected with a total of 388,835 particles. These particles were used to generate two initial models, and the one which displayed the features most resembling the *Ct*PDI:*Ct*EDEM heterodimer was taken forward to use as a reference for 3D classification. Four 3D classes were generated, one of which contained very few particles, while the remaining particles were split fairly evenly between the other three. The classes looked very noisy with strong preferential orientation.

To try and improve the alignment of the particles in the region for which the structure is known, a mask was generated around the core *Ct*PDI:*Ct*EDEM particle. The mask was generated by creating initial models from the particles in 2D classes which appeared as single particles with no extra density. Focused 3D classification using the core particle mask produced improved 3D classes. The particles and map of the best class was taken forward for a 3D refinement.

### Model building

Homology models of *Chaetomium thermophilum* protein disulfide isomerase (*Ct*PDI) and ER-degradation enhancing alpha-like mannosidase protein (*Ct*EDEM) in com-plex were generated using Alphafold-Multimer [112]. Sequences were obtained from UniProt (*Ct*PDI: [G0SGS2], residues 21–505; *Ct*EDEM: [G0SCX7], residues 64–1092) and processed with standard AlphaFold3 parameters. Disordered regions with low confidence (local distance difference test score ≤ 75) were removed using a custom pruning script, yielding truncated models for docking into the EM map.

The Cryo-EM density maps were generated in cryoSPARC and truncated AlphaFold2 models were initially docked into the Cryo-EM map by phased molecular replacement using CCP4-Molrep [113], followed by iterative model building and refinement. docked and refined with Phenix [114]. Models were rigid-body fitted with Phenix.dock in map, and real-space refinements were performed with Phenix.real space refine [115] using secondary structure and Ramachandran restraints. Missing loops were added iteratively with Phenix.fit loops and manually built in Coot [116, 117] where required. Glycosylation sites were modeled using Coot’s carbohydrate module, followed by refinement in Phenix.

The Coot validation tools Validate/Ramachandran Plot, Validate/Rotamer analysis and Validate/Validation Outliers were used to improve errors in the structure. Favoured side chain rotamers were selected using the Rotamers tool. Real-space fit in Coot was carried out with Ramachandran restraints and the automated weight calculated by the program for the Cryo-EM map. A final pass using the model tools of Isolde [118] ironed out a few remaining errors.

The final apo structure model (PDB ID 9TPA) analysed in MolProbity [119] has RMSD_bonds_= 0.016 Å and RMSD_angles_= 0.9^◦^; 7.2% (1433/1439) of all residues are in favored (98%) regions and 2% (33/1439) of all residues were in allowed regions in the Ramachandran plot. The final A1AT complex structure model (PDB ID 29TE) analysed in MolProbity [119] has RMSD_bonds_= 0.002 Å and RMSD_angles_= 0.539^◦^; 94.3% (1369/1451) of all residues were in favored (98%) regions; 100.0% (1451/1451) of all residues were in allowed (*>*99.8%) regions.

### In silico docking

**Guided docking protocol** Modeling high-mannose glycans presents a significant computational challenge; therefore, to ensure predictive accuracy, we established a guided docking protocol integrating knowledge-based energetic rewards to favour the identification of biologically relevant binding poses and it is implemented via the AutoDock Vina Site (VS) (code available at https://gitlab.com/modenuttilab/vina-site) [120]. The protocol was tuned using the crystal structure of human ER alpha-mannosidase I (ERManI) in complex with Man_9_GlcNAc_2_ pyridylamine (PDB ID 5KIJ) as a benchmark [76]. Parameters were iteratively optimized to reproduce the crystallographic pose of the Man_8_GlcNAc_1_ isomer A (M8A) in the crystal structure (Supplementary Figure **??**).

The model was built using the Carbohydrate Builder from GLYCAM-web [121] and prepared with the AutoDock Tools package from MGLTools 1.5.7 [122]. The model includes the catalytic Ca^2+^ ion and three structural water molecules (HOH 871, HOH 967, and HOH 1000) structurally equivalent to those implicated in the catalytic mechanism of the glycosyl hydrolase family 47 (EC 3.2.1.113) in the Catalytic Site Atlas [74, 123]. These water molecules serve as steric shields to prevent the formation of non-native contacts during docking.

Prior structural studies have reported CH–*π* interactions between aromatic residues and glycans in GH47 enzymes. For example, in mouse Golgi *α*-mannosidase IA a tryptophan residue at the edge of the substrate-binding groove contacts either a mannose unit (PDB ID 5KKB, [76]) or a GlcNAc moiety of the glycan (PDB ID 1NXC, [73]), while in an ERManI structure (PDB ID 5KIJ, [76]) analogous interactions are observed between a tryptophan residue and a GlcNAc moiety of the glycan. Energetic rewards to favour such CH–*π* interactions were defined for exposed aromatic residues in the cavity (W284 and W389), alongside specific polar rewards for the amino acids that stabilise the terminal mannose, such as R597, E599, E663, T688 and E689 (see Supplementary Figure S22).

Five independent docking runs with a high exhaustiveness setting of 80 and default CHI parameters, yielded a total of 150 predicted binding poses. Conformations were clustered based on the Root-Mean-Square Deviation (RMSD) of glycan ring atoms (C1, C2, C3, C4, C5, and O5) with a 2.0 Å cutoff [124], using the TCL script qt clustering-pdb 2.0.tcl (available at https://gitlab.com/modenuttilab/vina-site). Only ligand poses that optimized the scoring function’s energetic rewards were retained for visual inspection. The representative binding pose was identified as the top-ranked conformation that also exhibited the lowest RMSD relative to the reference crystallographic structure.

**M8B docking** We modelled the interaction between *Ct*EDEM and the GlcNAc_1_Man_8_ isoform B glycan (M8B). To inform the docking setup, we examined available GH47 structures. Our apo *Ct*EDEM:*Ct*PDI structure contains a D-mannose bound in the catalytic site corresponding to the product of the reaction. This mannose defines the geometry of the −1 subsite and was used as a template to position the terminal *α*1,2-linked mannose from branch C of the M8B glycan (mannose “K”) in the catalytic site.

In particular, in the apo *Ct*EDEM structure, residue D370 stabilizes the O2 atom of the D-mannose, while T608 coordinates both the Ca^2+^ ion and the O3 atom. E609 forms a contact with O4, and R520 and E522 interact with O5 and O6, respectively. These residues are highly conserved and form part of the canonical GH47 (*α/α*)_7_-barrel active site at the base of the barrel. These interactions were used to guide the docking procedure by prioritizing placement of the terminal *α*1,2-linked mannose from branch C (mannose “K” in Supplementary Figure S1) in a catalytically relevant position.

Crystal structures of GH47 *α*-mannosidases in complex with glycans were also analysed, including human ER *α*-mannosidase I (PDB ID 5KIJ [76]), mouse Golgi *α*-mannosidase IA (PDB ID 5KKB [76]), *Saccharomyces cerevisiae* class I *α*-1,2-mannosidase (PDB ID 1DL2 [62]), and mouse Golgi *α*-1,2-mannosidase IA (PDB ID 1NXC, [73]). In these structures, the conserved residue pair E501 and R520 establishes contacts with hydroxyl groups of bound glycans, and contacts between these residues and glycan OH groups were therefore included as bias restraints. A small number of structural water molecules observed in the crystal structures were also included explicitly in the docking protocol, based on their conservation and their interactions with residues structurally equivalent to *Ct*EDEM Q188, E191, R195, and E576.

Specific interactions were incorporated into the guided docking protocol AutoDock Vina Site (VS) described above [120]. Docking calculations were carried out with AutoDock Vina Site (VS) 1.0, a modified version of AutoDock Vina incorporating the CHI functions described in [125]. These functions penalize unfavourable glycosidic dihedral angles while favouring chemically plausible carbohydrate conformations. In addition, VS enables spatial restraints to guide placement of ligand functional groups, thereby enhancing sampling of interactions identified as important. The output reports glycosidic torsional angles and corresponding CHI penalties, allowing assessment of carbohydrate conformational plausibility.

Docking of the M8B glycan to the GH47 domain of *Ct*EDEM was initially performed without restraints on aromatic–carbohydrate interactions. Prior structural studies have reported CH–*π* interactions between aromatic residues and glycans in GH47 enzymes. For example, in mouse Golgi *α*-mannosidase IA a tryptophan residue at the edge of the substrate-binding groove contacts either a mannose unit (PDB ID 5KKB, [76]) or a GlcNAc moiety of the glycan (PDB ID 1NXC, [73]), while in an ERManI structure analogous interactions are observed between a tryptophan residue and a GlcNAc moiety of the glycan (PDB ID 5KIJ, [76]).

In these unbiased runs, poses consistently formed CH–*π* interactions between the GlcNAc moiety and Trp516, whereas no such interactions were observed with other solvent-exposed aromatic residues (*e.g.* Trp461).

Based on these observations, and in addition to the polar and metal-coordinating interactions described above, the final rounds of guided docking included a soft restraint favouring CH–*π* interactions between the GlcNAc moiety and solvent-exposed aromatic side chains. This restraint was applied in a general manner, without specifying a particular atom of the sugar ring, thereby allowing any geometrically compatible CH–*π* interaction to form. The purpose of this bias was to focus sampling on chemically plausible carbohydrate–protein interactions, rather than to impose a specific binding geometry, and it was implemented as a soft restraint rather than a fixed positional constraint.

Five independent docking runs were performed, generating 30 poses per run. The resulting conformations were clustered using the RMSD of glycan ring atoms (C1, C2, C3, C4, C5, and O5) with a 2.0 Å cutoff, as described in [124]. CHI parameters were set to chi coeff = 1 and chi cutoff = 2, and docking exhaustiveness was set to 80 to ensure sufficient exploration of conformational space.

### Homology modelling

Searches in the *Ct* genome using the search tool on the server at https://c-thermophilum.bork.embl.de/ and the protein sequences of *Sc*Mnl1 (*Sc*Mnl1 ((UniProt P38888) and *Sc*PDI1 (UniProt P17967)) yielded UniProt entry https://www.uniprot.org/uniprot/G0SCX7 (hereinafter *Ct*EDEM), annotated as *α*-1,2-mannosidase and https://www.uniprot.org/uniprot/G0SGS2 (hereinafter *Ct*PDI), annotated as a protein disulfide isomerase. In agreement with the phylogenetic analysis, the same UniProt entries were hit by searching the *Ct* genome with the sequences of *Hs*EDEM3 [46] and its partner PDI *Hs*ERp46[42], respectively. The same top hits for the *Ct*EDEM and *Ct*PDI found proteins annotated as ERAD EDEMs and PDIs using BlastP [126].

Homology models of mammalian EDEM:PDI complexes were generated using the Alphafold3 server [127]:

1. *Hs*EDEM1:P4HB *aka* PDIA1: *Hs*EDEM1, UniProt Q92611, residues 112-596 and 626-657; *Hs*P4HB *aka* PDIA1: UniProt P07237, residues 21-476. Overall statistics: *<*pLDDT*>*=85.14. ipTM = 0.85, pTM = 0.76 [128];
2. *Hs*EDEM3:ERp46:EDEM2:TXNDC11: *Hs*EDEM3, UniProt Q9BZQ6, residues 43-796; *Hs*ERp46, domains TRX1 and TRX2, UniProt Q8NBS9, residues 60-298; *Hs*EDEM2, UniProt Q9BV94, residues 29-578; *Hs*TXNDC11, UniProt Q6PKC3 residues 91-265, 292-422, 594-799. Overall statistics: *<*pLDDT*>*=85.39. ipTM = 0.81, pTM = 0.79 [128]. This Alphafold3 model of the *Hs*EDEM3:ERp46:EDEM2:TXNDC11 heterotetramer is compatible with mobility of the EDEM3 C-terminal arm enabling client transfer between the sequentially acting EDEM2 and EDEM3 catalytic sites: both EDEM2 and EDEM3 catalytic sites in the heterotetramer model are within reach of the C-terminal mobile arm of EDEM3. Association with EDEM3:ERp46 would enable ER retrieval of EDEM2:TXNDC11, both components of which lack a KDEL-like retrieval signal.

### Plant biology

**Plant material and growth conditions** *A. thaliana bri1-5* mutant and *mns4-1 mns5-1 bri1-5* triple mutant [51] were grown on soil at 22 ^◦^C and 70% relative humidity, under a 16-h-light/8-h-dark cycle (approximately 120 *µ*mol m^−2^ s^−1^).

**Cloning of *Ct*EDEM:mRFP** The full-length coding sequence of *Ct*EDEM was amplified by PCR using the Accuprime Pfx DNA Polymerase (Invitrogen), cloned into pDONR^™^ 221 vector (Invitrogen) and sequenced. DNA primers used for cloning are “*Ct*EDEM pDONR^™^ 221” in Table S4. Full-length *Ct*EDEM was then cloned using the Gateway^®^ Technology into two destination vectors pB7RWG2 and pK7WGF2, fused at the C-terminus with monomeric RFP (mRFP) and enhanced GFP (eGFP), respectively, according to the manufacturer’s instructions.

Binary vectors, containing the constructs, were amplified in *Escherichia coli* DH5*α* before transformation of *Agrobacterium tumefaciens* (strain GV3101).

**Plant stable/transient transformation** Later, the constructs *Ct*EDEM:mRFP and *Ct*EDEM-eGFP were used for stable transformation of the *A. thaliana mns4-1/mns5-1/bri1-5* triple mutant. *Arabidopsis* plants were transformed using the floral dip method [129]. Healthy *Arabidopsis* plants were grown until they were flowering. *Agrobacterium tumefaciens* carrying gene *Ct*EDEM was spun down and resuspended to OD_600_=0.8 in 500 mL of 5% Sucrose, 10 mM MgCl_2_, 200 *µ*M acetosyringone. Before dipping, Silwet L-77 was added to a concentration of 0.05% (500 *µ*L/L). *At* was dipped in *Agrobacterium* solution for 1 minute and kept under cover for 48 hours to maintain high humidity. After 5 days, the dipping procedure was repeated. Dry seeds were harvested and the transformants were selected on 1/2 MS medium with phosphinothricin (aka Basta, Duchefa P0159) for *Ct*EDEM:mRFP plants. With kanamycin (Duchefa K0126) for *Ct*EDEM:eGFP plants. Protoplasts were prepared from *Ct*EDEM:mRFP 2-week-old stable transformed leaves and transformed with 20 *µ*g of GFP:KDEL plasmid vector following the protocol reported in Leucci et al., 2007 [130].

**Confocal Laser Scanning Microscopy and *Ct*EDEM expression and localisation *in planta*** Confocal images were taken by a laser-scanning confocal microscope (LSM Pascal; Carl Zeiss). *Ct*EDEM-eGFP and eGFP:KDEL signal was excited at 488 nm with an argon laser and detected with a 505–530 nm filter set; whereas *Ct*EDEM:mRFP was excited at 543 nm with a He/Ne laser and detected in the 560–615 nm range. Chlorophyll autofluorescence was detected with the filter *>* 650 nm.

## Acknowledgements

We thank Radu Aricescu and Ed Hurt for donating the pHLsec vector and the *Ct* cDNA library, respectively. We acknowledge help with Cryo-EM sample preparation, data measurement and processing from C.G. Savva and T.J. Ragan. Taha Shahid, Kerry Blair, Claudia Lancey and Alfredo De Biasio helped with protein production and Cryo-EM grid preparation. This work benefited from access to facilities at Diamond Light Source and the Research Complex at Harwell (RCaH), both of which are Instruct-ERIC Centres.

## Data availability

Structure coordinates were deposited to the Research Collaboratory for Structural Bioinformatics PDB (https://www.rcsb.org/) and Cryo-EM density maps used in model building were deposited to the EM Data Bank (https://www.ebi.ac.uk/pdbe/emdb) under the following accession codes: 9TPA and EMD-56105 (apo form); 29TE and EMD-57361 (complex with A1AT-NHK). Other data supporting the findings of this study are available from the corresponding authors on request.

## Author contributions

C.J.H. and A.L. contributed equally to this work. P.R., A.S., B.G. and N.H. conceived and resourced the study. P.R. and A.L. cloned the *Ct*EDEM and *Ct*PDI DNA constructs. P.R., G.T., M.Bh., G.M.P., and C.J.H. purified protein. C.J.H. prepared Cryo-EM grids. C.J.H. collected Cryo-EM data. C.J.H. and M.Bh. processed Cryo-EM data. A.S., M.D.B. and A.L. carried out the experiments with *Ct*EDEM in plants. C.J.H, B.G, J.B., M.Bh. and A.Q. carried out the A1AT-NHK ternary complex Cryo-EM studies and conducted the *Ct*PDI trapping mutant studies. P.R., Y.B., I.C., M.Bh., G.M.P. and C.J.H. built and refined the structural models. S.G., S.P., G.N.C., C.V.A.M., I.W. and N.H. carried out the mammalian EDEM3 and ERp46 experiments. P.R., N.H., I.W., C.J.H. and C.V.A.M. planned the MS experiments. J.R.O. and C.P.M. carried out the phylogenetic analysis. J.L., P.R. and C.P.M. carried out the *in silico* docking. P.R., L.S., D.L.A., D.D.B. and M.B. carried out *in vitro* assays. All authors contributed to the writing of the manuscript.

## Extended Data

**Figure S1.**
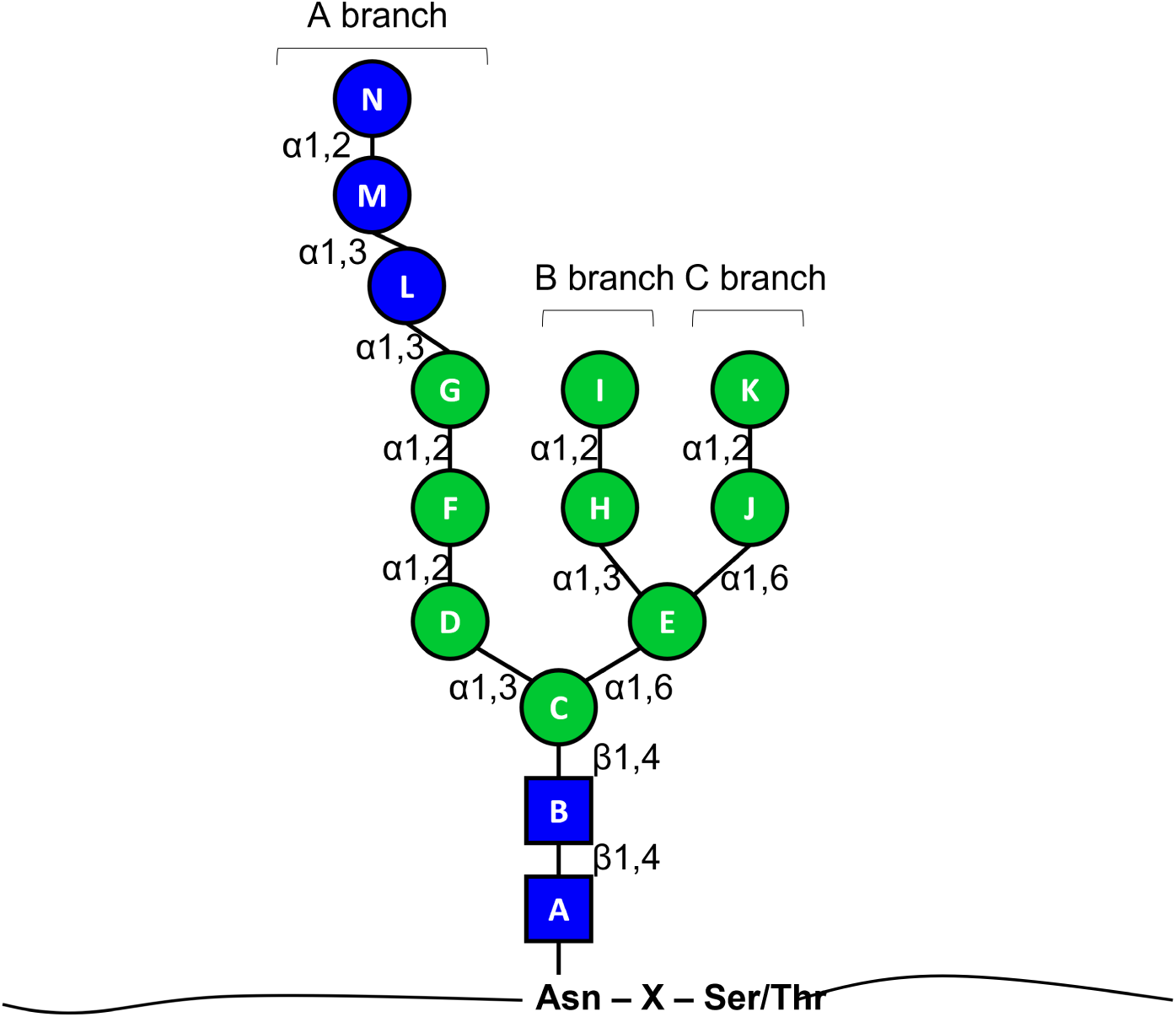
GlcNAc_2_Man_9_Glc_3_ glycan nomenclature. Saccharide moieties are depicted and colour-coded according to [131].

**Figure S2.**
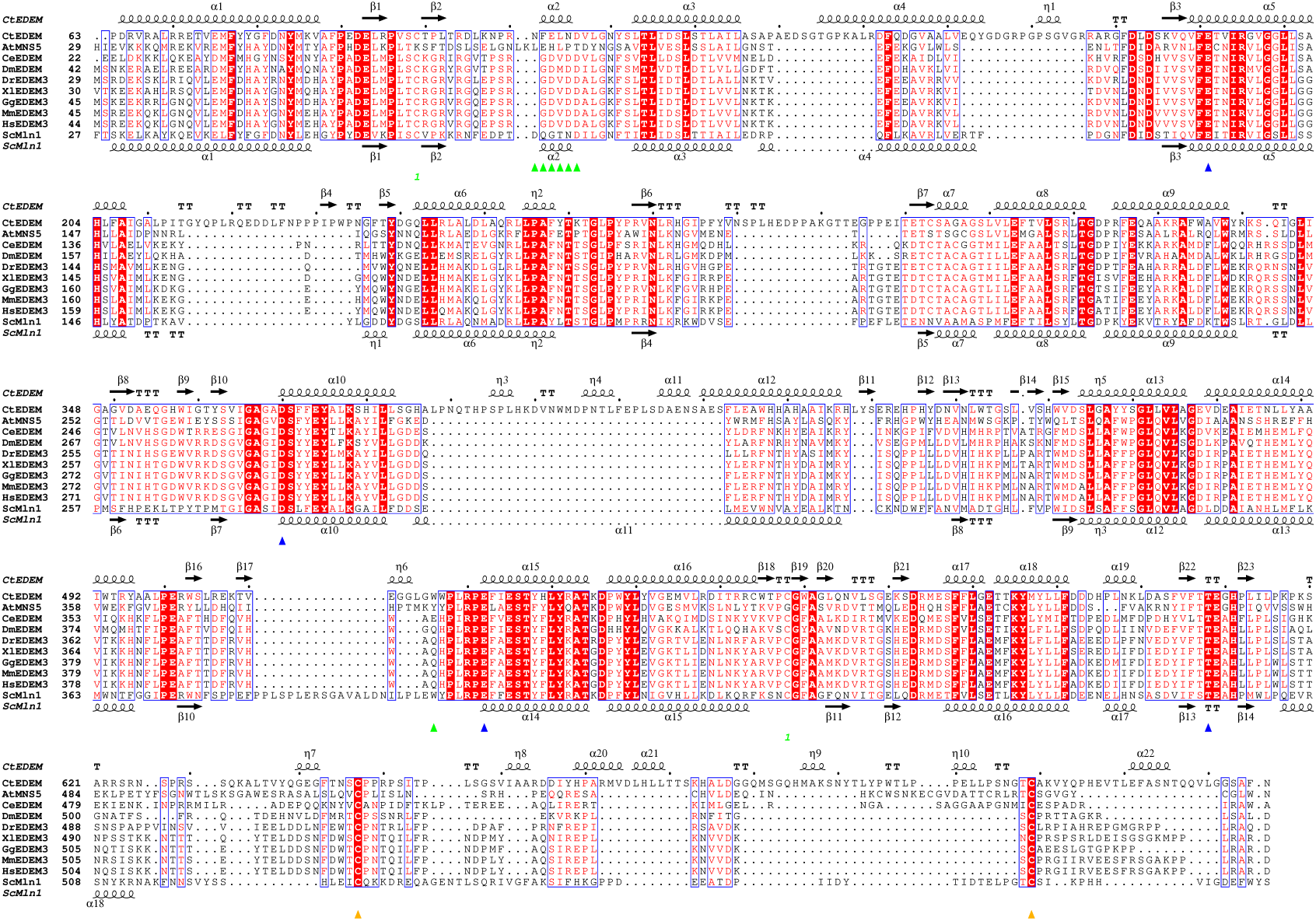
Sequence alignment of the catalytic domain (plus the linker region) of a few eukaryotic EDEM3 orthologues. UniProt codes: Ct: *Chaetomium thermophilum*, G0SCX7 · G0SCX7_THET3; Sc: *Saccharomyces cerevisiae*, P38888 · Mnl1_YEAST; At: *Arabidopsis thaliana*, Q9SXC9 · MNS5_ARATH; Ce: *Caenorhabditis elegans*, Q09641 · Q09641_CAEEL; Dm: *Drosophila melanogaster*, Q9VK27 · Q9VK27 DROME; Dr: *Danio rerio*, A0A8M2BEF7 · A0A8M2BEF7_DANRE; Xl: *Xenopus laevis*, Q6GQB9 · EDEM3_XENLA; Gg: *Gallus gallus*, A0A1D5PEI6 · A0A1D5PEI6_CHICK; Mm: *Mus musculus*, Q2HXL6 · EDEM3_MOUSE; Hs: *Homo sapiens*, Q9BZQ6 · EDEM3_HUMAN. Secondary structure elements from the Cryo-EM *Ct*EDEM_D_ :*Ct*PDI structure. The intramolecular bridge (conserved in each but *At*MNS5 sequence) is labelled 1 in green. The canonical GH47 acidic catalytic residues E191, D370, E522, and the conserved T608 coordinating a Ca^2+^ ion with its main chain carbonyl and O*_γ_*_1_ atom (see Supplementary Figure S13) are marked by blue triangles. The conserved Cys residues Cys A and Cys B which can form an intermolecular bond to the PDI are marked by orange triangles. The *α*2 helix and the Trp516 residue that direct branch C of the glycan into the active site are marked by green triangles. Figure made with the ESPript3.2 server at https://espript.ibcp.fr [132]. Alignment obtained with Clustal Omega [133].

**Figure S3.**
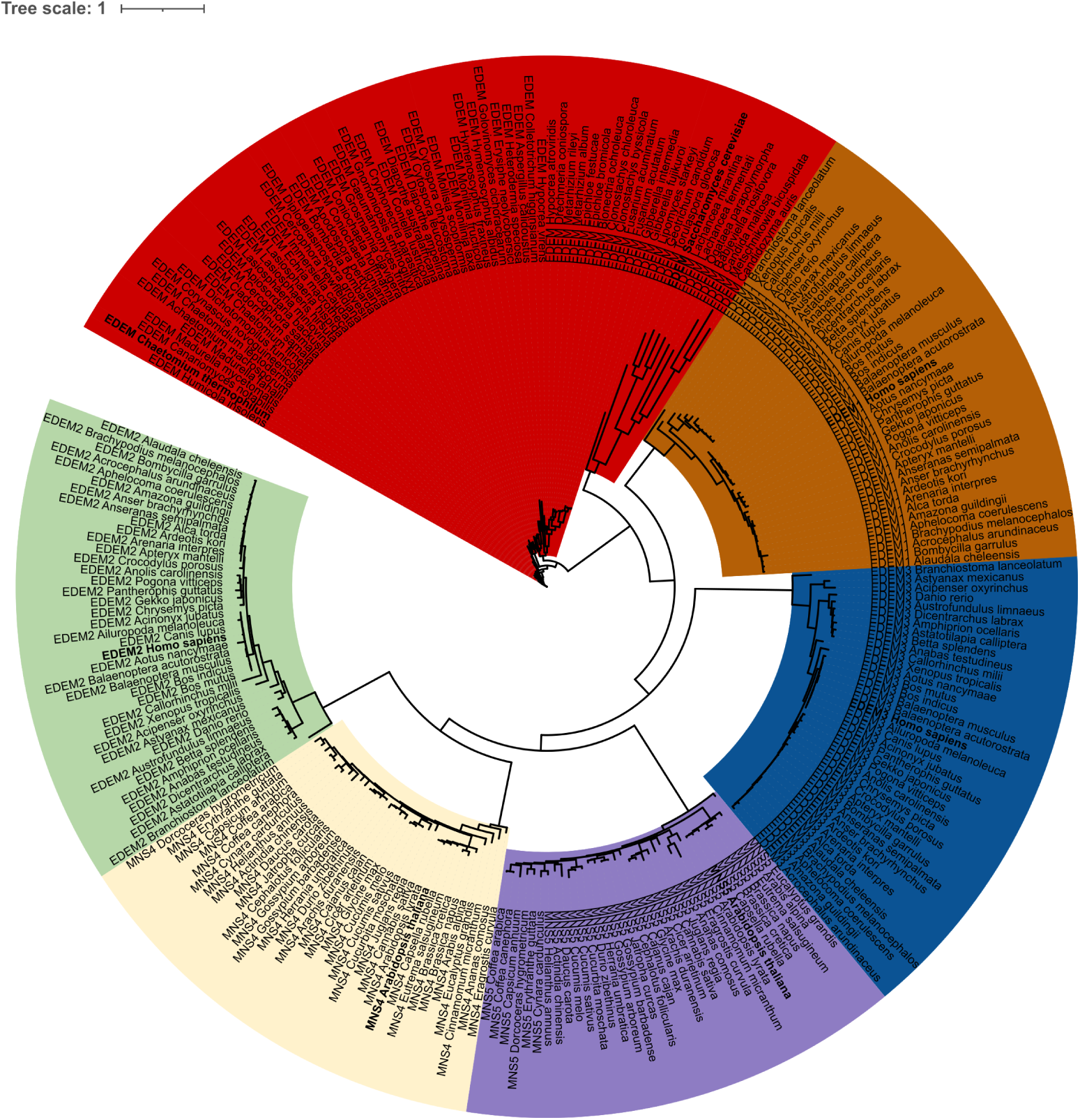
EDEM phylogeny. Six branches emerge from the analysis. Red branch: fungal EDEMs; these proteins feature an N-terminal membrane associated region, a GH47 domain followed by a linker region, and one or two distinct C-terminal domains. Dark orange branch: animal EDEM1s (isoform 1). The unstructured N-terminal region of animal EDEM1s (isoform 1) aligns well with the N-terminal section of fungal EDEMs - and have a ∼ 50 residues extension after the GH47 domain but no extra C-terminal domains. Blue branch: animal EDEM3s, featuring the linker region and one or two C-terminal domains - but lacking an obvious N-terminal membrane-associated region [56]; their PAD also contains the conserved, exposed disulfide bridge seen in the C-terminal domain of fungal EDEMs (*e.g. Hs*EDEM3 Cys693-Cys714). Violet branch: plant MNS5s: similar to mammalian EDEM3s but without any membrane-associated N-terminal helix. Yellow branch: plant MNS4s: they also feature a C-terminal extension following the GH47 domain. Green branch: EDEM2s, which have an N-terminal membrane-associated region [56].

**Figure S4.**
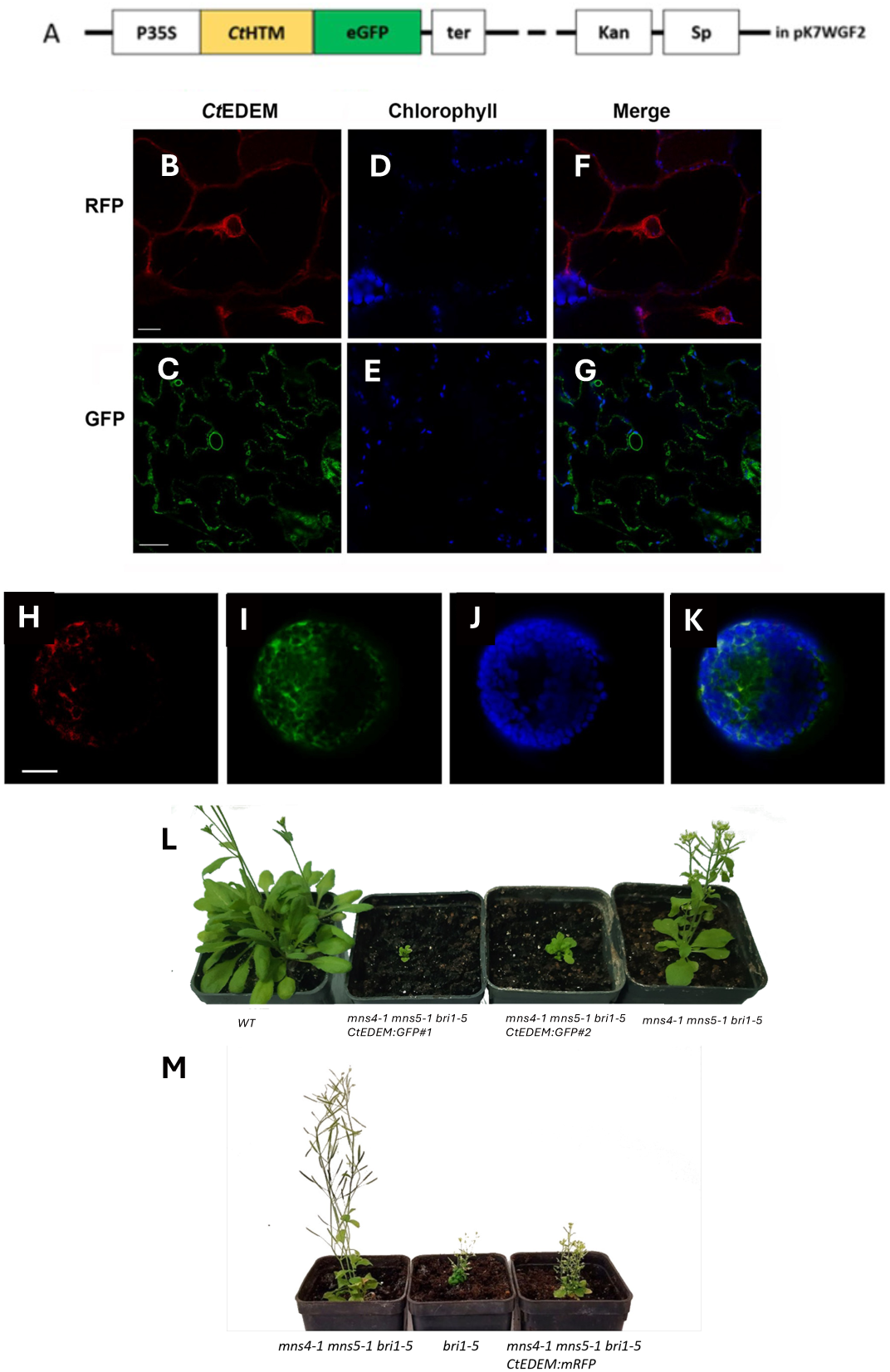
*Ct*EDEM has ERAD mannosidase activity *in planta*. **A:** Scheme of the insert portion of the *Ct*EDEM:GFP pK7WGF2 DNA plasmid used for the plant transformation. **B,C:** *Ct*EDEM, fused with mRFP or eGFP in *A. thaliana* seedlings; **D,E:** chlorophyll autofluorescence of the same seedlings; **F,G:** merge of the previous images; **H-K:** *Ct*EDEM, fused with mRFP localises to the ER in *A. thaliana* seedlings. Protoplasts obtained from leaves of *Ct*EDEM:mRFP stably transformed line were transformed for transient expression with the ER marker eGFP:KDEL. **H:** *Ct*EDEM:mRFP; **I:** eGFP:KDEL; **J:** chlorophyll autofluorescence; **K:** merge of all channels. Scale bar = 10*µ*m. **L**: Stable transformation of the *mns4-1/mns5-1/bri1-5* triple mutant (on the right) with *Ct*EDEM:eGFP (plants in the middle), compared to the WT *At* (on the left). The plants indicated as *Ct*EDEM-GFP#1 and *Ct*EDEM-GFP#2 are just two samples from different plant batches with identical genetic background and treated in the same way. **M**: Representative phenotypes of *A. thaliana* genotypes grown under the same condition. *mns4-1/mns5-1/bri1-5* triple mutant (on the left), *bri1-5* (in the middle), and the triple mutant complemented with *Ct*EDEM-mRFP (plant on the right).

**Figure S5.**
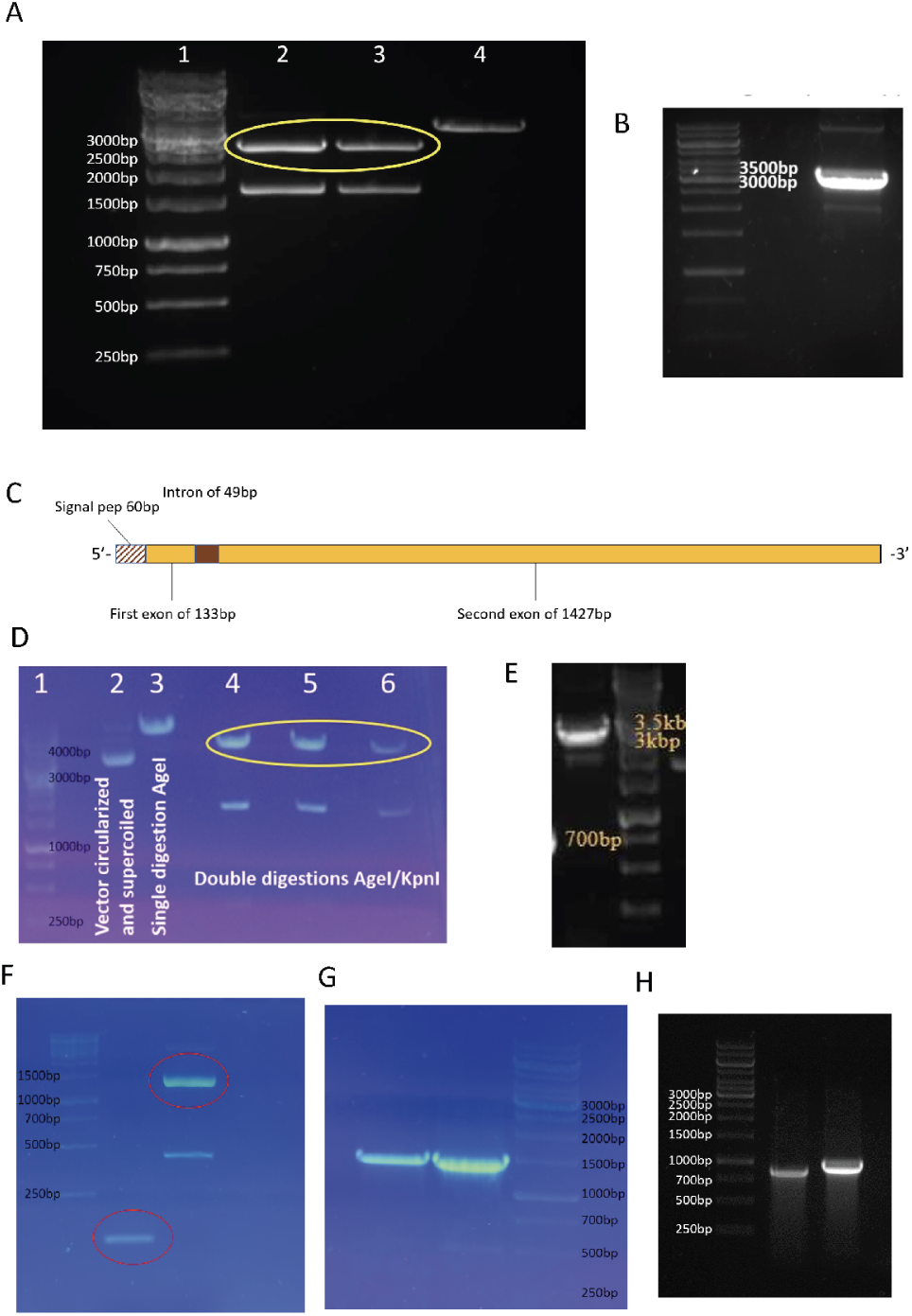
Cloning of *Ct*EDEM and *Ct*PDI. **A**: Linearisation of Litmus28i. **1**: Molecular Mass marker; Lanes 2 and 3: Litmus28i GBAN370S digested with the restriction enzymes EcoRI and XhoI (linearised Litmus28i circled in yellow); **4**: Litmus28i GBAN370S digested with the restriction enzyme EcoRI. **B**: *Ct*EDEM full length amplification. The gene was amplified by PCR creating the EcoRI/XhoI overlapping ends for the insertion in the linearised Litmus28i vector. **C**: Schematic sequence of *Ct*PDI gene as found in the *Ct* cDNA library. The *Ct*PDI gene is constituted by a 5’short signal peptide of 60bp (white with brown diagonal stripes), the first exon of 133bps (yellow), an intron of 49 bps (brown) and the second exon of 1427 bps (yellow). **D**: Linearisation of pHLsec. **1**: Molecular Mass marker; **2**: circular and supercoiled pEGFP SLC35B1-pHLsec; **3**: pEGFP SLC35B1-pHLsec digested with the restriction enzyme AgeI; **4,5,6**: pEGFP SLC35B1-pHLsec digested with the restriction enzymes AgeI and KpnI (linearised pHLsec circled in yellow). **E**: PCR amplification of GFP and *Ct*EDEM gene sequence. **D2**: amplification of *Ct*EDEM soluble domain sequence with the addition of overlapping ends, 5’overlaps GFP sequence and 3’overlaps pHLsec sequence. 1% agarose in 50mL TAE buffer, electrophoresis conditions: 100V for 1 hour. **F**: PCR amplification of *Ct*PDI two exons. **Left**: molecular-mass size marker; **middle**: amplification of the first exon (133bp); **right**: amplification of the second exon (1427bp). Agarose 2% in TAE buffer. Electrophoresis conditions: 100V for 1 hour. **G** Colony-PCR of *E. coli* colonies transformed with *Ct*PDI pHLsec vector. The success of the transformation with the *Ct*PDI pHLsec vector was investigated in two different colonies through colony-PCR. The PCR-amplified bands show DNA segments of the expected size (1560bp). **H** Colony-PCR of *Agrobacterium tumefaciens* colonies transformed with the *Ct*EDEM pK7FWG2 vector. The success of the transformation with the *Ct*EDEM pK7FWG2 vector was investigated with two different sets of primers through colony-PCR. The PCR-amplified bands show DNA segments of the expected size: the band on the left is 900 bp and the band on the right is 986 bp.

**Figure S6.**
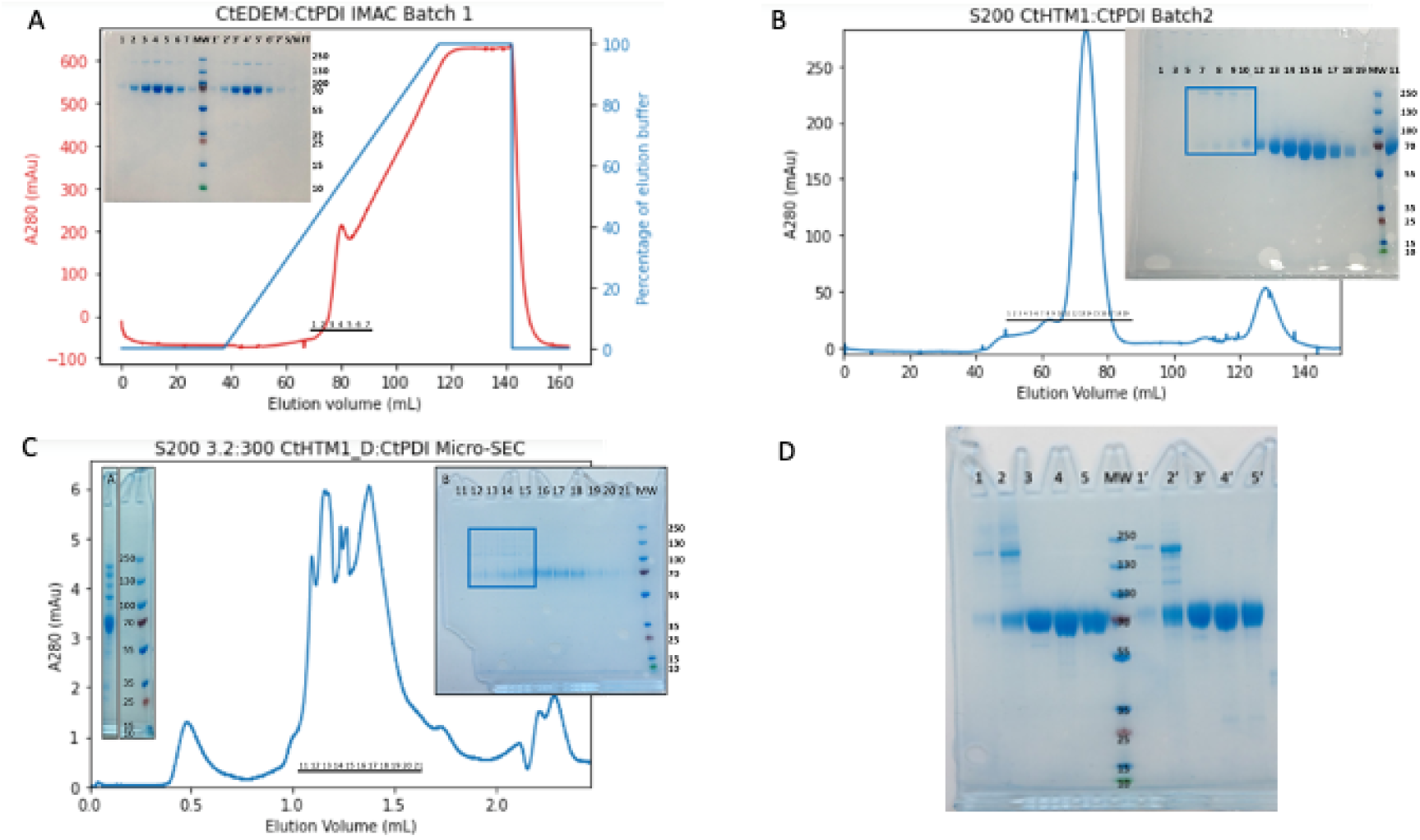
Expression and purification of *Ct*EDEM_D_ :*Ct*PDI. **A**: Elution profile of the IMAC purification of the first batch of *Ct*EDEM_D_ :*Ct*PDI. The blue curve describes the gradient of elution buffer (IMACE). The red curve describes the UV absorbance at *λ*=280 nm. The elution buffer absorbs at *λ*=280 nm, likely because of metal impurities forming a ferrocene complex with the imidazole. Eluted protein, contained in fractions from 1 to 6, is visible as a peak on top of the linear imidazole gradient. In the upper left panel is shown the SDS-PAGE 4-12% Bis-Tris gel of the fractions 1-6, indicated in the chromatogram. **Left**: Lane 1 to 7 show samples from eluted fraction 1 to 7, respectively; **Centre**: MM marker; **Right**: lanes labelled with a prime contained samples from eluted fraction 1 to 7 treated with reducing agent. Gel run at 200V for 55’ in MOPS buffer. **B**: Elution profile of the S200 16/60 SEC purification of Batch 2 of *Ct*EDEM_D_ :*Ct*PDI. The blue line describes the UV absorbance at *λ*=280 nm. Eluted proteins, contained in fractions from 1 to 19, are visible as a peak. In the upper right panel is shown the SDS-PAGE 4-12% Bis-Tris gel of some of the fractions indicated in the chromatogram. The boxed fractions contain the *Ct*EDEM_D_ :*Ct*PDI heterodimer. Gel run at 200V for 55’ in MOPS buffer. **C**: Elution profile of the S200 3.2:300 Micro-SEC purification of the pooled Batches 1 and 2 of *Ct*EDEM_D_ :*Ct*PDI. The blue line describes the UV absorbance at *λ*=280 nm. The numbers on the black bar indicate the elution fraction numbers. **Inset A**: a 4-12% SDS-PAGE lane of the Batch 2 *Ct*EDEM_D_ :*Ct*PDI complex sample at OD_280_=0.741 after 10 days at 4 *^◦^*C (gel run in MOPS buffer at 200 V for 55’). Proteolysis of *Ct*EDEM_D_ is visible as three additional bands at MM ≈ 135, 120 and 115 kDa. **Inset B**: a 4-12% SDS-PAGE gel of elution fractions 11-21 of the S200 3.2:300 Micro-SEC purification of the pooled Batches 1 and 2 of *Ct*EDEM_D_ :*Ct*PDI (gel run in MOPS buffer at 200 V for 55’). Each sample was prepared with 15 *µ*L of S200 3.2:300 Micro-SEC fraction plus 5 *µ*L of non-denaturing 4x SDS-PAGE loading dye, and denatured for 1’at 90 *^◦^*C prior to loading on the gel. Boxed fractions 12 to 15 appear to contain the 70 kDa *Ct*PDI and a number of proteolytic fragments derived from *Ct*EDEM_D_ (the lightest of which likely corresponds to the loss of GFP, ≈ 115 kDa, see Inset A). Fraction 13 was used for Cryo-EM on a graphene-oxide grid. **D**: SDS-PAGE 4-12% Bis-Tris gel of the pooled and concentrated S200 16/60 SEC fractions of the *Ct*EDEM_D_ :*Ct*PDI heterodimer and the *Ct*PDI. Gel run at 200 V for 55’in MOPS buffer. Lanes with primed numbers contain reducing agent. Lanes 1,1’: 2 *µ*L of *Ct*EDEM_D_ :*Ct*PDI complex Batch 2, OD_280_=0.265; lanes 2,2’: 4 *µ*L of *Ct*EDEM_D_ :*Ct*PDI complex Batch 2, OD_280_=0.741; lanes 3,3’: 4 *µ*L of *Ct*PDI Batch 1, OD_280_=0.877; lanes 4,4’: 3 *µ*L of *Ct*PDI Batch 2, OD_280_=1.218; lanes 5,5’: 1 *µ*L of *Ct*PDI Batch 2, OD_280_=3.082. Lane 6: MM ladder with molecular mass in kDalton.

**Figure S7.**
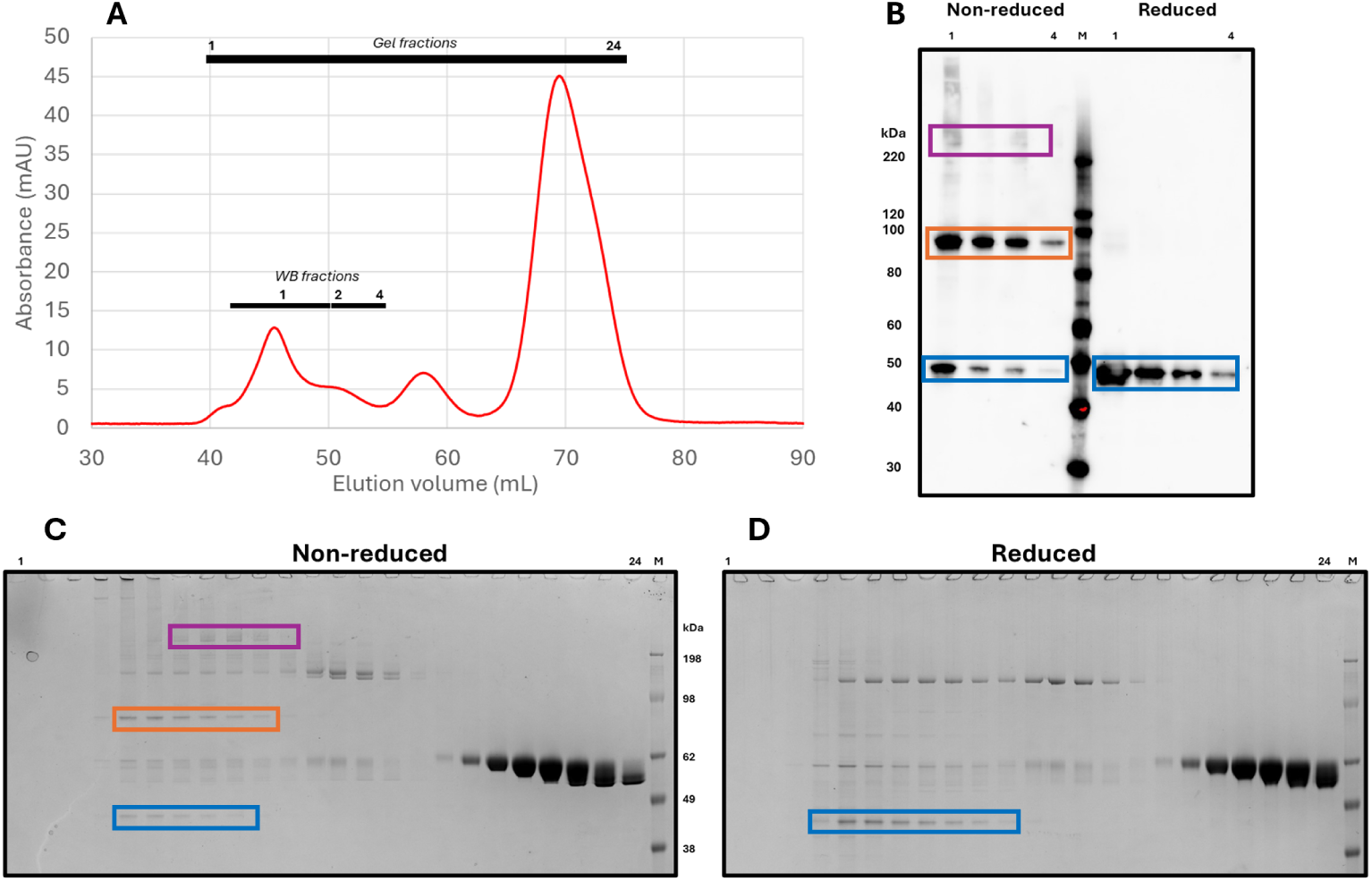
Purification of a *Ct*EDEM:PDI:A1AT-NHK ternary complex. The SEC experiment was carried out on a Superdex 200 16/60 pg column after an initial IMAC step. The fractions that were taken forward for SDS-PAGE and Western blotting (WB) are highlighted with the black line. The SDS-PAGE and WB fractions are labelled to match the gel and WB respectively. Protein molecular mass marker standards are indicated (M) and with sizes specified adjacently. **A**. Size exclusion chromatography (SEC) trace for a *Ct*EDEM:PDI:A1AT-NHK ternary complex purified from the cell supernatant. **B**. Western blot of SEC chromatogram fractions using an anti *α*-1 antitrypsin antibody. Fractions 1 and 4 are labelled with fraction 2-3 in between. The left-hand side lanes are in non-reducing conditions, the right-hand side lanes are in reducing conditions. **C. D.** SDS-PAGE gel of the media *Ct*EDEM:PDI:A1AT-NHK SEC sample in non-reducing (C) and reducing (D) conditions. Fraction 1 and fraction 24 are labelled with fractions 2-23 in between. The blue box highlights A1AT-NHK, the orange box highlights a likely *Ct*PDI:A1AT-NHK heterodimer, the purple box high-lights a likely *Ct*EDEM:PDI:A1AT-NHK ternary complex.

**Figure S8.**
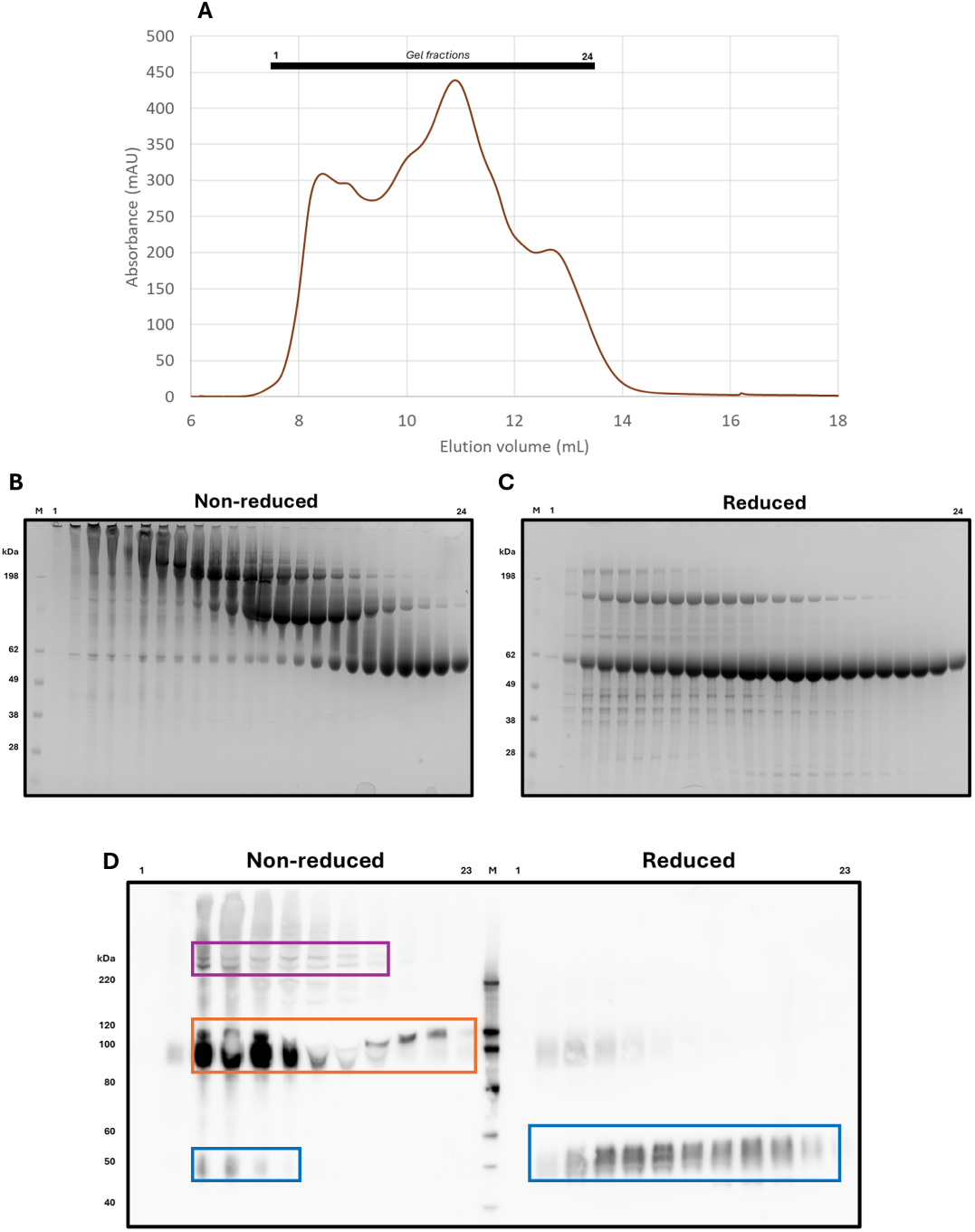
Purification of a *Ct*EDEM:PDI Cys53Ser:A1AT-NHK ternary complex. SEC of *Ct*EDEM:PDI Cys53Ser:A1AT-NHK protein purified from the cell supernatant. The SEC experiment was carried out on a Superdex 200 10/300 Increase GL column after an initial IMAC step. The fractions that were taken forward for SDS-PAGE and Western blotting (WB) are highlighted with the black line. The SDS-PAGE and WB fractions are labelled to match the gel and WB respectively. Protein molecular mass marker standards are indicated (M) and with sizes specified adjacently. **A**. SEC trace. **B,C.** SDS-PAGE gel of the supernatant Cys53Ser PDI trapping mutant *Ct*EDEM:PDI Cys53Ser:A1AT-NHK SEC sample in non-reducing (B) and reducing (C) conditions. Fraction 1 and fraction 24 are labelled with fractions 2-23 in between. **D**. Western blot of SEC fractions using an anti *α*-1 antitrypsin antibody. Fractions 1 and 23 are labelled with odd fractions 3-21 in between. The left-hand lanes are in non-reducing conditions, the right-hand lanes are in reducing conditions. The blue box highlights A1AT-NHK, the orange box high-lights a putative *Ct*PDI Cys53Ser:A1AT-NHK heterodimer, the purple box highlights a putative *Ct*EDEM:PDI Cys53Ser:A1AT-NHK ternary complex.

**Figure S9.**
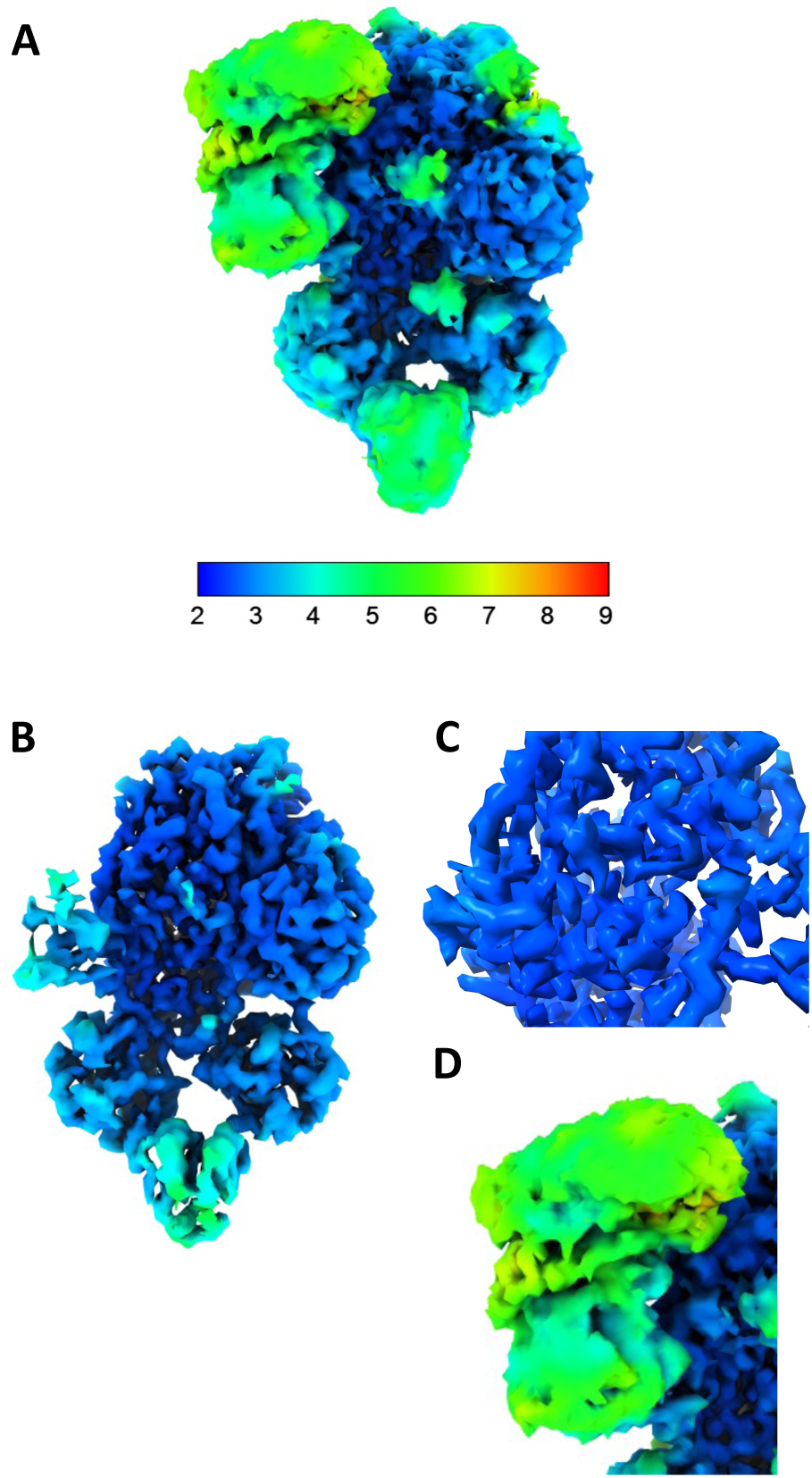
Local resolution of the EM map of the *Ct*EDEM:PDI heterodimer. EM map of *Ct*PDI:*Ct*EDEM at different display thresholds with local map resolution represented in surface colour. **A.** Low threshold map displaying areas of density in the *Ct*PDI and *Ct*EDEM map. The scale bar encodes map resolution. Dark blue means 2.0 Å and red 9.0 or lower Å local resolution. **B.** Higher threshold map focusing on the higher resolution core and immediate peripheral domains. **C.** Close in view of the highest resolution part of the map, at the centre of the GH47 catalytic domain. **D.** Close-in view of the lowest resolution part of the map, displayed at the lowest threshold. These maps were visualised in ChimeraX [57].

**Figure S10.**
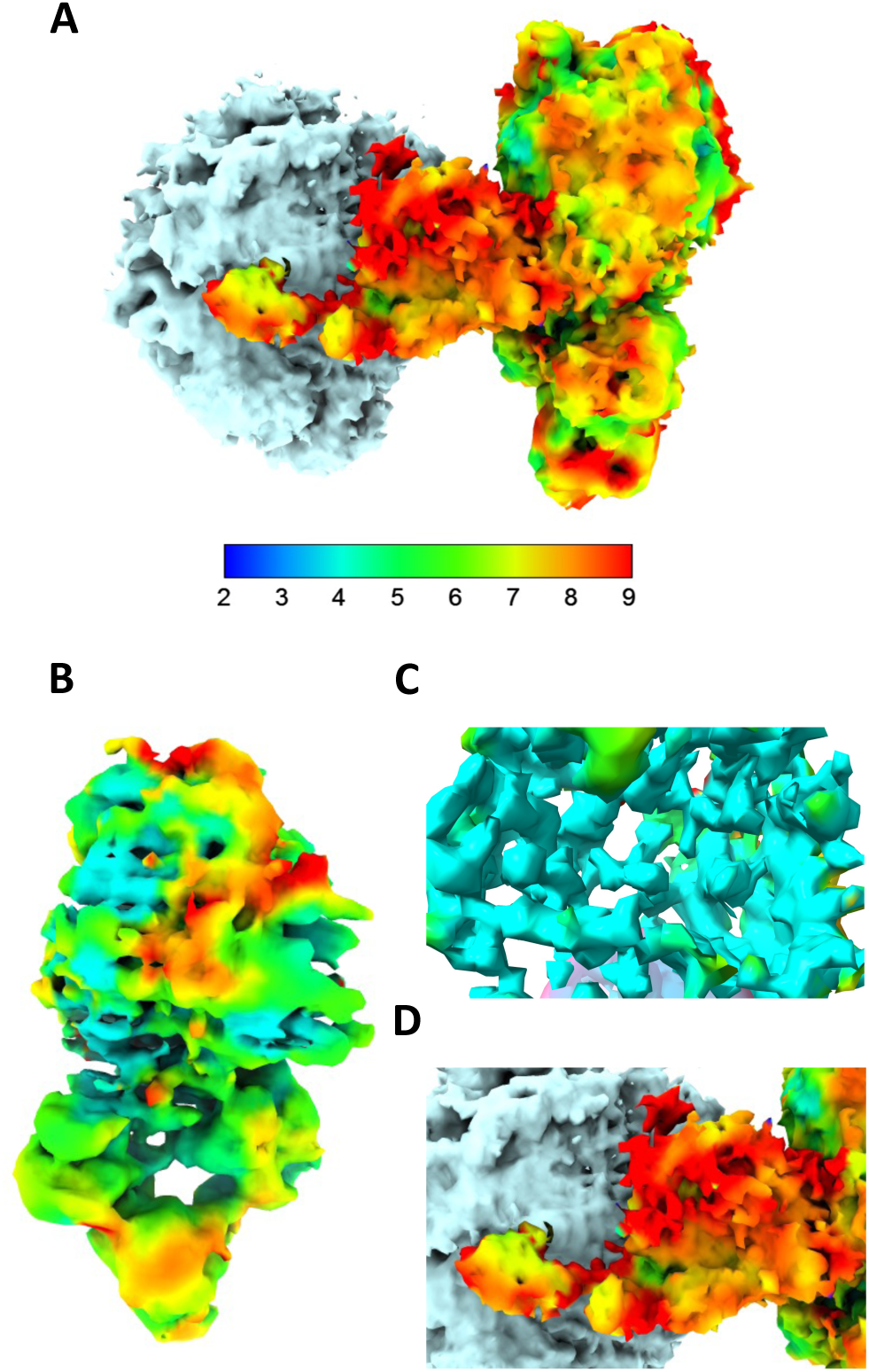
Local resolution of the EM map of the *Ct*EDEM:PDI A1AT-NHK complex. EM map of *Ct*EDEM:PDI A1AT-NHK complex at different display thresholds with local map resolution represented in surface colour. **A.** Low threshold map displaying areas of density in the *Ct*PDI and *Ct*EDEM map when bound to A1AT-NHK (the latter density shown in grey on the left). The scale bar encodes map resolution. Dark blue means 2.0 Å and red 9.0 or lower Å local resolution. **B.** Higher threshold map focusing on the higher resolution core and immediate peripheral domains. **C.** Close-in view of the highest resolution part of the map, at the centre of the GH47 catalytic domain. **D.** Close-in view of the lowest resolution section of the map displayed at the lowest threshold. These maps were visualised in ChimeraX [57].

**Figure S11.**
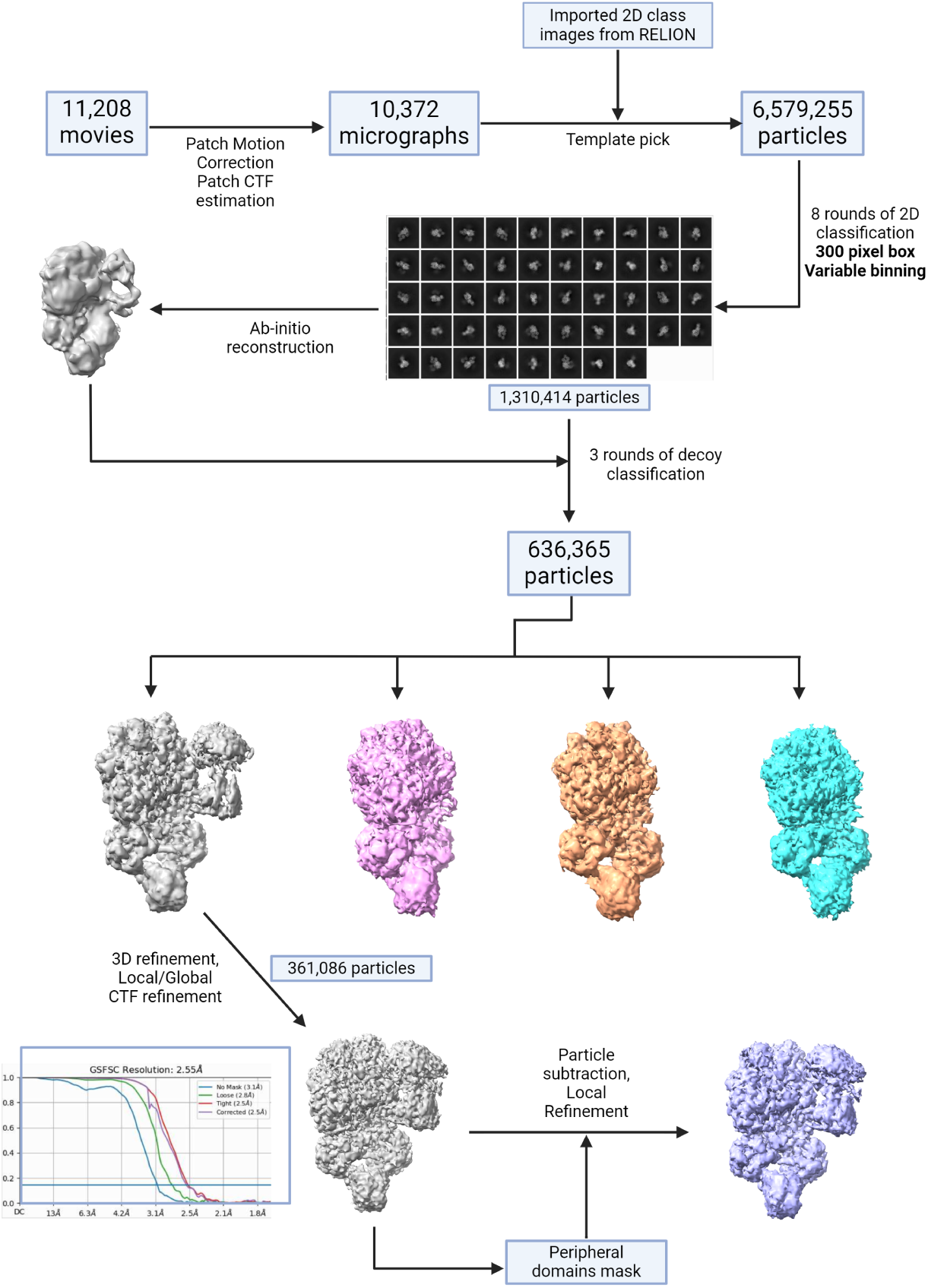
Cryo-EM processing workflow for the *Ct*EDEM:PDI ERAD checkpoint structure. Movies collected on the Titan Krios were motion corrected and CTF parameters estimated. Exposures were curated to remove those with poor CtfAstigmatism or those which were outliers in ice thickness. The 2D class average images from RELION were imported and used as templates to pick particles from the micrographs using the cryoSPARC template picker. The total number of particles (6,579,255) were cut down to leave only good particles (1,310,414) by eight rounds of 2D classification at variable binning. The particles were used to generate an initial 3D model, which was then used as a reference model for decoy classification. Three rounds of decoy classification were carried out to further select only the best particles. After decoy classification, a single round of 3D classification was run, with one class showing the best density for the peripheral domains taken forward for 3D refinement. 3D refinement with local and global CTF refinement included produced an EM map at 2.55 Å. Particle subtraction to remove all density except for the peripheral domains, followed by masked local refinement improved the density for the peripheral domains.

**Figure S12.**
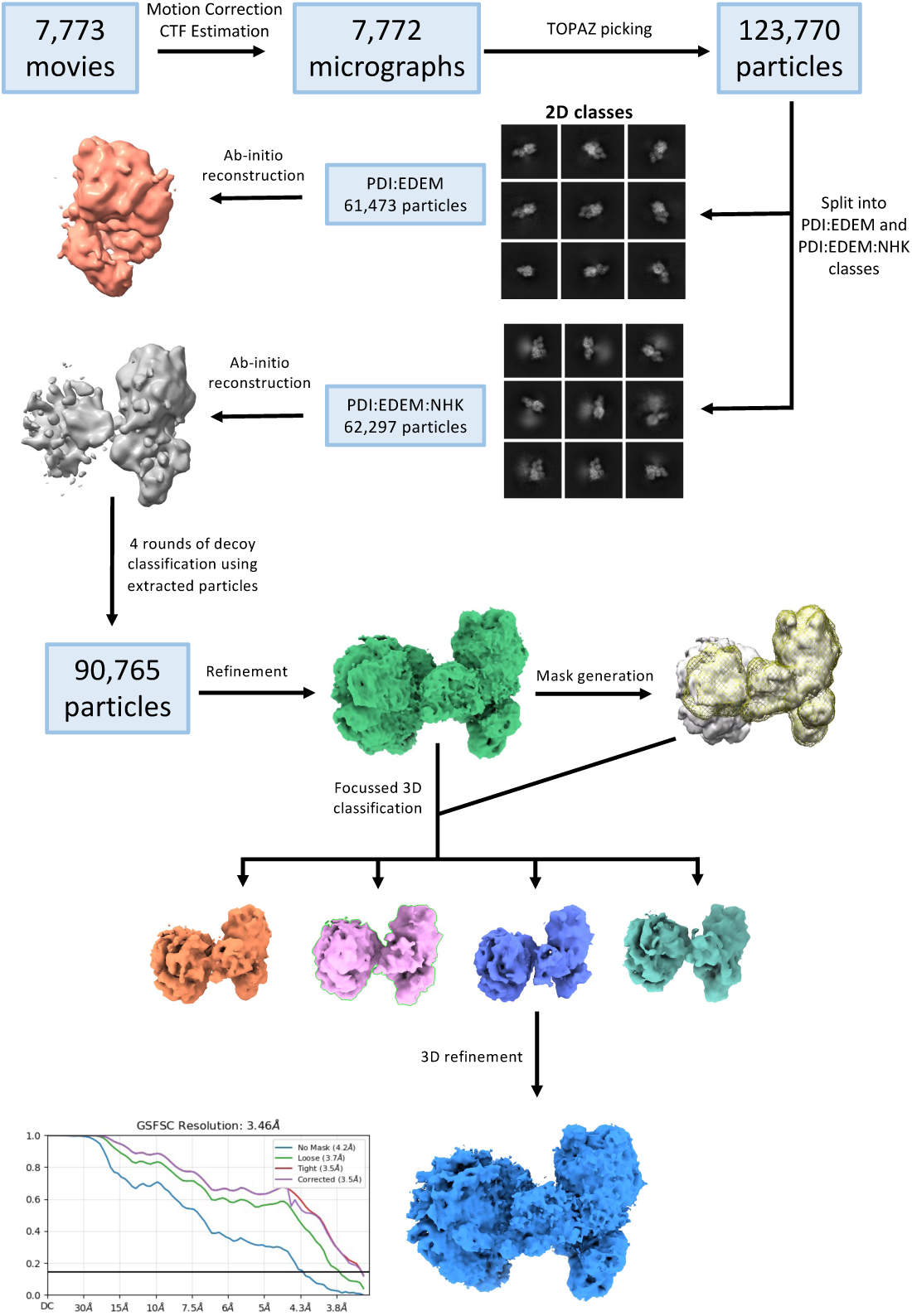
Flowchart of the RELION 4.0 processing steps for the *Ct*EDEM:PDI:A1AT-NHK Cryo-EM dataset. Movies collected on the Titan Krios were motion corrected and CTF parameters estimated. The in-built Topaz autopicker was used to pick particles which were extracted with a box size of 360 pixels. 4 rounds of 2D classification were carried out to remove junk particles after which particles were clearly split into two sets: those which resembled the PDI:EDEM heterodimer classes and those with extra density, likely PDI:EDEM:A1AT-NHK. Both sets of particles were used to generate an initial 3D model of apo (red) and NHK-bound *Ct*EDEM:PDI complex (grey). Both *ab-initio* models were used along with 3 junk particle models for decoy 3D classification using all the particles extracted, and particles were sorted into their respective classes. Classes containing the *Ct*EDEM:PDI:A1AT-NHK ternary complex were refined, and a mask generated around the core *Ct*EDEM:PDI portion (yellow mesh). Focussed 3D classification was done using this mask, and the best class was refined to produce the final structure

**Figure S13.**
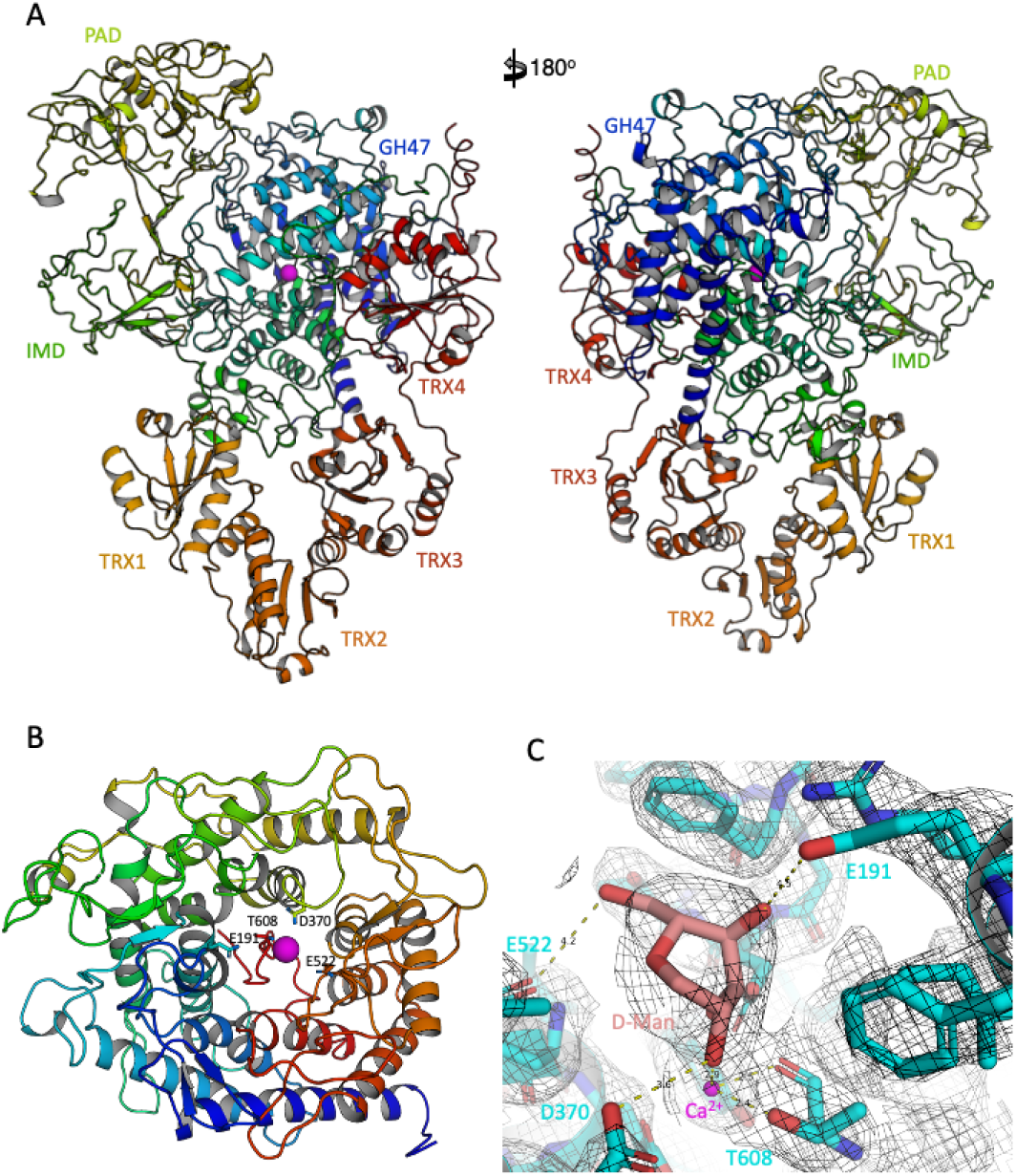
Details of the apo *Ct*EDEM:*Ct*PDI structure. **A:** two views of the apo *Ct*EDEM:*Ct*PDI structure, in cartoon representation and coloured blue-to-red from N- to C-terminus. The catalytic Ca^2+^ ion is a magenta sphere; **B:** a cartoon representation of the *Ct*EDEM GH47 domain, centred around its active site. At the centre of the image, the hairpin that plugs the bottom of the catalytic pocket is coloured red. The canonical GH47 acidic catalytic residues E191, D370, E522, and the conserved T608 coordinating a Ca^2+^ ion with its main chain carbonyl and O*_γ_*_1_ atom (see blue triangles in Supplementary Figure S2). *Ct*EDEM E191 is the residue corresponding to *Mm*EDEM3E147, *Hs*EDEM1E225, *Mm*EDEM1 E220 and *Hs*EDEM2 E117 each of which when mutated to Q abrogate mannosidase activity respectively [18, 43, 46, 134]. *Ct*EDEM D370 is the residue corresponding to *Sc*Mnl1 D279 which when mutated to N abrogates mannosidase activity [17]. *Ct*EDEM E522 is the residue corresponding to *At*MNS4 E376 and *At*MNS5 E388 which when mutated to N abrogate mannosidase activity [51]. The octahedral coordination of this Ca^2+^ ion is usually completed by six water molecules [61, 62], and substrate binding usually displaces two of the water molecules through coordination with the O2’and O3’hydroxyls of the terminal *α*1,2-mannose residue within the catalytic −1 sub-site. **C:** The *D*-Mannose residue trapped in the apo structure active site, built with the pose and coordination geometry observed in homologous structures, such as the one of ER Man IB co-crystallised with the *D*-Mannose analogues 1-deoxymannojirimycin and kifunensine (PDB IDs 1FO2 and 1FO3) [61] and with a thio-disaccharide substrate analogue (PDB ID 5KK7) [76]. The catalytic Ca^2+^ ion is a magenta sphere. The map was contoured at 1.5 *σ* level (grey mesh). Images made in PyMol.

**Figure S14.**
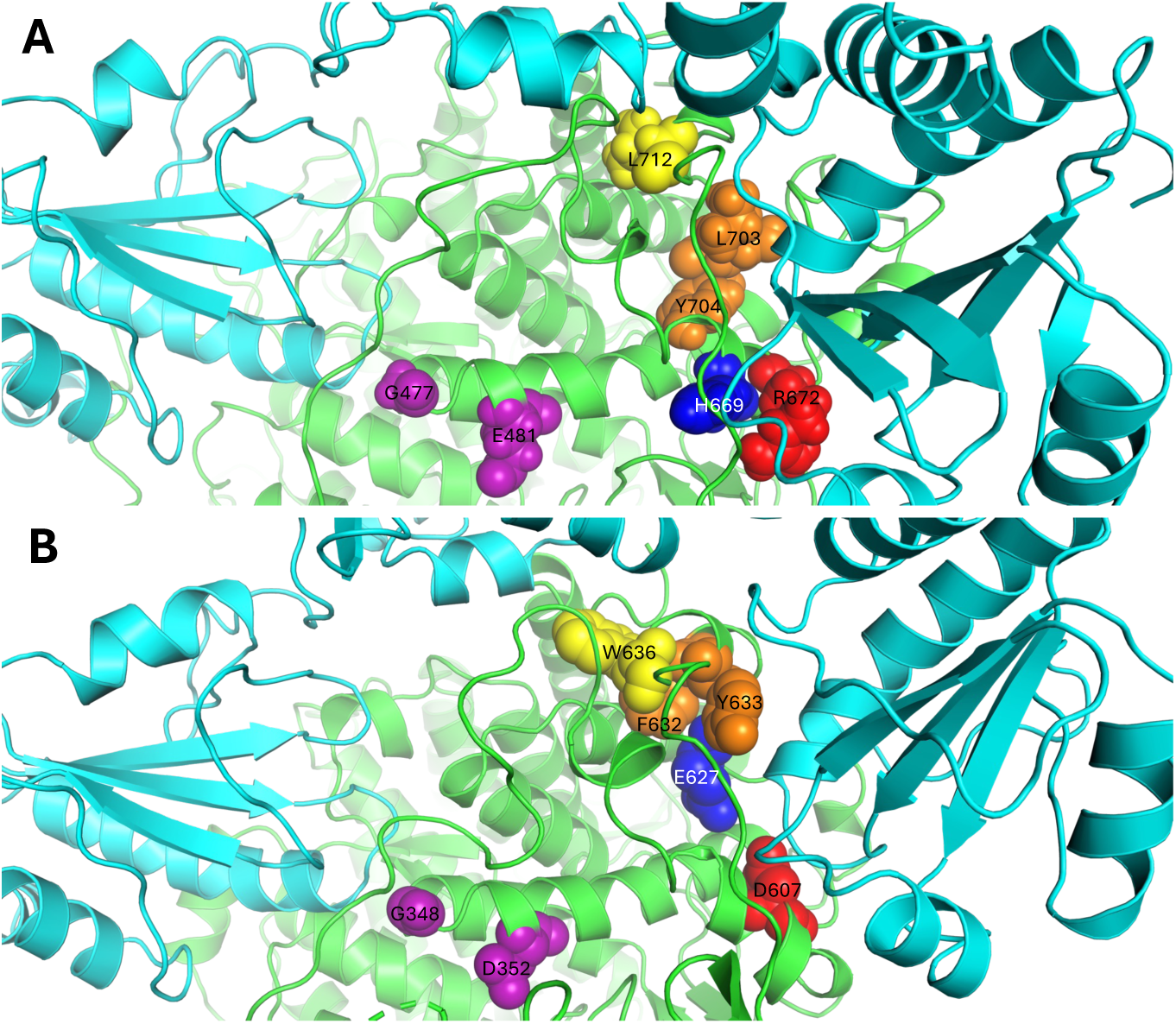
EDEM missense mutations that abrogate function and localise close to the EDEM:PDI interface. *Ct*EDEM:*Ct*PDI and *Sc*Mnl1:*Sc*PDI1 structures are displayed with EDEMs as green cartoons and PDIs as cyan cartoons. *Sc*Mnl1 mutants D607A, E627A, W636A, F632L and Y633S abrogate the association with *Sc*PDI1 [34, 135]; the G343E and A347V mutants of the *Arabidopsis thaliana, (At)* plant EDEM orthologue MNS5 abrogate ERAD mannosidase function [48, 136]. The mutations map close to the interface between EDEM and PDI in the *Ct*EDEM:*Ct*PDI and *Sc*Mnl1:*Sc*PDI1 structures. **A:** equivalent residues in *Ct*EDEM: R672 (red), H669 (blue), L712 (yellow), L703 and Y704 (orange), G477 and E481 (purple), shown as spheres; **B:** *Sc*Mnl1: D607 (red), E627 (blue), W636 (yellow), F632 and Y633 (orange), and the residues equivalent to *At* MNS5 G343 and A347, *i.e. Sc*Mnl1 G348 and D352 (purple), are shown as spheres. Mutants *At*MNS5 G424R and A447V also impair activity [48] but do not localise to the PDI interface and rather seem to disrupt the GH47 fold (not shown). The loss of function associated with the deletion of the 4 C-terminal residues of *Sc*Mnl1 [34] is also explained by the damage inflicted to the fold of the IMD domain. Image made in PyMol.

**Figure S15.**
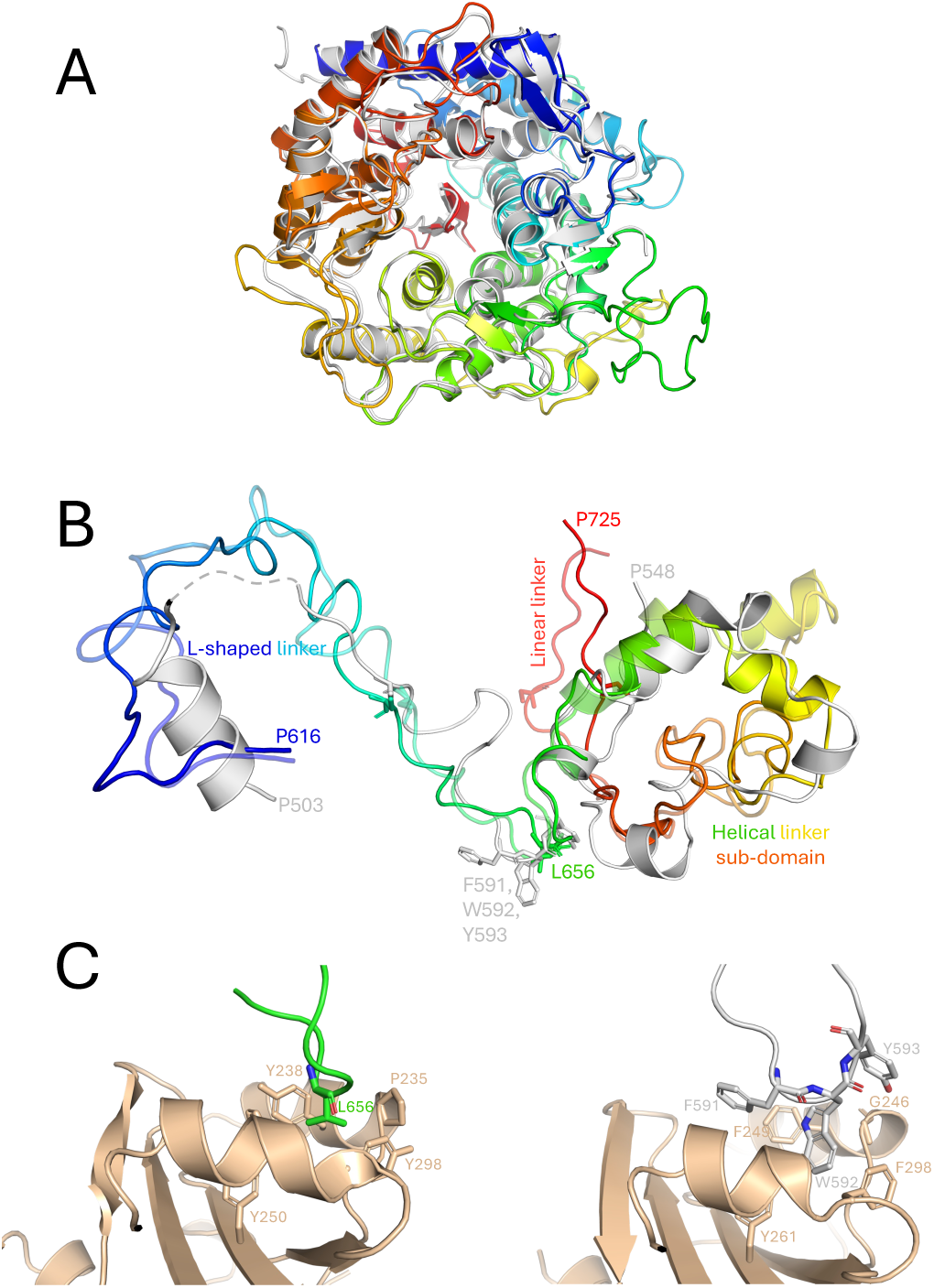
Structural comparison of *Ct*EDEM:PDI and *Sc*Mnl1:*Sc*PDI1. **A**: overall GH47 domains superposition. *Sc*Mnl1-GH47 (PDB ID 8ZPW) in white cartoon representation; *Ct*EDEM-GH47 (apo structure, PDB ID 9TPA) in cartoon representation, coloured blue to red from N- to C-terminus. At the centre of the image, the hairpin that plugs the bottom of the catalytic pocket is coloured red. In both *Ct*EDEM:PDI structures, the GH47 catalytic domain is very similar to the *Sc*Mnl1 one in PDB ID 8ZPW: the overall *Sc*Mnl1_GH47_:*Ct*EDEM_GH47_ RMSD_C_*_α_* over 429 residues is 1.03 or 1.36 Å respectively in the apo or A1AT-NHK bound forms. **B**: comparison of linkers regions. The *Sc*Mnl1 linkers (residues 503-648, with the missing portion 516-574 as a dashed line) are in white cartoon representation; *Ct*EDEM-GH47 (apo structure in full colours and A1AT-NHK complex in faded colours) in cartoon representation, coloured blue to red from N- to C-terminus. The residues where the linker associates with the PDI TRX3 domain are in stick representation (see following panel for details); **C**: Left-hand side panel: the exposed *Ct*EDEM Leu656 (where the L-shaped linker joins the helical linker subdomain) forms an intermolecular hydrophobic contact with the *Ct*PDI surface pocket formed by Tyr238, Tyr250 and His290 (PDB ID 9TPA), equivalently to what is observed in the *Sc*Mnl1:*Sc*PDI1 orthologous complex structure (right-hand side panel), in which *Sc*Mnl1 Trp592 and Tyr593 interact with *Sc*PDI1 Phe249, Tyr261 and His301 (PDB ID 8ZPW, [31]). *Ct*EDEM: residues 653-660 in green C carbon cartoon, Leu656 in green sticks. *Sc*Mnl1: residues 586-595 in white cartoon, Phe591, in white sticks. The *Ct*PDI and *Sc*PDI1 TRX3 domains are in wheat cartoon, with residues forming the surface hydrophobic pocket represented in sticks.

**Figure S16.**
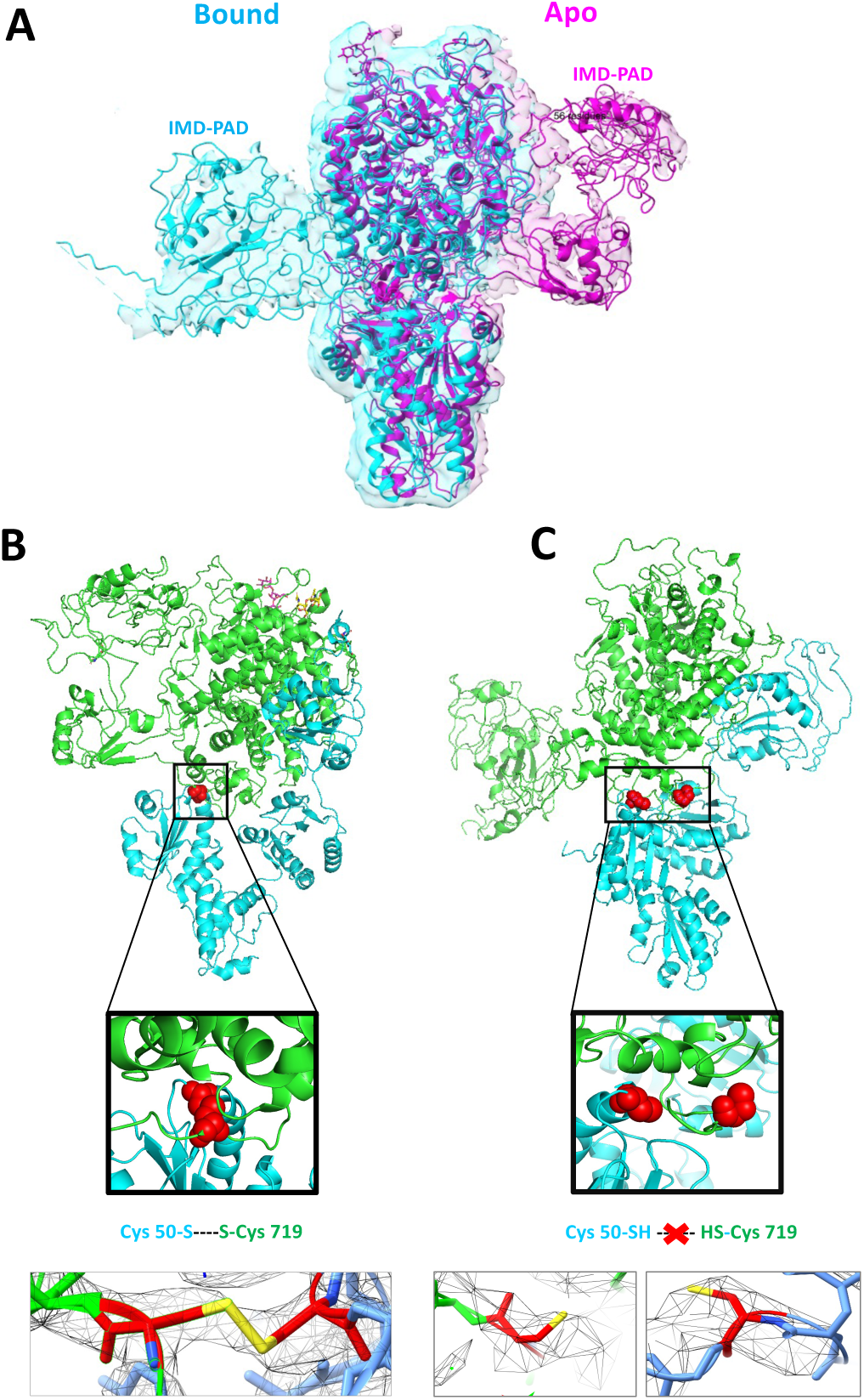
Comparison of the apo and A1AT-NHK-bound *Ct*EDEM:*Ct*PDI structures. **A**: superimposition of the bound (blue) and apo (magenta) structures and maps after aligning features in the core GH47 catalytic region of *Ct*EDEM (Leu 540 – Arg 553, Arg 325 – Arg 340). **B,C**: *Ct*PDI (blue) and *Ct*EDEM (green) for both the apo (panel **B**) and A1AT-NHK-bound (panel **C**) structures. Red spheres indicate the *Ct*PDI Cys 50 and *Ct*EDEM Cys 719, which form an inter-molecular disulphide in absence (**B**) but not in presence (**C**) of the A1AT-NHK substrate. Cysteine side chains and map contoured at 0.145 are shown below each panel.

**Figure S17.**
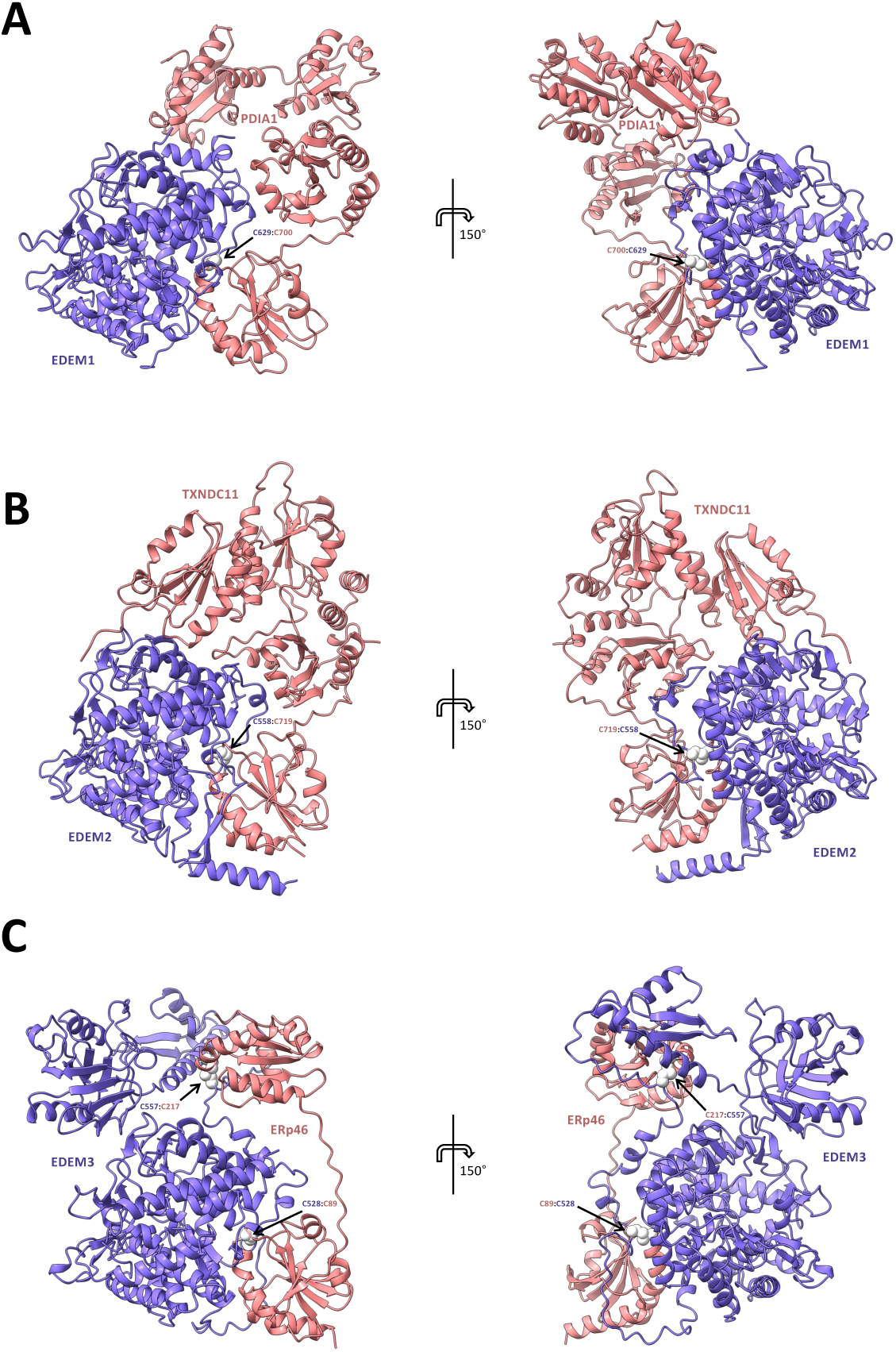
AlphaFold3 models of the three human EDEM:PDI complexes. The hu-man EDEM is in purple and its partner PDI in salmon cartoon representation. The left-hand and right-hand side views are related by a rotation of 150 degrees around the vertical axis. **A**: *Hs*EDEM1, UniProt Q92611, residues 112-596 and 626-657, *Hs*P4HB *aka* PDIA1: UniProt P07237, residues 21-476. The disulfide bond between *Hs*EDEM1 Cys629 and *Hs*PDIA1 Cys397 is in white spheres. **B**: *Hs*EDEM2:*Hs*TXNDC11: *Hs*EDEM2, UniProt Q9BV94, residues 29-578; *Hs*TXNDC11, UniProt Q6PKC3 residues 91-265, 292-422, 594-799. The disulfide bond between *Hs*EDEM2 Cys558 and *Hs*TXNDC11 Cys719 is in white spheres. **C**:*Hs*EDEM3:*Hs*ERp46_TRX1-2_: *Hs*EDEM3, UniProt Q9BZQ6, residues 43-796; *Hs*ERp46, domains TRX1 and TRX2, UniProt Q8NBS9, residues 60-298; In the EDEM3:ERp46_TRX1-2_ model, the SS1 intermolecular disulfide between Cys217 in the TRX2 domain of *Hs*ERp46and *Hs*EDEM3 Cys528 [19] (“Cys A”) and the disulphide bridge between *Hs*EDEM3 Cys557 (“Cys B”) and *Hs*ERp46 Cys89 are in white spheres. AlphaFold3 model statis-tics in the Methods section. Figure made with Ch_6_i_7_meraX [57].

### EDEM phylogeny suggests a later divergence of the EDEM2/MNS4 lineage

To elucidate the evolutionary relationships between EDEM proteins [28], we deployed structure-based and sequence-based phylogeny [137, 138], analyzing fungal eukaryotic EDEMs, plant MNS4s and MNS5s, and mammalian EDEMs. Supplementary Figure S3 illustrates the EDEMs phylogenetic tree [137, 138]. Animal EDEM1s and EDEM3s (orange and blue branches in Supplementary Figures S3) appear to be the closest orthologues to fungal EDEMs (red branch). Plant MNS5s (violet branch) are also closely related to EDEM1s/EDEM3s. The lineage giving rise to EDEM2 (yellow branch) which due to its specificity for the outer mannose on the B branch of the glycan is no longer considered a *bona fide* ERAD checkpoint mannosidase [19, 44] appears to derive from a later duplication event than that giving rise to EDEM1/EDEM3/MNS5; plant MNS4s (green branch) cluster with this EDEM2 lineage.

### The *Ct*EDEM:PDI heterodimer functions as an ERAD checkpoint in plants and human cells

First, we set out to assay ERAD mannosidase activity of *Ct*EDEM. We did so *in planta*, using a complementation transfection assay in *Arabidopsis thaliana* (*At*). The assay readout relies on the dwarf plant phenotype associated with expression of a mutant of the Brassinosteroid-Insensitive 1 (BRI1) growth receptor: the *bri1-5* mutant retains residual growth receptor activity but it is degraded by the plant EDEMs, called MNS4 and 5, stunting the growth of the plant. When MNS4 and MNS5 are deleted in the *bri1-5* background (triple mutant *At mns4-1 mns5-1 bri1-5*) the *bri1-5* receptor is not degraded, its secretion is restored and the associated residual activity is sufficient to rescue the WT growth phenotype [51, 52]. Co-localisation of *Ct*EDEM:mRFP construct with the ER marker GFP:KDEL confirmed its localisation to the plant ER (Supplementary Figures S4A-K). *Ct*EDEM transfection of the triple mutant *At mns4-1 mns5-1 bri1-5* resulted in functional re-complementation of EDEM, commitment of *bri1-5* to ERAD and restoration of the dwarf phenotype (Supplementary Figures S4L,M).

We next investigated whether *Ct*EDEM can target canonical ERAD substrates for degradation, such as the A1AT-NHK variant of *α*1-antitrypsin (A1AT-NHK) and soluble tyrosinase (ST-TYR). Both A1AT-NHK and ST-TYR have been previously shown to be substrates of human EDEM1 [54, 139] and EDEM3 [16, 30]. For this purpose, HEK293T EDEM3-KO cells were co-transfected with *Ct*EDEM, *Ct*PDI, or their combination, together with either A1AT-NHK or ST-TYR, and the cell lysates analysed by Western Blotting to follow degradation of the ERAD client. Following signal peptide cleavage in the ER, A1AT retains a single cysteine residue (Cys 256) that can form intermolecular disulfide bonds either with ERAD checkpoint complexes or with itself, leading to dimerization and prolonged ER retention [54, 140]. Overexpression of *Ct*PDI in human cells, in the absence of *Ct*EDEM, shifted the A1AT-NHK dimer:monomer equilibrium toward the monomer (Figure 1A, left and right upper panels). By contrast, expression of *Ct*EDEM – alone or together with *Ct*PDI—enhanced degradation of A1AT-NHK – reduced both monomeric and dimeric protein levels (Figure 1A, left and right upper panels). Cycloheximide chase experiments further confirmed that A1AT-NHK is an ERAD substrate for *Ct*EDEM and the *Ct*EDEM:PDI complex (Figure 1D, upper and lower panels). Notably, co-expression of *Ct*PDI increased *Ct*EDEM mannosidase activity, as evidenced by bands of higher electrophoretic mobility corresponding to mannose removal (marked with asterisks in Figures 1A and D).

A similar effect was observed for ST-TYR, where *Ct*EDEM expression—alone or with *Ct*PDI enhanced degradation and reduced protein half-life (Figures 1B and E, upper and lower panels). Again, *Ct*PDI co-expression increased *Ct*EDEM mannosidase activity, as indicated by the faster-migrating, mannose-trimmed species (asterisk-marked bands).

To assess whether mannosidase activity is essential for *Ct*EDEM-mediated degradation, we examined a misfolded tyrosinase mutant lacking its six N-glycosylation sites (non-glycosylated tyrosinase, NG-TYR). Human EDEM2 selectively promotes degradation of the glycosylated soluble tyrosinase ERAD substrate ST-TYR, and this process is sensitive to kifunensine and proteasome inhibition; but not non-glycosylated NG-TYR [141]. EDEM3 similarly promotes processing and degradation of glycosylated ST-TYR through its mannosidase-like/GH47 domain [30]. Co-expression of *Ct*EDEM, either alone or with *Ct*PDI, did not significantly affect NG-TYR degradation (Figure 1C, upper and lower panels), confirming that *N*-glycosylation is required for *Ct*EDEM-dependent degradation of this substrate.

**Figure S18.**
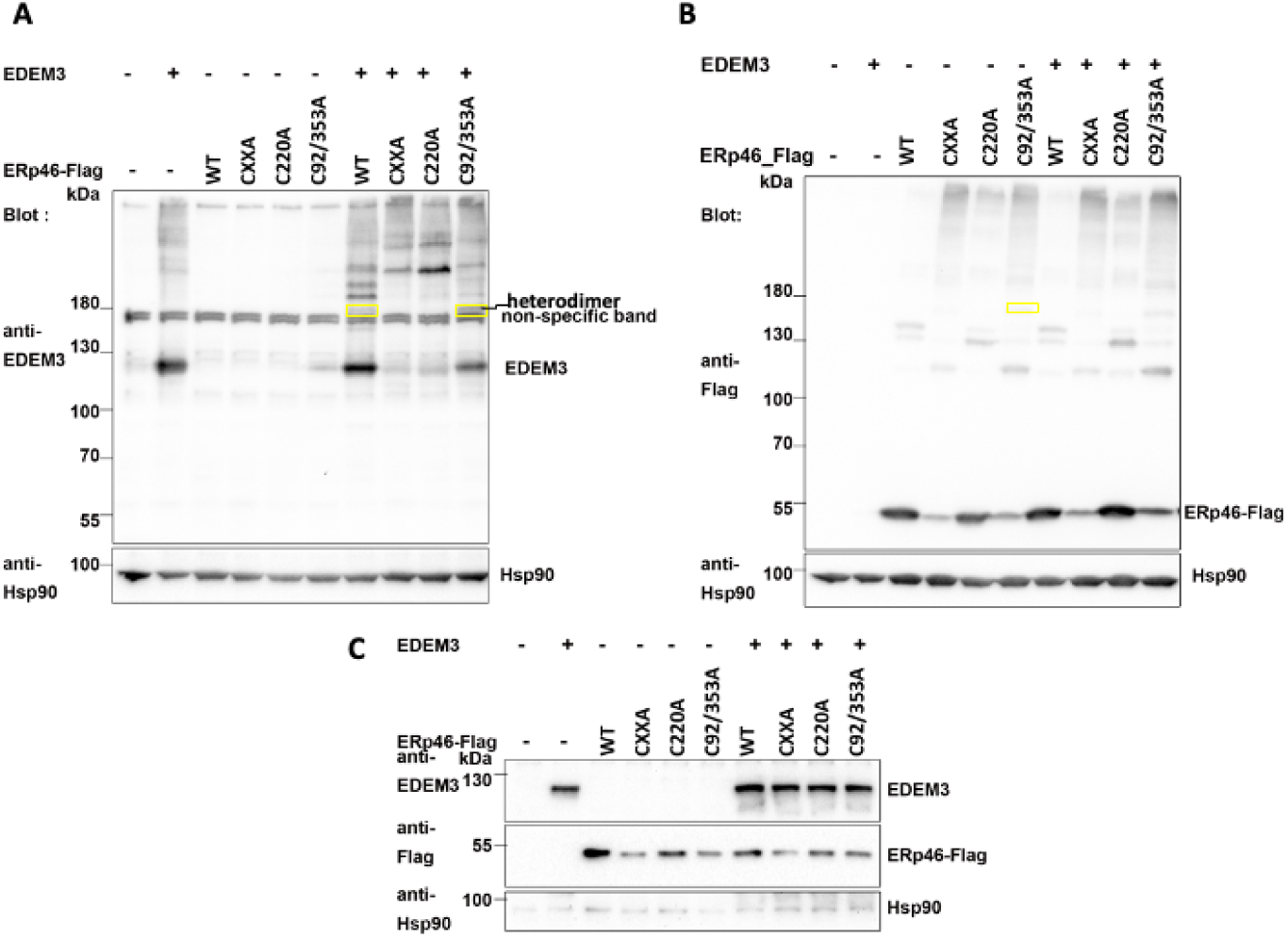
The C220A *Hs*ERp46 trapping mutant causes the disappearance of the HsEDEM3-SS-*Hs*ERp46 heterodimer band. EDEM3 KO HEK293T cells co-transfected with HsEDEM3 and *Hs*ERp46 WT and trapping mutants, were analysed by Western Blot in non-reducing (**A,B**) and reducing conditions (**C**). **A**: Detection of HsEDEM3 and its covalently associated complexes, separated on a 6% non-reducing polyacrylamide gel, transferred to a nitrocellulose membrane and detected with polyclonal anti-EDEM3 antibody; the yellow boxes indicate heterodimers and * a non-specific band. B: Detection of *Hs*ERp46 and its covalently associated complexes separated ona non-reducing 6% polyacrylamide gel, transferred on a nitrocellulose membrane and detected with monoclonal anti-FLAG HRP antibody. **C**: HsEDEM3 and *Hs*ERp46 WT and trapping mutants were reduced, separated on an 8% polyacrylamide gel, transferred to a nitrocellulose membrane and detected with the same antibodies as in A. and B. Hsp90 was used as loading control in all the experiments.

### *Ct*EDEM:PDI inter-domain conformational flexibility

As described in the previous section, the major difference in *Ct*EDEM upong going from the *Ct*EDEM:PDI:A1AT-NHK complex to the apo *Ct*EDEM:PDI structure is the swinging of the C-terminal domains from the GH47-distal to the GH47-proximal face of the complex, accompanied by a re-arrangement of the linear linker, and the oxidation of the intermolecular disulfide at Cys B.

3D variability analysis (3DVA) in cryoSPARC^™^ was utilised to investigate the flexibility of *Ct*EDEM:PDI portions built in volumes with lower local resolution. 3DVA analysis of the particles within the dataset suggested three specific motions. The first involves the IMD and PAD (Figure S19i) and the second and third involves the three-leaf clover PDI tail (Figure S19ii and S19iii). The first motion can be described as a twisting and pivoting of the IMD and PAD around the main axis of the particle and away from the active site face of the GH47 domain, while the three-leaf clover and the GH47 domain remain relatively unperturbed (Supplementary Videos 1 and 2). The second and third motions involves twisting of the TRX1-3 domains with respect to TRX4 and EDEM (Supplementary Video 4). Figure S20 shows EM maps obtained from particles sampling the endpoints of the third movement: the PDI three-leaf clover straightens (bends) with respect to the main body of the complex, so that the particle toggles between a linear (Figure S20A-C, yellow map) and a bent conformation (Figure S20A-C, blue map). Supplementary Video 3, shows the three-leaf clover undergoing an angular change of approximately 15°. The observed conformational flexibility is highly consistent with biophysical studies of PDIs [142, 143].

**Figure S19.**
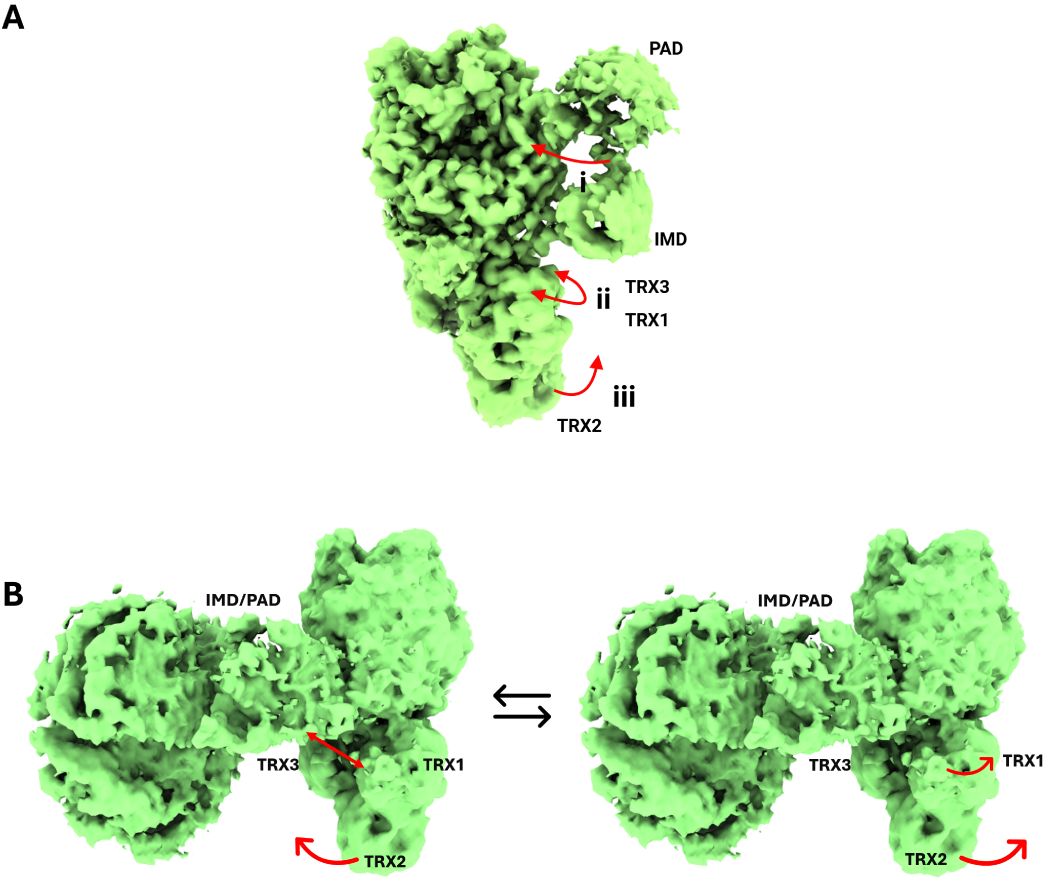
3D variability analysis of the *Ct*EDEM:PDI and PDI:EDEM:NHK complexes. Focussed 3D variability analysis in cryoSPARC^™^ of the *Ct*EDEM:PDI particles. **A:** apo model. Three principal ranges of motion: (i) front-to-back and up-and-down movements of the *Ct*EDEM IMD and PAD; (ii) twisting motion of the *Ct*PDI TRX1–3 domains; and (iii) flapping of the *Ct*PDI TRX2 domain. **B:** A1AT-NH bound model. The TRX domains move away from the A1AT-NHK model while the linker rearranges itself towards it and the intermolecular SS bond at Cys B is reduced.

**Figure S20.**
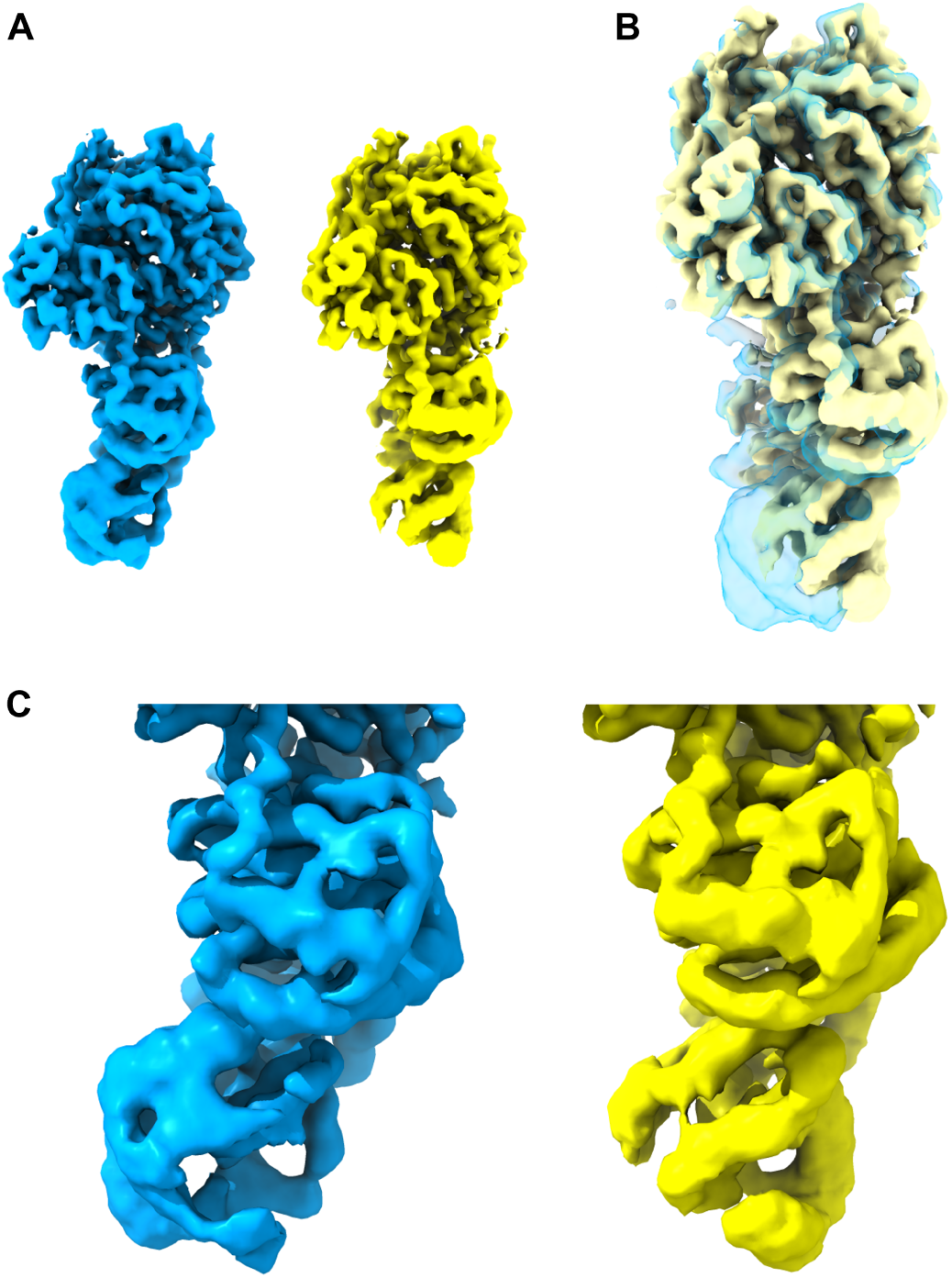
3D maps generated by 3D variability analysis of the apo *Ct*PDI:*Ct*EDEM particles. The surfaces illustrate the end conformations of the movements associated with principal components 1 (yellow) and 2 (blue) of the TRX2 swinging motion. **A.** Side by side comparison of the two extreme conformations. **B.** The two conformations are overlaid, with component 2 (blue) displayed at 70% transparency. **C.** Close-up view of the two conformations (TRX2 at the bottom). Figure made with ChimeraX [57].

**Figure S21.**
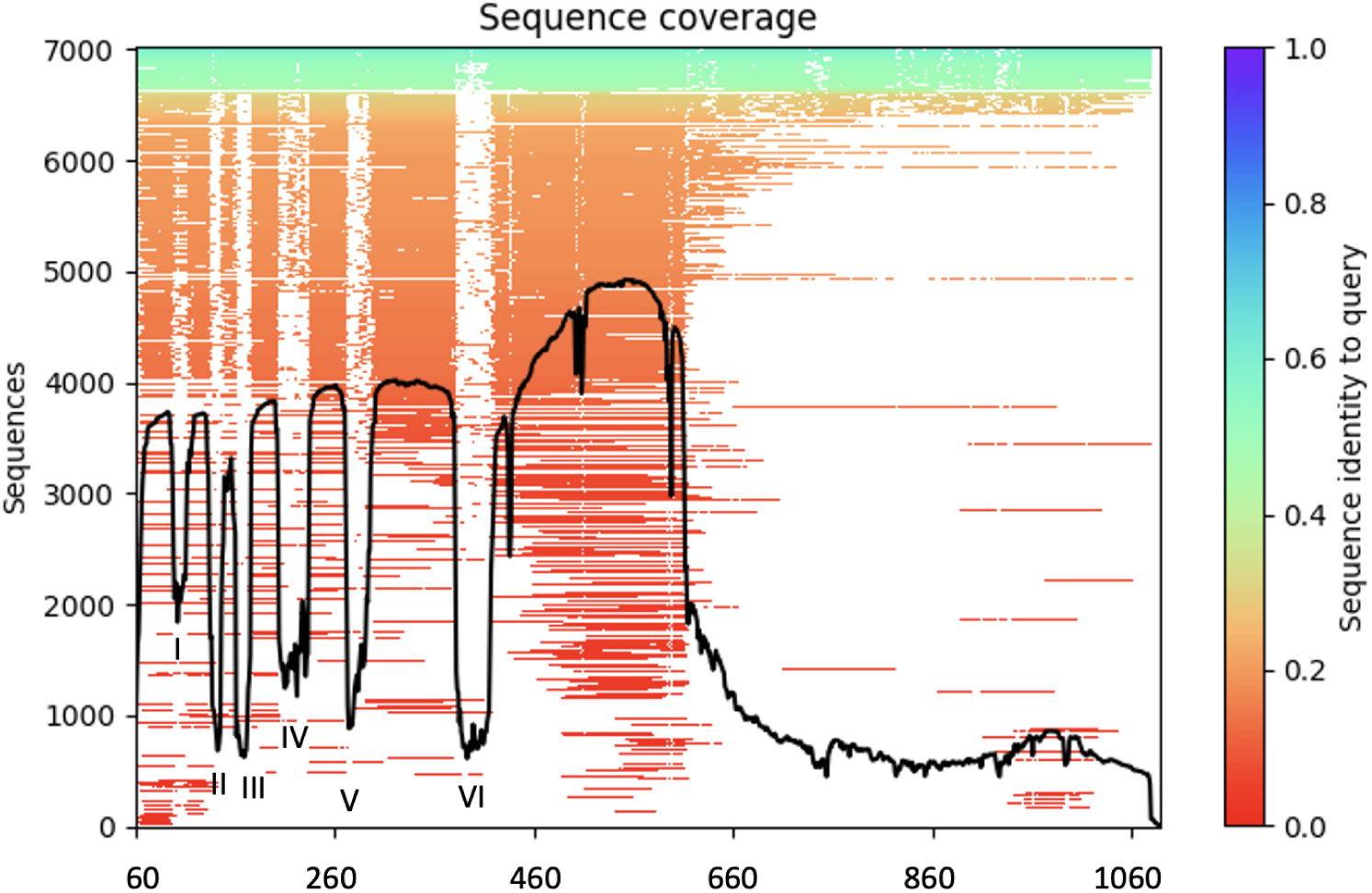
*Ct*EDEM sequence coverage as computed by AlphaFold3. About 600 sequences are highly homologous across the whole sequence (green-cyan) likely represent thermophilic EDEMs orthologous to *Ct*EDEM. Around 200 or so sequences are homologous to *Ct*EDEM in the GH47 domain and in the 100-120 substrate-binding region (labelled “I”): these are likely other eukaryotic EDEMs (yellow-orange). Some 2,200 GH47 mannosidases which are not EDEMs (dark orange) retain good homology in the GH47 domain but not in the five regions (“II” to “VI”) which are specific to fungal EDEMs.

**Figure S22.**
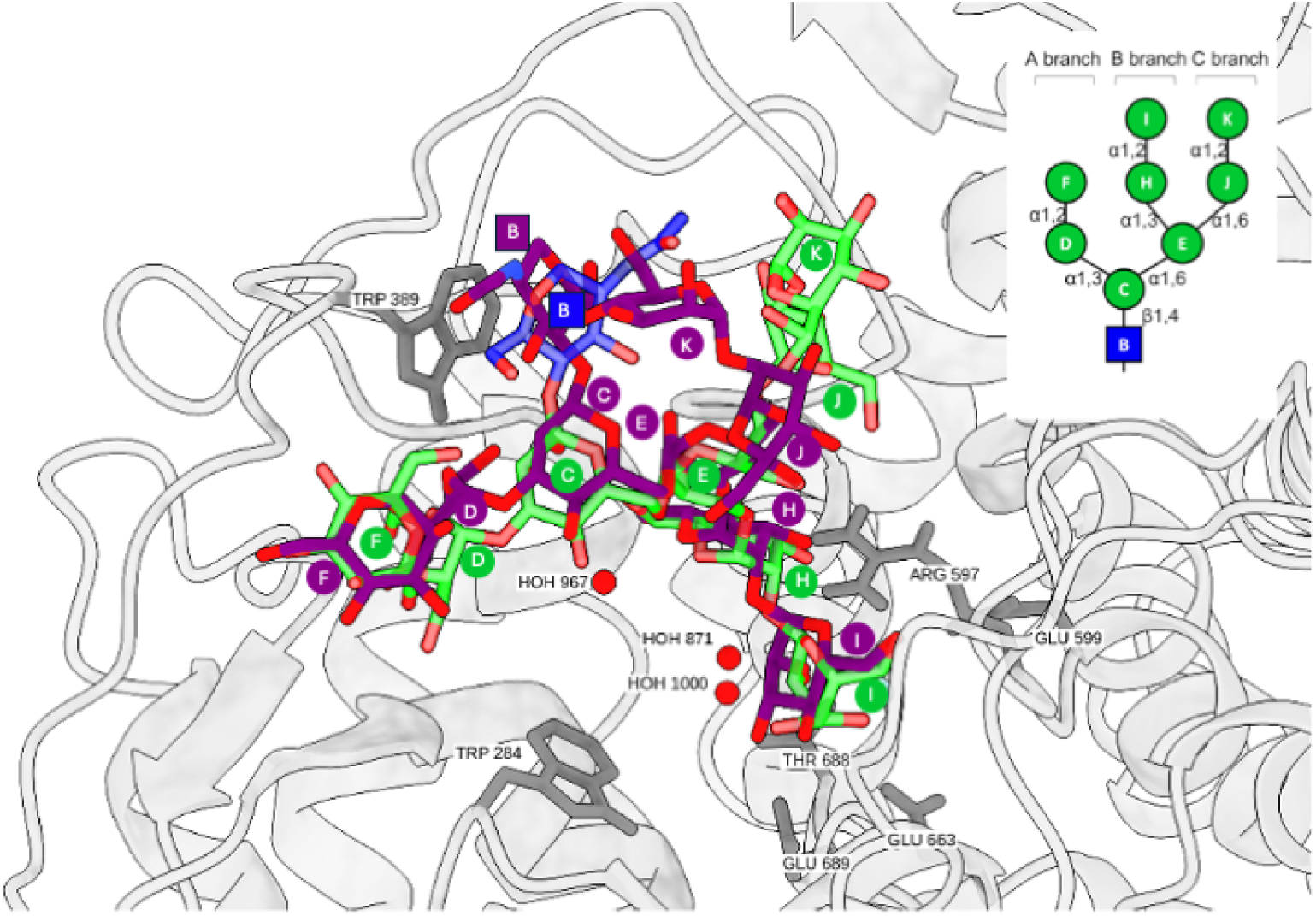
Re-docking of the GlcNAc_1_Man_8_ (M8A) *N*-glycan to the human ERManI active site. The crystal structure of the GlcNAc_1_Man_8_ (M8A) *N*-glycan (sticks, green/blue C atoms) bound t the ERManI active site (cartoon representation, silver) (PDB ID 5KIJ, [76]) superimposed with the docked model of the same *N*-glycan (sticks, purple C atoms) after optimisation of the parameters in the docking protocol. Individual sugar ring nomenclature as in the inset. Key residues stabilising the glycan are highlighted in grey stick representation. Three structural waters important for the catalytic mechanism [74, 123] are included (red spheres). Figure made with UCSF ChimeraX [57].

**Table S1.** Cross-species mapping of cysteine residues discussed in EDEM:PDI ERAD-L checkpoint complexes. *Ct*EDEM:*Ct*PDI is used as the structural reference system. Rows corresponding to conserved cysteine classes report the *C. thermophilum* residue together with the equivalent residues in *S. cerevisiae Sc*Mnl1:Pdi1 and mammalian EDEM:PDI complexes, as supported by structural comparison, sequence conservation or homology modelling. Additional species-specific cysteines discussed in the text, including residues with currently unassigned *Ct*EDEM:*Ct*PDI equivalence, are included as standalone entries. PDI cysteines are grouped according to the redox motif in which they occur. A1AT-NHK Cys 256 is included as the client cysteine associated with disulfide-linked A1AT-NHK species.

| Cysteine | S–S with |
| --- | --- |
| *CtEDEM* Cys 99,<br>*ScMnl1* Cys 65, *HsEDEM1* Cys 160,<br>*HsEDEM2* Cys 65, *HsEDEM3* Cys 82,<br>*MmEDEM3* Cys 83 | Intramolecular, in GH47: *CtEDEM* Cys 558,<br>*ScMnl1* Cys 445, *HsEDEM1* Cys 529,<br>*HsEDEM2* Cys 408, *HsEDEM3* Cys 441,<br>*MmEDEM3* Cys 442 |
| *CtEDEM* Cys 963,<br>*ScMnl1* Cys 721,<br>*HsEDEM3* Cys 693,<br>*MmEDEM3* Cys 694 | Intramolecular, in PAD: *CtEDEM* Cys 985,<br>*ScMnl1* Cys 787,<br>*HsEDEM3* Cys 714,<br>*MmEDEM3* Cys 715 |
| *CtEDEM* Cys 647,<br>*ScMnl1* Cys 579, *HsEDEM1* Cys 629<br>*HsEDEM2* Cys 558, *HsEDEM3* Cys 528<br>*MmEDEM3* Cys 529 | Intermolecular SS1: *CtPDI* Cys 385,<br>*ScPdi1* Cys 406, *HsPDIA1* Cys 397,<br>*HsTXNDC11* Cys 719, *HsERp46* Cys 217,<br>*MmERp46* Cys 203 |
| *CtEDEM* Cys 719,<br>*ScMnl1* Cys 644, *HsEDEM3* Cys 557,<br>*MmEDEM3* Cys 558 | Intermolecular SS2: *CtPDI* Cys 50 in apo<br>(free in A1AT-NHK -bound state),<br>*ScPdi1* Cys 61, *HsERp46* Cys 89,<br>*MmERp46* Cys 75 |
| *CtPDI* Cys 53,<br>*ScPdi1* Cys 64,<br>*HsERp46* Cys 92,<br>*MmERp46* Cys 78 | *CtPDI* Cys 50,<br>*ScPdi1* Cys 61,<br>*HsERp46* Cys 89,<br>*MmERp46* Cys 75<br>during redox cycling of the CXXC motif |
| *CtPDI* Cys 388,<br>*ScPdi1* Cys 409,<br>*HsPDIA1* Cys 400,<br>*HsERp46* Cys 220,<br>*MmERp46* Cys 206 | *CtPDI* Cys 385,<br>*ScPdi1* Cys 406,<br>*HsPDIA1* Cys 397,<br>*HsERp46* Cys 217,<br>*MmERp46* Cys 203<br>during redox cycling of the CXXC motif |
| A1AT-NHK Cys 256 | Itself in A1AT-NHK dimerisation<br>or a client's free Cys in a PDI mixed-disulfide species |

**Table S2.** AlphaFold3 statistics [128, 144] of models of *At*MNS5 in complex with *Arabidopsis thaliana* PDIs [53]. Entry IDs of UniProt and The Arabidopsis Information Resource (TAIR) identify the PDI [53]. Although the models omitted the signal peptide, residue numbering refers to the full length protein, not the mature form.

| PDI,<br>UniProt ID<br>TAIR ID | Name | Short<br>Names | ipTM | PTM | *AtMNS*-SS-*AtPDI* |
| --- | --- | --- | --- | --- | --- |
| *AtPDIL2*-1,<br>O22263,<br>At2g47470 | Protein disulfide<br>isomerase-like 2-1 | MEE30,<br>PDI11,<br>PDIL4-1,<br>UNE5 | 0.83 | 0.75 | Cys515-S-S-Cys52 |
| *AtPDIL1*-1,<br>Q9XI01<br>At1g21750 | Protein disulfide<br>isomerase-like 1-1 | PDI5 | 0.81 | 0.73 | Cys515-S-S-Cys404 |
| *AtPDIL1*-4,<br>F4K0F5,<br>F4K0F7 | Protein disulfide<br>isomerase-like 1-4 | PDI2,<br>MUP24.6 | 0.83 | 0.65 | Cys515-S-S-Cys407 |
| *AtPDIL1*-3,<br>Q8VX13,<br>At3g54960 | Protein disulfide<br>isomerase-like 1-3 | PDI1,<br>PDIL2-1,<br>PDI72 | 0.80 | 0.66 | Cys515-S-S-Cys467 |
| *AtPDIL1*-4,<br>Q9FF55,<br>At5g60640 | Protein disulfide<br>isomerase-like 2-2 | PDI2,<br>MUP24.6 | 0.78 | 0.62 | Cys515-S-S-Cys496 |
| *AtPDIL1*-2,<br>Q9SRG3<br>At1g77510 | Protein disulfide<br>isomerase-like 1-2 | PDI6 | 0.75 | 0.67 | Cys515-S-S-Cys401 |
| *AtPDIL5*-1,<br>Q8GYD1,<br>At1g07960 | Protein disulfide<br>isomerase-like 5-1 | PDIL6-1 | 0.61 | 0.68 | Cys515-S-S-Cys55 |
| *AtPDIL1*-6,<br>Q66GQ3,<br>At3g16110 | Protein disulfide<br>isomerase-like 1-6 | PDI4,<br>PDIL3-2 | 0.47 | 0.60 | – |
| *AtPDIL5*-3,<br>Q9LJU2,<br>At3g20560 | Protein disulfide<br>isomerase-like 5-3 | PDI12,<br>PDIL8-1 | 0.44 | 0.56 | Cys515-S-S-Cys170 |
| *AtPDIL2*-3,<br>O48773 | Protein disulfide<br>isomerase-like 2-3 | PDI9,<br>PDIL5-2 | 0.26 | 0.51 | Cys515-S-S-Cys192 |
| *AtPDIL2*-1,<br>F4IL53,<br>At2g47470 | Protein disulfide<br>isomerase-like 2-1 | PDI11,<br>UNE5 | 0.22 | 0.61 | Cys515-S-S-Cys74 |
| *AtPDIL5*-2,<br>Q94F09,<br>At1g35620 | Protein disulfide<br>isomerase-like 5-2 | PDI8,<br>PDIL7-1 | 0.21 | 0.54 | – |
| *AtPDIL2*-2,<br>Q9MAU6,<br>At1g04980 | Protein disulfide<br>isomerase-like 2-2 | PDI10,<br>PDIL5-1 | 0.21 | 0.52 | – |
| *AtPDIL1*-5,<br>A3KPF5,<br>At1g52260 | Protein disulfide<br>isomerase-like 1-5 | PDI3,<br>PDIL3-1 | 0.18 | 0.50 | – |

**Table S3.** AlphaFold3 statistics [128, 144] of models of *At*MNS4 in complex with *Arabidopsis thaliana* PDIs. Entry IDs of UniProt and The Arabidopsis Information Resource (TAIR) identify the PDI [53]. Although the models omitted the signal peptide, residue numbering refers to the full length protein not the mature form.

| PDI,<br>UniProt ID<br>TAIR ID | Name | Short<br>Names | ipTM | PTM | *AtMNS*-SS-*AtPDI* |
| --- | --- | --- | --- | --- | --- |
| *AtPDIL2-2*,<br>Q9MAU6,<br>At1g04980 | Protein disulfide<br>isomerase-like 2-2 | PDI10,<br>PDIL5-1 | 0.55 | 0.58 | – |
| *AtPDIL1-2*,<br>Q9SRG3 | Protein disulfide<br>isomerase-like 1-2 | PDI6 | 0.35 | 0.50 | Cys532-S-S-Cys401 |
| *AtPDIL1-3*,<br>Q8VX13,<br>At3g54960 | Protein disulfide<br>isomerase-like 1-3 | PDI1,<br>PDIL2-1,<br>PDI72 | 0.3 | 0.49 | Cys532-S-S-Cys442 |
| *AtPDIL2-3*,<br>O48773,<br>At2g32920 | Protein disulfide<br>isomerase-like 2-3 | PDI9,<br>PDIL5-2 | 0.26 | 0.51 | – |
| *AtPDIL1-1*,<br>Q9XI01 | Protein disulfide<br>isomerase-like 1-1 | PDI5 | 0.32 | 0.51 | Cys532-S-S-Cys404 |
| *AtPDIL1-4*,<br>Q9FF55,<br>At5g60640 | Protein disulfide<br>isomerase-like 2-2 | PDI2,<br>MUP24.6 | 0.25 | 0.48 | Cys532-S-S-Cys446 |
| *AtPDIL5-2*,<br>Q94F09,<br>At1g35620 | Protein disulfide<br>isomerase-like 5-2 | PDI8,<br>PDIL7-1 | 0.24 | 0.53 | Cys532-S-S-Cys38 |
| *AtPDIL1-6*,<br>Q66GQ3,<br>At3g16110 | Protein disulfide<br>isomerase-like 1-6 | PDI4,<br>PDIL3-2 | 0.23 | 0.49 | Cys532-S-S-Cys74 |
| *AtPDIL1-4*,<br>F4K0F5,<br>F4K0F7 | Protein disulfide<br>isomerase-like 1-4 | PDI2,<br>MUP24.6 | 0.21 | 0.48 | Cys532-S-S-Cys |
| *AtPDIL1-5*,<br>A3KPF5,<br>At1g52260 | Protein disulfide<br>isomerase-like 1-5 | PDI3,<br>PDIL3-1 | 0.21 | 0.48 |  |
| *AtPDIL2-1*,<br>F4IL53,<br>At2g47470 | Protein disulfide<br>isomerase-like 2-1 | PDI11,<br>UNE5 | 0.17 | 0.58 | – |
| *AtPDIL2-1*,<br>O22263,<br>At2g47470 | Protein disulfide<br>isomerase-like 2-1 | MEE30,<br>PDI11,<br>PDIL4-1,<br>UNE5 | 0.17 | 0.53 | Cys532-S-S-Cys382 |
| *AtPDIL5-1*,<br>Q8GYD1,<br>At1g07960 | Protein disulfide<br>isomerase-like 5-1 | PDIL6-1 | 0.16 | 0.63 | – |
| *AtPDIL5-3*,<br>Q9LJU2,<br>At3g20560 | Protein disulfide<br>isomerase-like 5-3 | PDI12,<br>PDIL8-1 | 0.14 | 0.46 | Cys532-S-S-Cys170 |

**Table S4.**
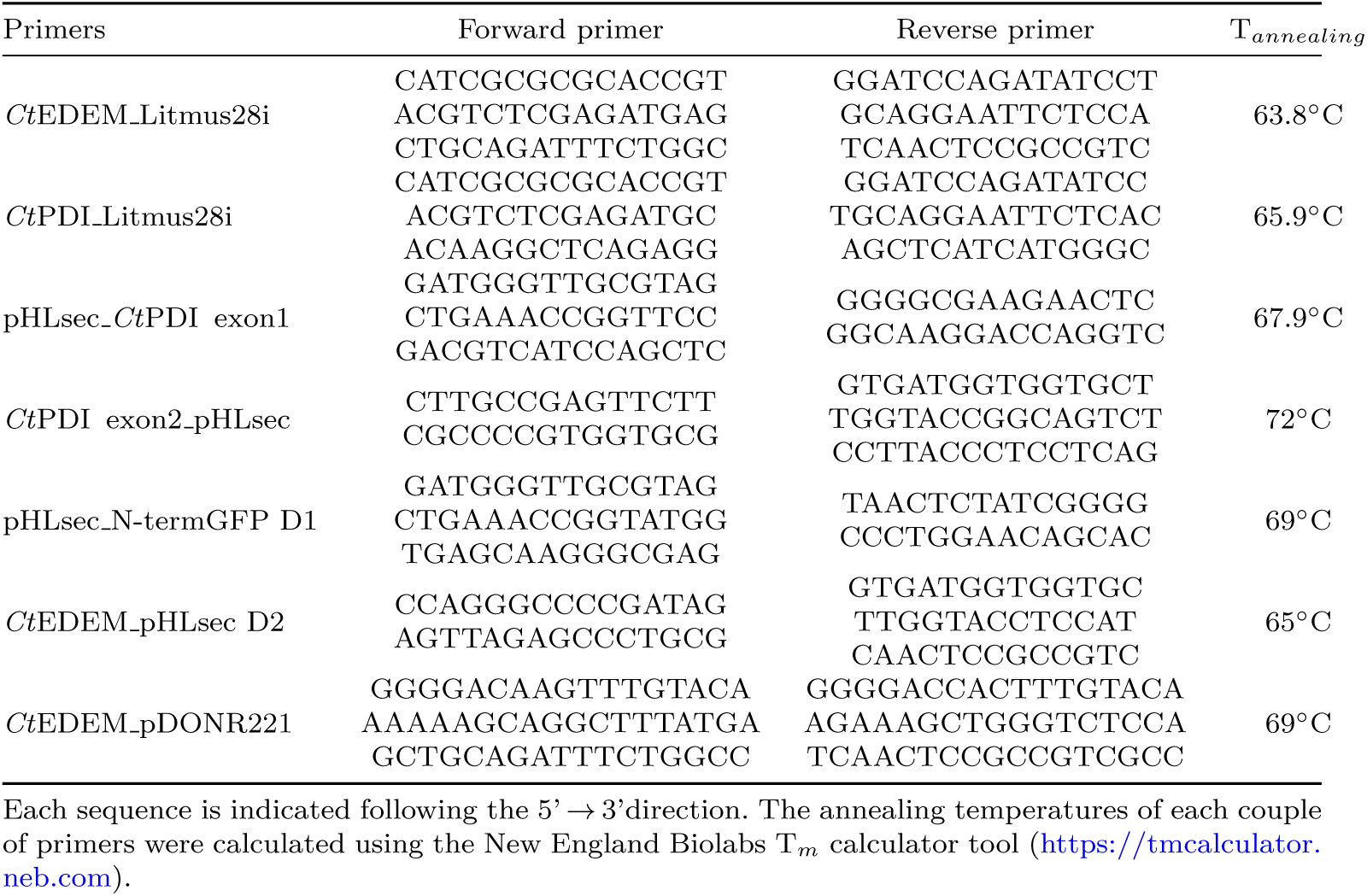
List of primers used for PCR cloning.

## Footnotes

1 The reported EDEM1 association with ERdj5 (*aka* DNAJC10) [37–39] is likely not direct, but mediated by the ERdj5 co-chaperone BIP [33, 40]. ERdj5 associates with EDEM1 without stimulating its mannosidase activity, suggesting a role in downstream ERAD events rather than in glycan trimming [40, 41].

2 Unlike EDEM1 and EDEM3, EDEM2 removes the outer Man residue specifically from branch B of the *N*-linked glycan [19, 43, 44]

3 The MNS4 and MNS5 partner PDIs are not known. Supplementary Tables S2 and S3 list quality statistics for AlphaFold3-generated models of complexes of *At*MNS5 and *At*MNS4 with a number of representative *At*PDI family members [53].

4 Between the deposition of the apo structure and its release, *Ct*EDEM:PDI was contributed to the list of CASP-15 protein prediction targets (CASP-15 Target H1157, https://predictioncenter.org/casp15/target.cgi?id=116&view=all): a brief description of how the predictors did on this target was published in [59].

5 Homology modelling and biochemical data also suggest equivalent intermolecular disulfide bridges in-volving Cys A and one of the redox-active CXXC motifs on the PDI in the three mammalian EDEM:PDI complexes [19, 45, 64]. The corresponding EDEM and PDI cysteine residues are listed in Table S1, and the structural models are shown in Supplementary Figure S17.

6 We cannot exclude the possibility that the *Mm*EDEM3:*Hs*ERp46 Cys220A mutant migrates as a higher molecular mass band of ∼ 250 kDa on the non-reducing gel because of a different inter-molecular disulfide link between subunits [88].

